# Glycodiversification of gentamicins through *in vivo* glycosyltransferase swapping enabled the creation of novel hybrid aminoglycoside antibiotics with potent activity and low ototoxicity

**DOI:** 10.1101/2023.11.29.569180

**Authors:** Xinyun Jian, Cheng Wang, Shijuan Wu, Guo Sun, Chuan Huang, Chengbing Qiu, Yuanzheng Liu, Peter F. Leadlay, Dong Liu, Zixin Deng, Fuling Zhou, Yuhui Sun

**Author notes:** These authors contributed equally (Xinyun Jian, Cheng Wang).

## Abstract

Aminoglycosides (AGs) are a class of potent antibiotics with a broad spectrum of activity. However, their use is limited by safety concerns associated with nephrotoxicity and ototoxicity, as well as drug resistance. To address these issues, semi-synthetic approaches for modifying natural AGs have successfully generated new generations of AGs, however, with limited types of modification due to significant challenges in synthesis. This study explores a novel approach that harness the bacterial biosynthetic machinery of gentamicins and kanamycins to create hybrid AGs, installing extensive natural modifications from gentamicins onto kanamycins. This was achieved by glycodiversification of gentamicins via swapping the glycosyltransferase (GT) in their producer with the GT from kanamycins biosynthetic pathway and resulted in the creation of a series of novel AGs with combined structural features of two, therefore referred to as genkamicins (GKs). The manipulation of the hybrid metabolic pathway enabled the target accumulation of different GK species and the successful isolation and characterization of six GK components. These compounds display retained antimicrobial activity against a panel of World Health Organization (WHO) critical priority pathogens, and GK-C2a, in particular, demonstrates low ototoxicity compared to clinical drugs in zebrafish embryos. This study provides a new strategy for diversifying the structure of AGs and a potential avenue for developing less toxic AG drugs to combat infectious diseases.

## Introduction

Aminoglycosides (AGs), originally isolated from soil bacteria, *Streptomyces* and *Micromonospora*, are highly potent and broad-spectrum antibiotics that have been used to treat bacterial infections since the 1940s. AGs act as bacterial protein synthesis inhibitors by binding to 16S ribosomal RNA of the 30S ribosome, which can promote mistranslation, block elongation, or inhibit the initiation of protein synthesis. Structurally, AGs are characterized by a core aminocyclitol moiety (ring I) connected to amino sugars (ring II, III, and IV) via glycosidic linkages and, in most cases, forming a pseudotrisaccharide scaffold. Most of the clinically important AGs contain 2-deoxystreptamine (2-DOS) core as aminocyclitol moiety and can be classified into two groups depending on the position of substituted sugars: 4,5-disubstituted 2-DOS containing AGs, represented by neomycins and burtirosins, and 4,6-disubstitued 2-DOS containing AGs, represented by gentamicins and kanamycins^1^ (Fig. 1). They can also be categorized into three subfamilies based on their distinct structures of pseudotrisaccharide scaffold, including kanamycin, gentamicin, and neomycin families, which feature modified glucose, xylose, and ribose as ring III, respectively (Fig. 1). Notably, the natural gentamicin family AGs have the most extensive modifications, including amination, *C*-methylation, *N*-methylation, didehydroxylation, dehydrogenation and epimerization compared to other family natural AGs.

**Fig. 1.**
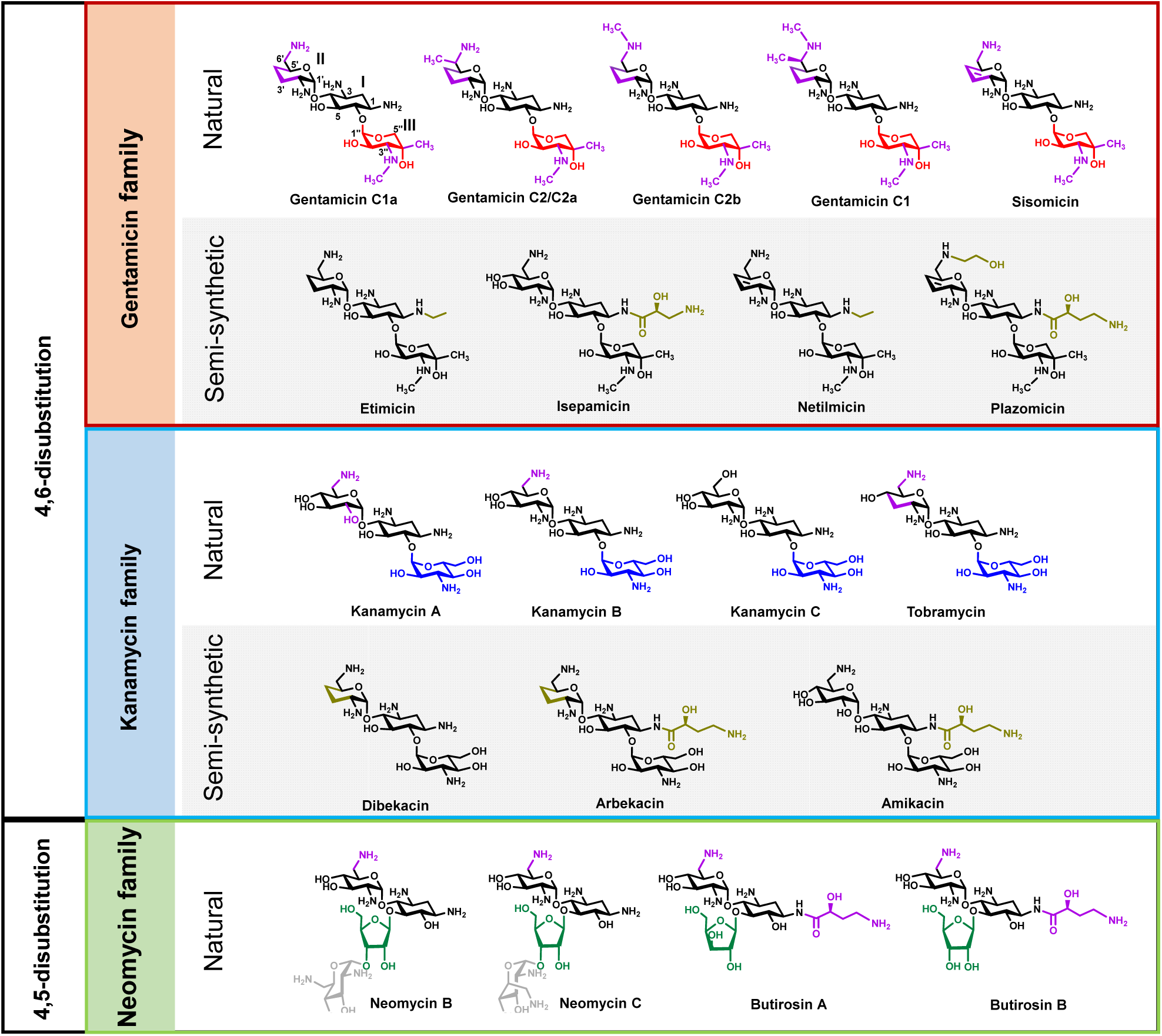
Structure and classification of natural and semi-synthetic 2-DOS containing AG drugs. Red, blue, and green highlighted moieties respectively represent xylose, glucose, and ribose as the different second sugar scaffold (ring III) in gentamicin, kanamycin, and neomycin family AGs. The additional fourth sugar in neomycin is highlighted in grey. Purple indicates modifications installed via natural biosynthetic process, while gold indicates modifications installed via semi-synthetic methods. Annotation to carbon and ring numbers is shown in gentamicin C1a.

Nephrotoxicity and ototoxicity have always been the safety concerns associated with the prolonged use of AGs. While nephrotoxicity is generally reversible due to the regeneration capability of tubular cells and diminished AG accumulation in the kidneys, ototoxicity, on the other hand, (in some cases) leads to permanent and irreversible hearing loss. The extensive clinical application is also the source of the severe bacterial resistance encountered today. AGs resistance can arise from various mechanisms, including enzymatic modification of the chemical structure of AGs, chromosomal mutation, or modification of the target site and efflux. Of these mechanisms deactivating the AGs through modifying amine and hydroxyl groups by the bacterial AG-modifying enzymes (AMEs), including the acetyl-CoA-dependent AG acetyltransferases (ACCs), the ATP (and/or GDP)-dependent AG phosphotransferases (APHs) and the ATP-dependent AG nucleotidyltransferases (ANTs), is the most prevalence mechanism underlying the widespread AG resistance^1^.

Despite the intrinsic toxicity and the ever-growing resistance threatening their long-term use, AGs remain a valuable component of the antibiotic armamentarium. Recent years have also seen revived interest in the new therapeutic potentials of this old class of antibiotics, such as treatment for fungal^2^ and viral infections^3^ and genetic diseases caused by premature termination codons (PTCs)^4,5^.

Substantial efforts have been made to develop chemical approaches for expanding the structural diversity of AGs to overcome the challenges associated with AGs resistances and toxicities. Direct modification of the natural AGs has successfully yielded a series of semisynthetic AG drugs, e.g., amikacin, dibekacin, and arbekacin as kanamycin derivatives; etilmicin and isepamicin as gentamicin derivatives; and netimicin and plazomicin as sisomicin derivatives^5^ (Fig. 1). However, the chemically regiospecific modification of densely functionalized and structurally diverse AGs commonly requires multi-step regiospecific protection/deprotection schemes. As a result, it has not only been a tedious synthetic endeavor for chemists, but it also commonly leads to poor yields of final products^6^.

Biotechnology offers an obvious appeal as a complementary strategy for efficient and regiospecific AG modifications, which has inspired researchers to harness the biosynthetic machinery of heavily modified natural AGs, like gentamicins and butirosins. The substrate tolerance of a few AGs biosynthetic enzymes has been exploited for the chemoenzymatic synthesis of non-natural AGs. For example, BtrH and BtrG, responsible for the addition of an (*S*)-4-amino-2-hydroxybutyric acid (AHBA) moiety to the butirosins^7^, were utilized to regiospecifically attach an AHBA side chain onto a range of natural AGs^8^; while GenN, one of the methyltransferases involved in gentamicins biosynthesis, has been shown to catalyze 3^″^-*N*-methylation in both kanamycin B and tobramycin^9^. GenN has also been coupled with the AAC(6’)-APH(2’’), the AME enzymes for C6’-amination from *Staphylococcus aureus*, to enzymatically synthesize amikacin analogous with improved pharmacological potential^10^.

Compared to chemical and chemoenzymatic synthesis, bioengineering approaches such as combinatory biosynthesis and biosynthetic pathway engineering can be more sustainable alternatives for efficiently modifying natural AGs, which, however, remain underexplored due to the limited understanding of their biosynthesis in the early years. The only reported attempt so far involved introducing seven butirosins biosynthetic genes (*btrI*-J-K-O-V-G-H), responsible for biosynthesis and incorporation of the AHBA moiety, to a kanamycins mini-biosynthetic gene cluster containing the homologous host. This resulted in the production of 1-*N*-AHBA-kanamycin X and amikacin but with meager yield (0.6 mg/L and 0.5 mg/L, respectively), limiting their further development (Park et al., 2011)^11^.

All the 2-DOS containing AGs biosynthesis starts with converting 6-P-Glc to the 2-DOS core by a set of conserved enzymes^12^. In the gentamicin biosynthesis, this is followed by the transfer of UDP-*N*-acetylglucosamine (UDP-GlcNAc) onto the C-4 position of 2-DOS by GenM1, the first glycosyltransferase (GT) in the gentamicin gene cluster, resulting in pseudodisaccharide scaffold paromamine, via deacylation of acetylparamomine by GenD (Fig. 2). Paromamine is then glycosylated at the C-6 position with UDP-xylose by the second GT, GenM2, to give the first pseudotrisaccharide scaffold, gentamicin A2^13^. In the last decade, significant progress has been made in characterizing the abundant subsequent modification steps. Notably, all these modifications are shown to occur on the pseudotrisaccharide scaffold after the formation of gentamicin A2. Gentamicin A2 first undergoes sequential C-3’’ amination and methylation by dehydrogenase GenD2, aminotransferase GenS2, and *N*-methyltransferase GenN to generate gentamicin A, which is further methylated at C-4’’ position by C-methyltransferase GenD1 to give gentamicin X2, the branch point pseudotrisaccharide intermediate leading to two major parallel pathways^14^. C6’-methylation of gentamicin X2 by C-methyltransferase GenK forms G418, which together with gentamicin X2 are subsequently aminated at C-6’ position by dehydrogenase GenQ and aminotransferase GenB1 in parallel to yield JI-20Ba and JI-20A, respectively^15,16^. The following C3’-C4’ didehydroxylation by phosphotransferase GenP and two aminotransferases GenB3 and GenB4 in sequence converts JI-20A and JI-20Ba to two of the gentamicin C complex, gentamicin C1a and C2a, respectively^17^. In addition, the last aminotransferase homolog GenB2 epimerize gentamicin C2a at C-6’ position to produce gentamicin C2^18,19^. Finally, the *N*-methyltransferase GenL catalyzes the 6’-*N*-methylation of both gentamicin C1a and C2 to afford gentamicin C2b and C1, respectively^15^. It is worth noting that recent research has revealed the complex modification system of gentamicins to possess broader substrate tolerance than previously believed, leading to the discovery of the multiple minor pseudotrisaccharide parallel modification pathways^15,20^ (Supplementary Fig. 1).

**Fig. 2.**
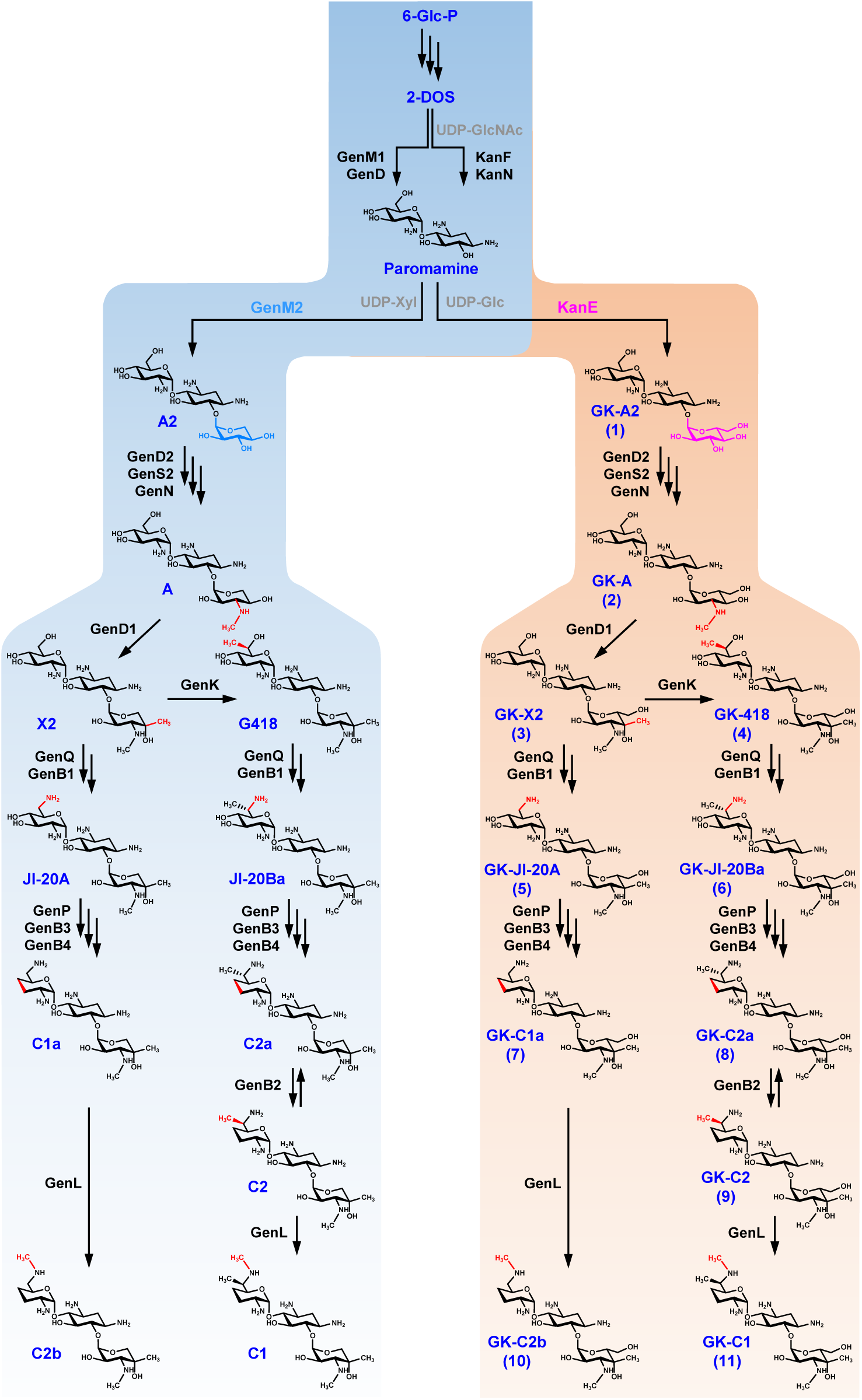
Biosynthetic pathways of gentamicins and GKs. The biosynthetic pathway of GKs is proposed based on the hypothesis that it follows the same modification orders of gentamicins biosynthetic pathway. The different sugar scaffolds incorporated in gentamicins and GKs are highlighted in blue (xylose) and pink (glucose), respectively. The structural changes resulting from each modification are indicated in red. 6-Glc-P, glucose 6-phosphate; 2-DOS, 2-deoxystreptamine; UDP, uridine diphosphate; UDP-GlcNAc, UDP-*N*-acetylglucosamine; UDP-Xyl, UDP-xylose; UDP-Glc, UDP-glucose.

Compared to gentamicins biosynthetic pathway that diverges in two main branches to incorporate all the modifications, the one of kanamycins has been demonstrated *in vitro* to split at two distinct points, formation of pseudodisaccharides by the first GT, KanF, and formation of the pseudotrisaccharides by the second GT, KanE, leading to 8 parallel pathways^11^ (Supplementary Fig. 1). KanF utilizes UDP-GlcNAc and UDP-Glc as different sugar donors to convert 2-DOS to paromamine and 2’-deamino-2’-hydroxyparomamine, respectively, which are then aminated at C-6’ by KanI-KanL to give another two pseudodisaccharides, neamine and 2’-deamino-2’-hydroxyneamine. KanE was shown to accept all four pseudodisaccharides as acceptor, as well as two different sugars, UDP-Glc and UDP-konasomine (UDP-Kns, converted from UDP-Glc by KanC-KanD), as donor. Nonetheless, the gentamicins and kanamycins biosynthesis share a biosynthetic route from Glc-6-P to their common pseudodisaccharide, paromamine.

The comprehensive understanding of the biosynthesis of these two 4,6-disustitutted-2-DOS-containing AGs provides the possibility for combinatory biosynthetic engineering that harnesses the robust modification system from gentamicin biosynthesis to modify kanamycin family scaffold and results in novel hybrid AGs. In principle, this strategy can be achieved by either introducing all the gentamicins modification enzymes into the kanamycins producer or introducing kanamycin pseudotrisaccharide scaffold biosynthetic machinery into the gentamicins producer. While the former poses a challenge to the successful expression of twelve genes in a heterologous host, we proposed that the latter can be achieved by glycodiversification of gentamicins by simply swapping the second GT GenM2 with its counterpart KanE. This was inspired by the fact that GenM2 and KanE share a common pseudodisaccharide, paromamine, as the sugar acceptor (Supplementary Fig. 1), and that Glc-6-P as a sugar donor of KanE can be derived from bacterial primary metabolism.

Herein, we report creating a novel class of AGs, called genkamicins (GKs), as a hybrid of gentamicins and kanamycins via combinatory biosynthesis and pathway engineering strategy. We began with assessing the feasibility of this approach by examining the capability of the gentamicins-producing strain *Micromonospora echinospora* ATCC15835 to modify exotic kanamycin B through a feeding experiment. Then, we employed the GT swapping approach within the gentamicin producer to replace the second GT, GenM2, with its counterpart KanE from kanamycin biosynthesis effectively producing a series of proposed GKs. Based on our understanding of gentamicin biosynthesis, we also achieved the targeted over-accumulation of the different GK products through metabolic engineering to facilitate the isolation of mono GK components. Finally, six GK products were successfully isolated, structurally characterized, and examined for antimicrobial activity and toxicity, revealing a less-toxic but still potent novel aminoglycoside GK-C2 (**9**).

## Results

### Gentamicin-producing strain is capable of modifying kanamycin B

Mimicking the biosynthetic logic of gentamicins that modifications primarily occur after the pseudotrisaccharide scaffold formation, we proposed a combinatory biosynthetic pathway in *M. echinospora* ATCC15835 to produce the hybrid AGs, genkamicins (GKs) (Fig. 2). In this pathway KanE is predicted to replace GenM2 to generate 3’’-dehydroxykanamycin C (GK-A2, **1**), the first pseudotrisaccharide equivalent to gentamicin A2 in gentamicins pathway, which is followed by a series of modifications to give the key intermediates GK-A (**2**), GK-X2 (**3**), GK-418 (**4**), GK-JI20A (**5**), GK-JI20Ba (**6**), and the five C complex components GK-C1a (**7**), GK-C2a (**8**), GK-C2 (**9**), GK-C2b (**10**), and GK-C1 (**11**), through two main parallel pathways.

This combinatory biosynthesis approach relies on two prerequisites: 1) the modification enzymes in gentamicin biosynthesis accept the kanamycin family trisaccharide scaffold as substrate, and 2) the gentamicin-producing strain can tolerate the novel AG products. As proof of concept, we probed the two prerequisites by the feeding experiment. Kanamycins is a mixture of three main mono components, kanamycin A, B, and C, with composition percentages varying from different vendors. Only two mono components, kanamycin A and B, are commercially available, and kanamycin B is structurally closer to GK-A2 (**1**) compared to that of kanamycin A. Thus, kanamycin B was fed to gentamicin-producing strain *M. echinospora* ATCC15835. Notably, four novel products with [M+H]^+^ ions corresponding to *m/z* 498. 2763 (calcd. for C_19_H_40_N_5_O_10_^+^: 498.2770), 512.2927 (calcd. for C_20_H_42_N_5_O_10_^+,^ 512.2926), *m/z* 480.3026 (calcd. for C_20_H_42_N_5_O_8_^+,^ 480.3028) and *m/z* 494.3178 (calcd. for C_21_H_44_N_5_O_8_^+^ 494.3184), respectively, were observed in the LC-ESI-HRMS analysis of culture extract, albeit at low levels even with increased feeding concentration (Supplementary Fig. 2). The calculated molecular formula and tandem MS/MS fragmentation pattern of these products are consistent with the ones of 3’’-*N*-methyl or 4’’-methyl-kanamycin B, GK-JI-20A (**5**), GK-C1a (**7**), and GK-C2 / C2a / C2b (**8** / **9** / **10**), respectively. However, due to the low yield, isolation and further structural elucidation of the novel products were impossible. To mitigate the potential competition from the native substrates for the modification enzymes, which may account for the low yield of GK products, we constructed the *genM2 in*-*frame* deletion mutant (Supplementary Fig. 3), ΔgenM2, that abolished all the native pseudotrisaccharides production (Supplementary Fig. 3b). However, feeding kanamycin B to ΔgenM2 did not significantly increase the production level of the new species, implying the modest yield of the GK products may stem from other factors, such as the restricted uptake of external kanamycin B by the producer.

Nonetheless, the observation of the new compounds demonstrated that the gentamicin-producing strain can tolerate the new products. It also suggested that modification enzymes involved in the two methylations on ring III (GenN and GenD1, respectively), one methylation (GenK or GenL), and didehydroxylation on ring II (GenP-GenB3-GenB4 cascade) are promiscuous and can accept kanamycin pseudotrisaccharide scaffold as substrate. Given that kanamycin B already bears amino groups at C-6 and C-3’’, the tolerance of enzymes for the amination of kanamycin pseudotrisaccharide scaffold at the corresponding positions (GenQ-GenB1 cascade and GenD2-GenS2 cascade, respectively) were not able to be predicted by the feeding experiment. However, on the other hand, our previous studies on GenD2-GenS2 cascade revealed that they were able to catalyze the reverse reaction, C-3’’-deamination of kanamycin B^14^, indicating they are likely to accommodate GK-A2 as a substrate. Taken together, these results validated the viability of the proposed combinatory biosynthesis strategy. The low yield of the novel GK products, on the other hand suggested an obvious abuse existed in generating novel AGs by feeding natural substrates.

### Producing a series of novel aminoglycosides GKs via glycosyltransferase swapping

We then cloned *kanE* onto an integrative vector, pWHU77, which carries a constitutive promoter P*ermE*\* and was previously used for gene complementation in the gentamicin producer to generate the *kanE* expression construct pWHU155. Swapping *genM2* with *kanE* was then achieved by introducing pWHU155 into *M. echinospora* ATCC15835 ΔgenM2 mutant. Remarkably, a series of new products with [M+H]^+^ ions and MS/MS fragmentation pattern corresponding to all the proposed GK products (Fig. 2) were successfully observed in the culture extract of ΔgenM2::kanE by HPLC-ESI-HRMS/MS (Fig. 3c and Supplementary Fig. 4). The number and yield of new products were significantly increased compared to those produced in the feeding experiment.

**Fig. 3.**
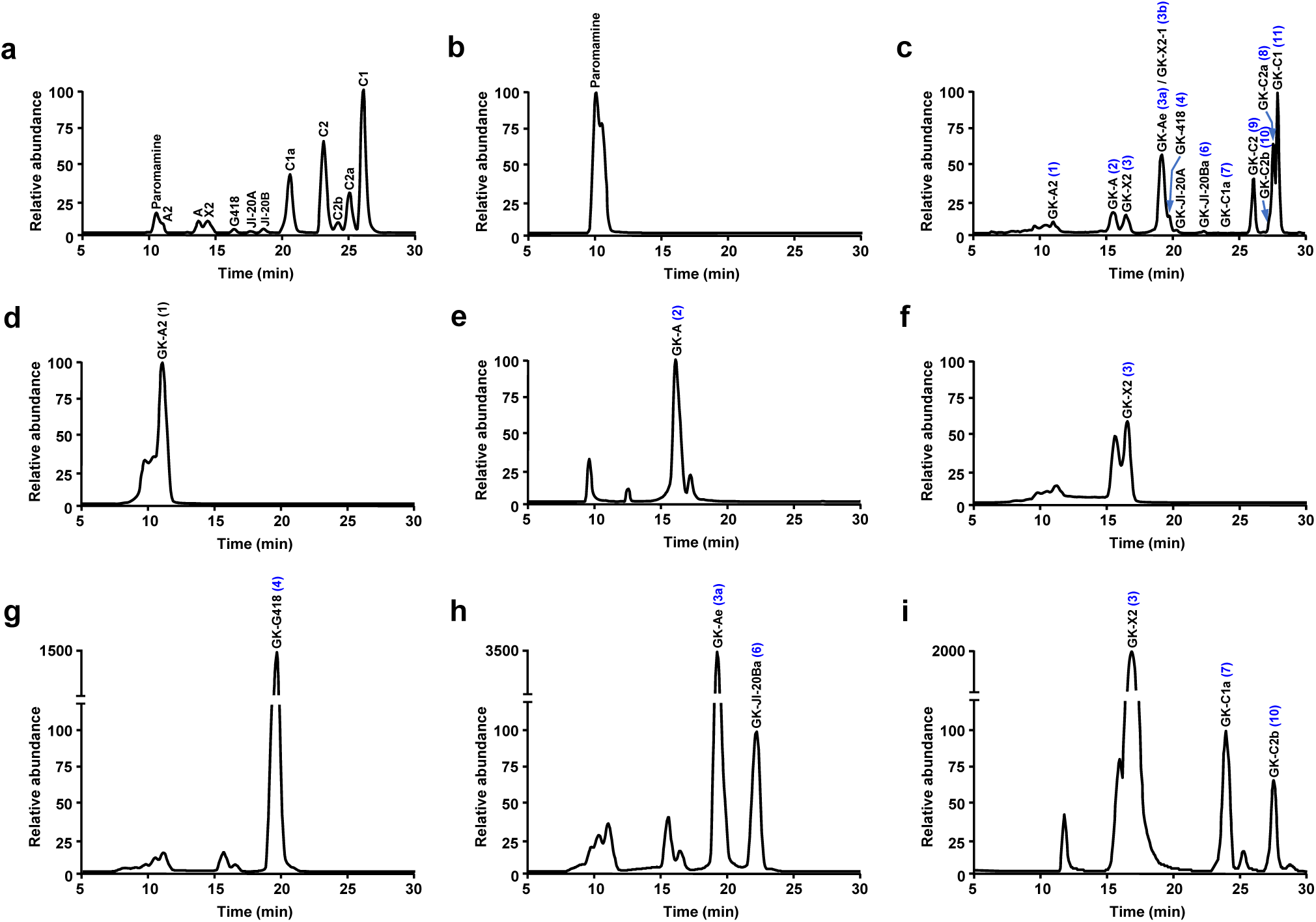
LC-ESI-HRMS analysis of gentamicins production in wild-type and GKs production in engineered mutants. Extracted ion chromatogram traces of gentamicins and proposed GK products in the fermentation culture extracts from (**a**) wild-type, (**b**) ΔgenM2, (**c**) ΔgenM2::kanE, (**d**) ΔgenS2ΔgenKΔgenM2::kanE, (**e**) ΔgenD1ΔgenKΔgenM2::kanE, (**f**) ΔgenQΔgenKΔgenM2::kanE, (**g**) ΔgenM2ΔgenQ::kanE, (**h**) ΔgenM2ΔgenB3::kanE, (**i**) ΔgenM2ΔgenK::kanE.

In comparison to the intermediates, the C-complex products are the primary gentamicin components produced in the wild-type *M. echinospora* (Fig. 3a). It has been noted that the right branch C-complex products, gentamicin C2, C2a, and C1, are produced at higher levels than the left branch ones, C1a and C2b. The composition of the majority of GK intermediates and C-complex components produced in ΔgenM2::kanE followed a similar pattern to the one of gentamicins production in wild-type strain, except for a surprisingly high level of an unknown intermediate with the same *m/z* of GK-X2 (**3**) and the trace amount of GK-C1a (**7**) (Fig. 3c). HPLC-MS/MS analysis of this unknown species suggested it is a structural isomer of GK-X2 (**3**). Unlike GK-X2 (**3**), which has both methyl groups installed on ring III, MS/MS fragmentation indicated that the two methyl groups in this intermediate are separately on rings I and III. (Supplementary Fig. 4). Our previous studies on minor methylation pathways in gentamicins biosynthesis uncovered two gentamicin X2 isomers, gentamicin Ae and X2-1 (Supplementary Fig. 5). We proposed that the biosynthetic route of this intermediate akin to that of gentamicin Ae or X2-1, making its structure likely to be GK-Ae (**3a**) or GK-X2-1 (**3b**). Elevated levels of **3a / 3b** could divert the metabolic flux to the right branch of the proposed GK pathway, which is consistent with the decreased yield of left branch products, GK-C1a (**7**) and GK-C2b (**10**), when compared to gentamicin C1a and C2b production in the wild-type strain.

The above results demonstrated that endogenously produced the kanamycin family pseudotrisaccharides via GT swapping approach were recognized by all the modification enzymes and modified with comparable efficiency to the native substrates. Furthermore, the combinatorial biosynthesis approach not only generated a greater diversity of novel products compared to the feeding process, resulting in the creation of 12 new GK products but also significantly higher production levels for these products.

### Targeted accumulation of GK components by metabolic engineering

*In vivo* production of a series of structurally similar GKs in ΔgenM2::kanE made the isolation, structural confirmation, and activity assessment of individual mono-components, particularly those with lower yields, extremely challenging. This promoted us to focus on targeted accumulating GK species. The successful production of all the proposed GK products, coupled with the observation that the production level of the majority of the species mirrors those of the corresponding gentamicin species, suggested that the GK biosynthetic pathway in ΔgenM2::kanE is likely mimicking the one of gentamicin. Our previous work on characterizing the gentamicin pathway generated a series of gene(s) deletion mutants that individually accumulate various gentamicin components. Based on those mutants, we hypothesized that targeted accumulation of corresponding GK products could be achieved via metabolic engineering.

The three *M. echinospora* mutants, ΔgenS2ΔgenK, ΔgenD1ΔgenK, ΔgenQΔgenK, and ΔgenQ were found to eliminate the downstream and/or minor parallel pathways’ metabolites and direct the metabolic flux toward the endpoint, resulting in exclusive accumulation of gentamicin A2, A, X2, and G418, respectively^14,17,18^. Accordingly, to facilitate the isolation of corresponding GK compounds, we constructed ΔgenS2ΔgenKΔgenM2::kanE, ΔgenD1ΔgenKΔgenM2::kanE, ΔgenQΔgenKΔgenM2::kanE and ΔgenQΔgenM2::kanE, respectively (Supplementary Fig. 3). HPLC-ESI-HRMS analysis of these mutants showed that, while ΔgenS2ΔgenKΔgenM2::kanE, ΔgenD1ΔgenKΔgenM2::kanE, and ΔgenQΔgenM2::kanE successfully accumulated GK-A2 (**1**), GK-A (**2**), and GK-418 (**4**) at a 10-fold, 9-fold and 20-fold level, respectively, and with no substantial rise in the upstream products production in all three cases, the ΔgenQΔgenKΔgenM2::kanE, on the other hand, increased the yield of both GK-X2 (**5**) and the upstream GK-A (**2**) (Fig. 3d-g).

The three GK products from the left branch pathway, including GK-JI20A (**5**), GK-C1a (**7**), and GK-C2b (**10**), along with one right branch intermediate, GK-JI20Ba (**6**), were produced in trace amount in ΔgenM2::kanE. These products are in lower positions in the proposed pathway, making it difficult to accumulate them through metabolic engineering exclusively. Nevertheless, their production can still be boosted by blocking the downstream or the right paradelle pathways. Gene deletion of *gen*B3 was found to raise the production of both immediate upstream intermediates, JI-20A and JI20Ba, in *M. echinospora*^17^. ΔgenB3ΔgenM2::kanE was accordingly constructed and found to increase the production of GK-JI20Ba (**6**) by 70 folds compared to ΔgenM2::kanE (Fig. 3h). Surprisingly, while no change in the GK-JI20A (**5**) yield was observed, the production of the GK-X2 isomer **3a** increased by 50 folds in ΔgenB3ΔgenM2::kanE compared to ΔgenM2::kanE. The methyltransferase GenK is a critical enzyme that directs the gentamicin biosynthesis to the right branch pathway. Deletion of *genK* in *M. echinospora* led to the elimination of gentamicin right branch products and largely lifted the production of all the right branch ones^18^. We subsequently constructed ΔgenKΔgenM2::kanE, which brought up the yield of the two C-complex products, GK-C1a (**7**) and GK-C2b (**10**), by approximately 30-fold and 5-fold, respectively. Interestingly, although the GK-X2 (**3**) exhibited an accumulation increase of around 100-fold in this mutant, the downstream GK-JI20A (**5**) still remained at a trace level (Fig. 3i).

### Purification and structural characterization of six GK compounds

To facilitate the isolation of mono GK components, the engineered mutants were initially subjected to 2L scale lab-condition fermentation using a modified industrial formulation medium (F50), which reduces all ingredients by 50% and incorporates filtered broth of soy powder to lower viscosity. Fermentation extracts were purified successively through ion-exchange and cation-exchange resins prior subjected to mono components preparation using an LC-evaporative light scattering detector (ELSD). However, owing to the lack of an optimal separation of the target components from impurities, GK-A2 (**1**), GK-A (**2**), and GK-JI20B (**6**) could not be obtained in the required purity for structural and activity characterization (Supplementary Fig. 5). Additionally, the yields of GK-C2a (**8**) in ΔgenM2::kanE and GK-C2b (**10**) in ΔgenKΔgenM2::kanE were insufficient to support the effective purification of an adequate quantity of both compounds based on the 2L lab-condition fermentation (Supplementary Fig. 5). Ultimately GK-418 (**4**) (derived from ΔgenQΔgenM2::kanE), GK-X2 (**3**) and GK-C1a (**7**) (derived from ΔgenKΔgenM2::kanE) and GK-C1 (**11**) (derived from ΔgenM2::kanE) were successfully isolated in satisfactory quantities and purities under the laboratory setting (Supplementary Fig. 6).

The C-complex products constitute the primary components of clinically used gentamicin mixtures and have also been reported to exhibit superior antibacterial activities compared to the intermediates. We hypothesized that the activity profile of GK products follows a similar pattern. Consequently, we turned to large-scale industrial fermentation to aid in preparing the remaining two C-complex GK products, GK-C2a (**8**) and GK-C2b (**10**), which were not achievable in the lab setting. This was conducted through fermentation of ΔgenM2::kanE and ΔgenKΔgenM2::kanE, respectively, using a full formulation medium in a 30 L scale bioreactor.

Notably, both mutants displayed a different metabolite profile compared to the lab-condition fermentation. The metabolite flux in both mutants was predominantly directed towards endpoint C-complex products during industrial fermentation while substantial amounts of intermediates remained converted in the lab settings (Supplementary Fig. 5d, e). This could potentially be attributed to an extended bacterial growth phase and sustained secondary metabolism facilitated by a continuous nutrient supply and tightly controlled conditions in the bioreactor fermentation. This led to a significant increase in the production of GK-C2a (**8**) in ΔgenM2::kanE and facilitated its subsequent isolation. Interestingly, though the yield of GK-C1a (**7**) was greatly raised in ΔgenKΔgenM2::kanE during industrial fermentation, it did not convert to the downstream GK-C2b (**10**), which is in contrast to what was observed in lab-condition fermentation. While industrial fermentation of ΔgenKΔgenM2::kanE made the GK-C1a (**7**) purification more efficient, we were not able to obtain GK-C2b (**10**) eventually.

Overall, six GK mono-components, including GK-X2 (**3**), GK-418 (**4**), GK-C1a (**7**), GK-C2a (**8**), GK-GK-C2 (**9**), and C1 (**11**) were purified, and their structures were confirmed by 1D and 2D-NMR characterization (Supplementary Table 3-8 and Supplementary Fig. 7-54).

### GKs display promising antimicrobial activity

The six purified GKs were subsequently screened for antimicrobial activity against the ESCAPE panel pathogens, including *<u>E</u>nterococcus faecium*, *<u>S</u>taphylococcus aureus*, *<u>K</u>lebsiella pneumoniae*, *<u>A</u>cinetobacter baumannii*, *<u>P</u>seudomonas aeruginosa* and *<u>E</u>nterobacter cloacae*. The minimal inhibitory concentration (MIC) of each compound against the reference strain recommended by the Clinical and Laboratory Standards Institute (CLSI) for each pathogen was determined using a microbroth dilution assay following the CLSI guidelines. To compare with the clinically used AGs and offer insights into the structural-activity relationship, the MIC assay included the six corresponding gentamicin mono components, kanamycin B, and the synthetic kanamycin derivative, dibekacin (Table 1).

**Table 1.**
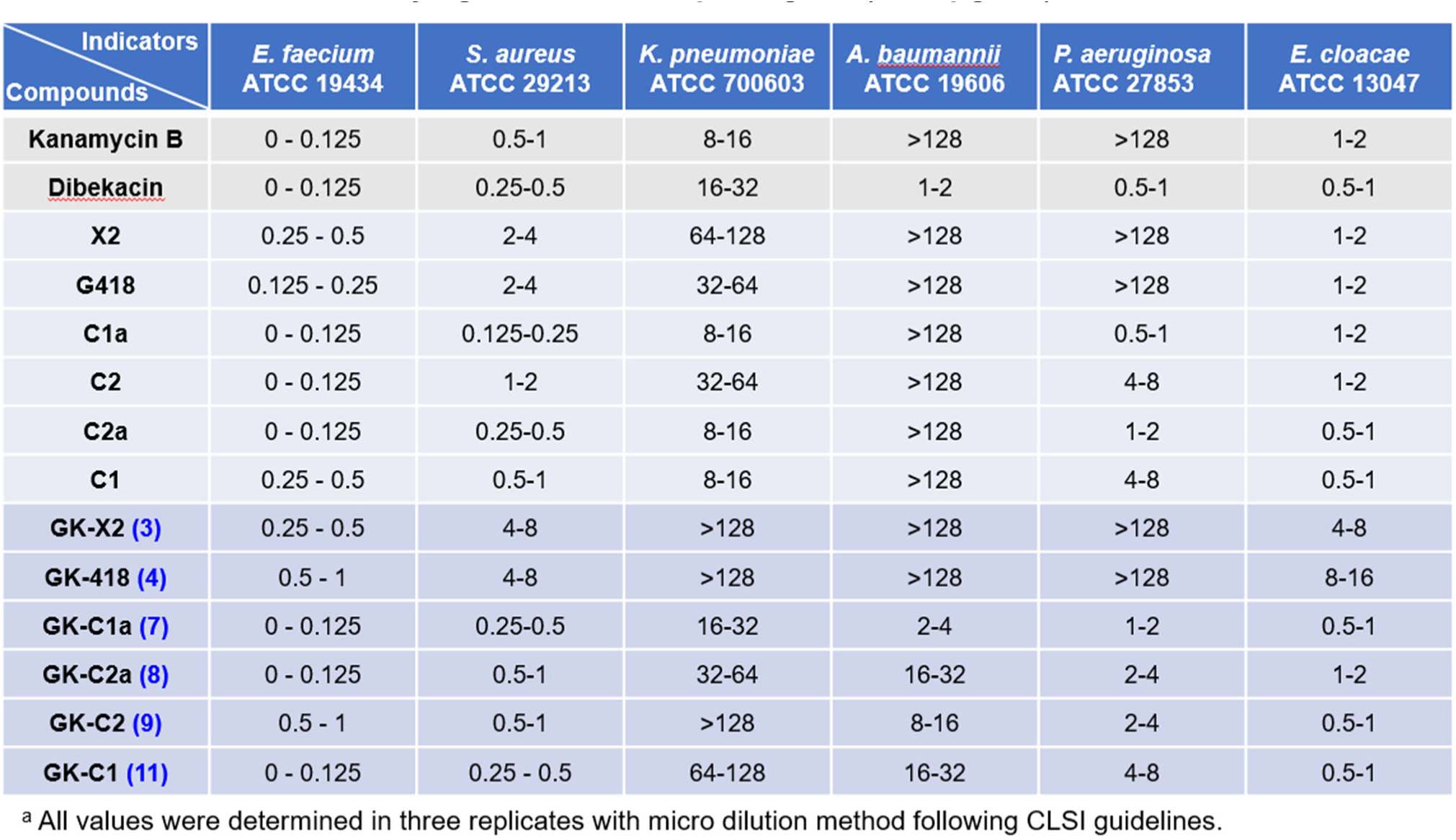
Antimicrobial activity against ESKAPE pathogens (MIC, μg/mL)^a^.

In general, all the GK C-complex products displayed potent antimicrobial activities against four out of the six tested strains, including *E. faecium* ATCC 19434, *S. aures* ATCC 29213, *P. aeruginosa* ATCC 27853 and *E. cloacae* ATCC 13407. Similar to the gentamicin intermediates, GK-X2 (**3**) and GK-418 (**4**) only showed potent activity against *E. faecium* ATCC 19434, *S. aures* ATCC 29213, and *E. cloacae* ATCC 13407. Additionally, consistent with the trend for gentamicins, the GK C-complex products exhibited superior activity against all the pathogens than the two tested intermediates, indicating that modifications increase the antimicrobial activity. This was particularly true for the case of *P. aeruginosa* ATCC 27853. While it was not sensitive to all tested gentamicin and GK intermediates and kanamycin B, all gentamicin and GK C-complex species, as well as dibekacin, displayed potent activity against it. This suggested that the 3,4-didehydroxylation for both kanamycin and gentamicin pseudotrisaccharide scaffolds is critical for the activity against *P. aeruginosa* ATCC 27853. In most cases, GK products showed slightly decreased activities compared to the corresponding gentamicin species against all tested pathogens (except for *A. baumannii* ATCC 19606), implying that swapping the xylose with glucose in ring III may cost the activity. It is worth noting that, while *A. baumannii* ATCC 19606 was found to be resistant to all forms of gentamicin and kanamycin B, both dibekacin and the four GK C-complex components presented promising inhibition activities. Through the structural comparison, this result suggested that both 3,4-didehydroxylation on ring II and glucose instead of xylose as ring III are essential structure features for inhibiting *A. baumannii* ATCC 19606. All the tested AGs showed only moderate or no activity against *K. pneumoniae* ATCC700603.

### GK-C2a shows low ototoxicity in zebrafish embryos

Ototoxicity has been a long-standing issue of AGs’ clinical application due to their irreversible damage to inner hair cells (HCs), and thus arouse more concern than their nephrotoxic side-effects, which are reversible. Zebrafish is an excellent and powerful tool for ototoxicity tests due to its surficial sensory lateral line HCs and inner ear HCs, similar to mammalian HCs in morphology and function^[7,8]^. Here, the transgenic line *Tg(Brn3c:mGFP)* larvae were utilized to comprehensively assess the toxicity of GK C-complex mono components to HCs in neuromasts of the posterior lateral line (pLL) at concentrations ranging between 0-100 μM (Fig. 4 and Supplementary Fig. 55). As comparison, the four corresponding gentamicin C-complex components, kanamycin B, and dibekacin were included in the test. Results showed that the tested AGs induced the loss of HCs exhibiting the dose-dependent mode and their toxicity to HCs notably varied (Fig. 4c-i). Generally, the two compounds belonging to the kanamycin family exhibited higher toxicity than both gentamicins and GKs across all tested concentrations (Fig. 4c-e). In evaluating the four GKs C-complex compounds, it suggested that GK-C2a displayed the least ototoxicity among them (Fig. 4c). A comparative analysis of the corresponding C-complex species between the GK and gentamicin families revealed that GK-C1a (**7**), GK-C2 (**9**) and GK-C1 (**11**) exhibited toxicity levels comparable to gentamicin C1a, C2 and C1, respectively, while GK-C1 only demonstrated increased toxicity than gentamicin C1 at concentrations between 10-100 μM (Fig. 4f-i). Remarkably, GK-C2a (**8**) exhibited a significantly reduced toxicity profile compared to gentamicin C2a, particularly at concentrations ≥ 2.5 μM (Fig. 4g). While possessing potent antimicrobial activity against multiple pathogens in ESKAPE panel (Table 1), GK-C2a (**8**) demonstrated an ototoxicity profile comparable to that of the least ototoxic gentamicin C-complex species, gentamicin C1, suggesting the promising potential of GK-C2a (**8**) as a mono-component in clinical applications.

**Fig. 4.**
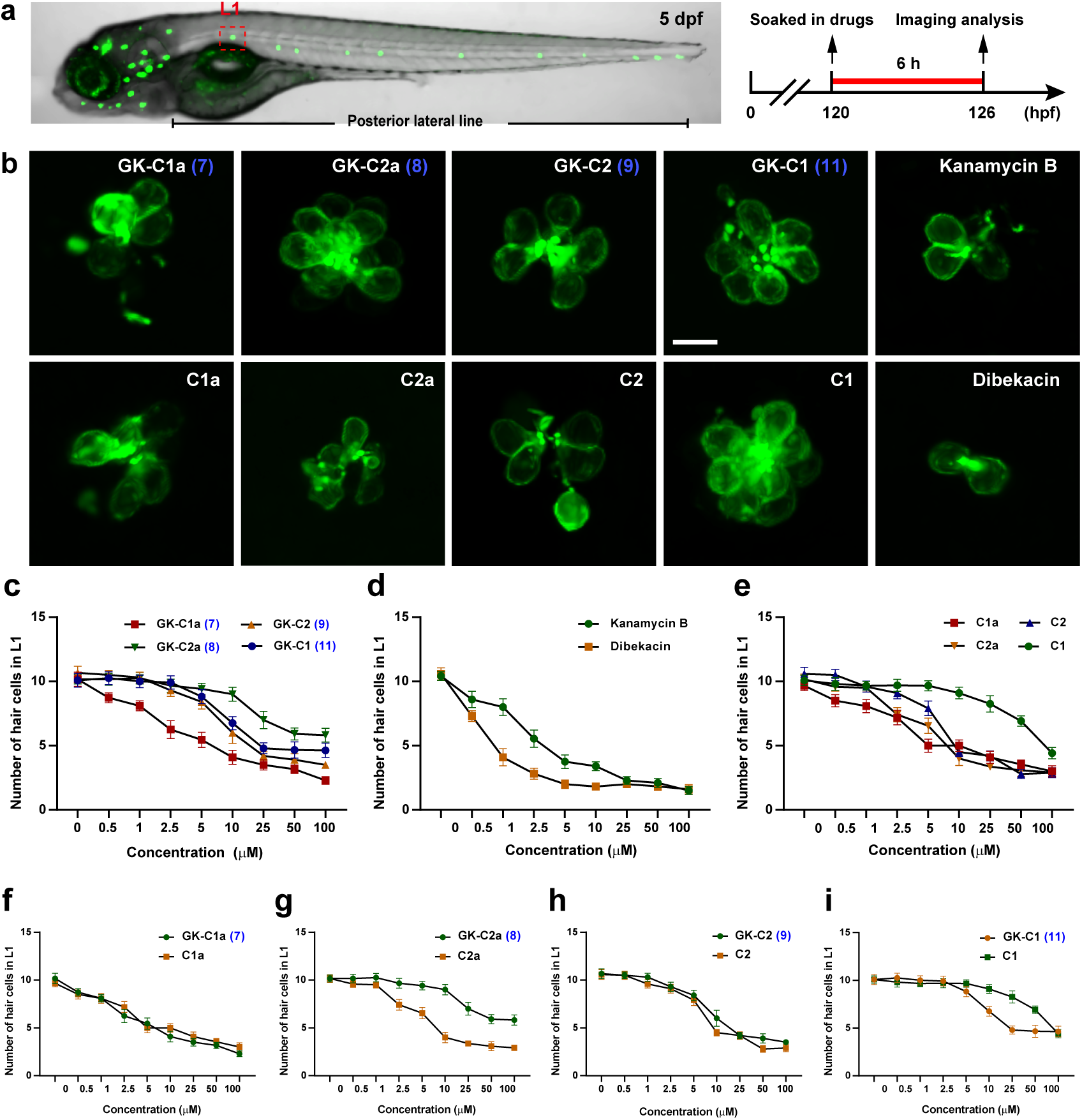
The toxicity assessment of AG compounds to HCs in pLL of zebrafish. (**a**) Confocal image of a *Tg(Brn3c:mGFP)* larvae at 5 d post-fertilization (dpf) and schematic diagram of methods used in toxicity testing. The neuromast in L1 (lateral 1) of pLL analyzed in this experiment was marked by red dashed box. (**b**) Representative fluorescent images of HC cluster in L1 neuromast of pLL under different compounds treatment at concentration of 10 μM. Scale bar: 10 μm. (**c**-**i**) Statistical results of the number of HCs in L1 neuromast under different compounds treatment with a series of concentrations (*n* = 10).

To further characterize the ototoxicity of GK-C2a in larvae, the HCs existing in the main auditory organs of zebrafish, including functional HCs in neuromasts of pLL and cristae HCs in the otic vesicle, were analyzed under treatment of GK-C2a (**8**). This was also compared with gentamicin C2a, kanamycin B and semi-synthetic derivative drug, dibekacin, in attempt to provide insights into structure and ototoxicity relationship (Fig. 5). Results showed that the four AGs compounds caused the reduced functional HCs, while the number of functional HCs and GFP-labeled HCs under GK-C2a (**8**) treatment was dramatically higher than that of others (Fig. 5b, d). The influence of AG compounds on HCs in the inner ear was also examined by microinjection of the compounds into otic vesicles (Fig. 5a). As shown, the semicircular canal filled with Texas Red labeled dextran indicated that the drugs were successfully conveyed (Fig. 5c). Images showed that there were obviously damaged HCs in anterior cristae (AC), lateral cristae (LC), and posterior cristae (PC) under treatment. However, the damages varied slightly in GK-C2a (**8**) while seriously in dibekacin (Fig. 5c). The number of three clusters of crista HCs was analyzed and consistent statistical results were also shown (Fig. 5e). To investigate the effect of GK-C2a (**8**) on auditory function of larvae, a startle response assay was performed (Supplementary Fig. 56). The results indicated that larvae exposed to GK-C2a (**8**) were more sensitive to external sound stimuli embodied in higher swimming distance and peak velocity compared to other compounds (Fig. 5f, g and Supplementary Fig. 56).

**Fig. 5.**
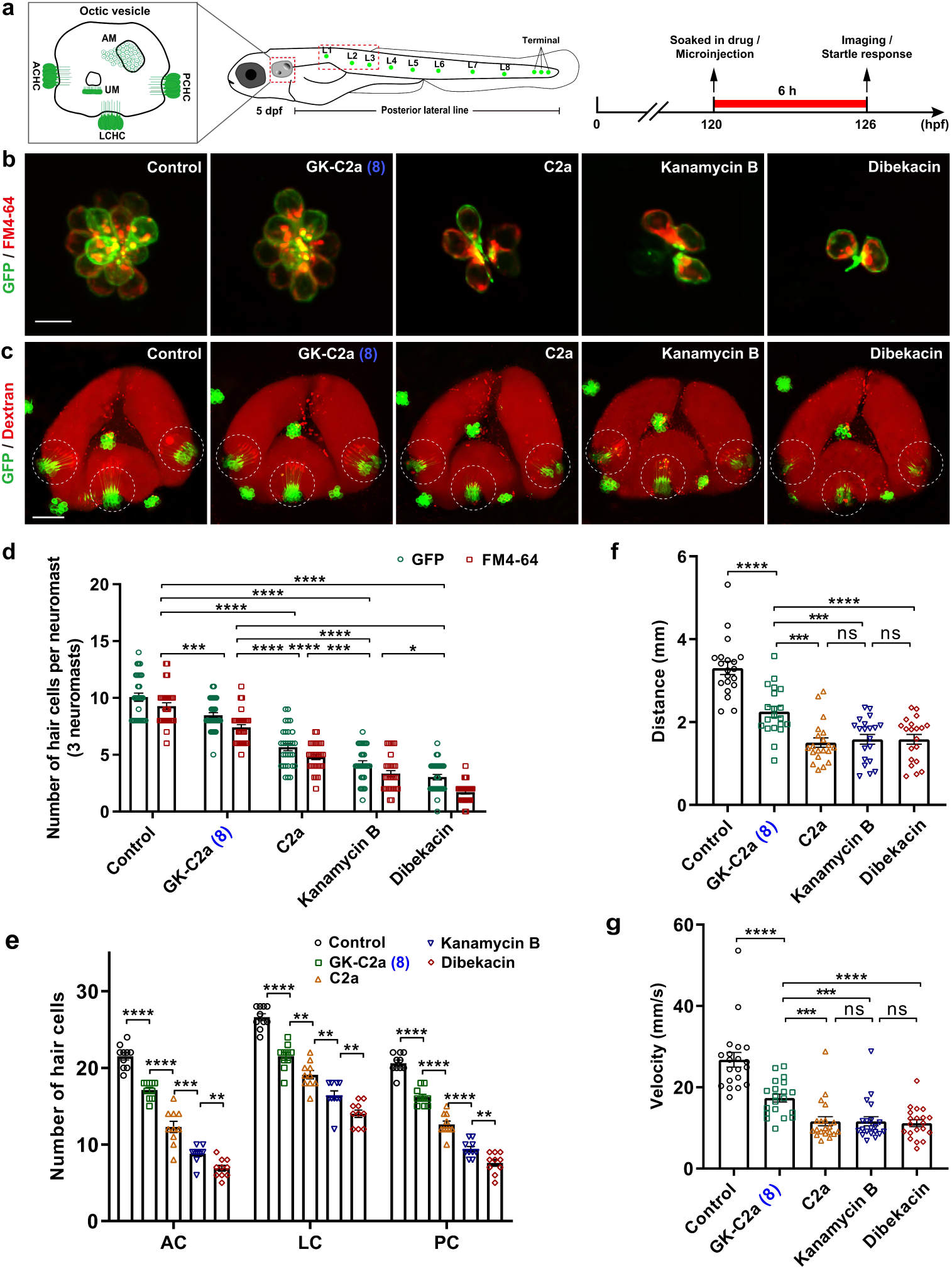
Toxicity assessment of GK-C2a (8) in zebrafish HCs morphology and function. (**a**) Schematic structure of main auditory organs in zebrafish at 5 dpf, including otic vesicle and pLL system. Green patches indicated the locations of neuromasts in pLL. The neuromasts in L1, L2 and L3 analyzed in this experiment were marked by red dashed box. Detailed structure of otic vesicle was depicted. ACHC, anterior crista hair cell; LCHC, lateral crista hair cell; PCHC, posterior crista hair; UM, utricular macula; AM, saccular macula. (**b**) Representative merged fluorescent images of HC cluster (green color) and functional HC cluster (red color) in L1 neuromast under 10 μM different compounds treatment. Scale bar: 10 μm. (**c**) Representative merged fluorescent images of crista HC clusters (marked by white dot circle lines) in otic vesicles treated with 10 mg/mL different compounds. Scale bar: 50 μm. (**d**) Statistical results of the number of HCs and functional HCs per neuromast (L1, L2 and L3) under 10 μM different compounds treatment (*n* = 30). (**e**) Statistical results of the number of cristae HCs in AC, LC, and PC regions (*n* = 10). (**f**, **g**) Statistical results of swimming distance and velocity in startle response experiment exposed to 9 dB re ms^-2^ sound stimulus (*n* = 20). **, ***, and **** represented *P* < 0.01, *P* < 0.001, and *P* < 0.0001, respectively.

In summary, these results sufficiently indicated that the novel aminoglycoside GK-C2a (**8**), displayed low ototoxicity in zebrafish embryos. It also indicated that swapping the second sugar from xylose to glucose in gentamicins provides comparable or reduced ototoxicity, while modification on kanamycin scaffolds can significantly reduce the ototoxicity.

## Discussions

AGs have been essential antibiotics for treating bacterial infections for decades. However, their clinical use is hindered by toxicity concerns as well as ever-growing resistance. Nevertheless, AGs continue to be clinically valuable due to their potent and broad-spectrum antimicrobial activities, especially their potential renewed application in the anti-cancer treatment that has been discovered. To overcome the limitations and aid in new applications of these important family compounds, the development of AG structural derivative therefore remains imperative.

Semi-synthetic approaches via direct modification of the natural AGs have been utilized to develop the second and third generation AGs in the early years. Despite a few successes, such as dibekacin, etilmicin, and plazomicin, chemically regiospecific modification of densely functionalized and structurally diverse AGs remains challenging and consequently leads to low efficiency. A few successes in the chemoenzymatic modification of natural AGs using enzymes from heavily modified natural AG biosynthesis in the early years offered an appealing alternative and demonstrated the potential of harnessing the microorganisms’ biosynthetic machinery to create novel AGs. However, they were still limited by challenges in production in scale due to the low yields.

The extensive knowledge acquired regarding understanding the biosynthetic pathway of the most heavily modified natural AG, gentamicins, opens up new opportunities to employ a combinatory biosynthesis strategy for AG structural diversification. Based on the biosynthetic logic of gentamicins, the modifications all occur after the pseudotrisaccharide formation, and with substrate promiscuity previously revealed to multiple modification enzymes in their biosynthesis, it leads to a potential glycodiversification strategy for the bioengineering the AGs.

In this study, we explored the novel approach that centered on gentamicin glycodiversification through GT swapping in bacterial producers to diversify the structures of AGs efficiently. As a preliminary attempt, through a feeding experiment, we showed that gentamicin-producing strains can modify kanamycin B, resulting in four new hybrid species. This demonstrated the feasibility of combining the biosynthetic pathways of gentamicins and kanamycins by swapping gentamicin pseudotrisaccharide formation GT with the corresponding one from kanamycins biosynthesis to create their novel hybrid AGs. Indeed, the mutant ΔgenM2::kanE successfully led to the production of a series of proposed GKs. Metabolic engineering of the GK pathway is then employed for the targeted accumulation of specific GK products, making their isolation and structural characterization feasible. Six genetically engineered mutants were constructed to target the accumulation of GK components that facilitated the purification of individual GK species. Six GK compounds were eventually isolated and structurally characterized via combination of fermentation in the lab and industrial settings. The large-scale fermenter production of the GK-C complex components from ΔgenM2::kanE and ΔgenM2ΔgenK::kanE showcased the potential in green manufacturing of these valuable AGs. The AHBA chain at the *N*-1 position of the 2-DOS ring is a structural feature of the new generation of AGs, including amikacin, arbekacin, and plazomicin demonstrating improved activity against many resistant strains. The novel GKs can serve as novel structural leads for further modifications via semi-synthesis to attach the AHBA chain side chain.

The antimicrobial activity of the six GK compounds was evaluated against a panel of WHO’s critical priority pathogens, the ESKAPE panel, which are six highly virulent pathogens with increasing multi-drug resistance revealed. The results demonstrated that GKs retained promising antimicrobial activity as clinically used gentamicin and kanamycin family compounds against ESKAPE panel pathogens, with C-complex products showing the highest activity. While GK intermediates, GK-X2 and GK-418, display limited activity against some strains, this was not unexpected given the similar antimicrobial profiles of corresponding gentamicin species. The antimicrobial activity of GK species was also compared to that of the natural gentamicins and kanamycin B, as well as a clinically used semi-synthetic AG, dibekacin. Notably, the four GK C-complex components, particularly GK-C1a, presented promising inhibition activities against *A. baumannii* ATCC 19606, while the pathogen was found to be resistant to all forms of gentamicins and kanamycin B. The activity comparison revealed a few structural-activity relationships. Swapping the xylose with glucose in ring III of gentamicins has different effects on the activities of each species against different pathogen species. Extensive modifications on kanamycin scaffolds, leading to the GK C-complex compounds, increase the activity against all tested pathogens except for *K. pneumoniae* ATCC700603. While 3,4-didehydroxylation generally increases the activities of all three family AGs against all pathogens tested, it is critical for the activity against *P. aeruginosa* ATCC 27853. Both 3,4-didehydroxylation and having glucose instead of xylose as ring III are essential structural features for inhibiting *A. baumannii* ATCC 19606. The promising antimicrobial activities of the obtained GK-C-complex products against the reference strain of the ESKAPE panel pathogens suggest their potential to combat the drug-resistant species, especially the ones of *A. baumannii,* which shows high resistance to almost all AGs. It is, therefore, worth testing the antimicrobial activity of obtained GK-C-complex products against clinically isolated drug-resistant ESKAPE panel pathogens in future work.

Zebrafish embryos were used as a model system to assess the ototoxicity of the four GK C-complex compounds in comparison with the four gentamicin C-complex components, kanamycin B, and dibekacin. Remarkably, extensive modifications on kanamycin scaffolds leading to GK C-complex compounds significantly reduced the ototoxicity in zebrafish embryos. Commercial gentamicin is a complex of several compounds with major ingredients as C-complex species. Different ototoxicity in cochlear explants of mono gentamicin C-complex components has been reported.^21^ This is consistent with our findings of the corresponding GK C-complex components showing different ototoxic profiles. The four GK-C complex components showed comparable ototoxicity to that of the corresponding gentamicin species. Notably, while GK-C2a showed potent antimicrobial activity against four out of six ESKAPE pathogens, toxicity comparative data revealed that it displayed much lower auditory toxicity than the corresponding gentamicin C2a, and has comparable ototoxicity to the least ototoxic gentamicin species, gentamicin C1, in the test model. As ototoxicity and antimicrobial activity do not directly correlate for the C-complex compounds of both family AGs, it is possible to reformulate these AGs with less ototoxic components without altering antimicrobial activity.

Overall, the study demonstrates the potential of glycosyltransferase swapping to diversify AG structures, leading to the creation of novel hybrid AGs with promising antimicrobial activity, particularly GK-C2a, with low toxicity, making it a candidate for further development as an antibiotic with improved safety profiles. This research offers a valuable contribution to developing AG antibiotics in the face of drug toxicity and antibiotic resistance concerns.

## Acknowledgments

This work was supported by the National Key R&D Program of China (2018YFA0903200) and the Funds for International Cooperation and Exchange of the National Natural Science Foundation of China (31920103001).

## Author contributions

C.W. did ototoxicity assays of several compounds in zebrafish model. S.W. performed antimicrobial activity against ESKAPE pathogens. G.S. identified the structures of GKs. X.J., C.H., C.Q. and Y.L. carried out mutant construction, fermentation and product purification, respectively. D.L., F.Z. P.F.L and Z.D. participated in discussion, data analysis and manuscript revision. Y.S. and X.J. conceived the overall project, designed the experiment, analyzed the data, and wrote the manuscript.

## Additional information

Correspondence and requests for materials should be addressed to Y.S.

