## Supporting information for "Glycodiversification of gentamicins through *in vivo* glycosyltransferase swapping enabled the creation of novel hybrid aminoglycoside antibiotics with potent activity and low ototoxicity"

### Supplementary Information

#### Table of Contents

#### Materials and Methods

#### Supplementary Tables

**Supplementary Table 1.** The bacterial strains and plasmids used in this study.
**Supplementary Table 2.** Oligonucleotide primers used in this study.
**Supplementary Table 3.** $^{1}H$ NMR (600 MHz, $D_2O$) and $^{13}C$ NMR (150 MHz, $D_2O$) data of GK-X2 (**3**).
**Supplementary Table 4.** $^{1}H$ NMR (600 MHz, $D_2O$) and $^{13}C$ NMR (150 MHz, $D_2O$) data of GK-418 (**4**).
**Supplementary Table 5.** $^{1}H$ NMR (600 MHz, $D_2O$) and $^{13}C$ NMR (150 MHz, $D_2O$) data of GK-C1a (**7**).
**Supplementary Table 6.** $^{1}H$ NMR (600 MHz, $D_2O$) and $^{13}C$ NMR (150 MHz, $D_2O$) data of GK-C2a (**8**).
**Supplementary Table 7.** $^{1}H$ NMR (600 MHz, $D_2O$) and $^{13}C$ NMR (150 MHz, $D_2O$) data of GK-C2 (**9**).
**Supplementary Table 8.** $^{1}H$ NMR (600 MHz, $D_2O$) and $^{13}C$ NMR (150 MHz, $D_2O$) data of GK-C1 (**11**).

#### Supplementary Figures

**Supplementary Fig. 1** Comprehensive biosynthetic pathway of gentamicin and kanamycin complex.
**Supplementary Fig. 2** Predicted structures and their observed ions by LC-ESI-HRMS and MS/MS analysis of fermentation extract of ΔgenM2 fed with kanamycin B.
**Supplementary Fig. 3** Genetic confirmation of in-frame gene deletion and *kanE* complementation mutants.
**Supplementary Fig. 4** LC-ESI-HRMS and MS/MS analysis of GK C-complex components and intermediates in ΔgenM2::kanE.
**Supplementary Fig. 5** Proposed methylation pathway in gentamicin and GKs biosynthesis.
**Supplementary Fig. 6** HPLC-ELSD chromatogram of culture extracts of GKs producing and accumulating mutants.
**Supplementary Fig. 7** Chemical structure, key $^{1}H$–$^{1}H$ COSY and HMBC correlations of GK-X2 (**3**).
**Supplementary Fig. 8** ESI-HRMS spectrum of GK-X2 (**3**).
**Supplementary Fig. 9** $^{1}H$ NMR spectrum (600 MHz, $D_2O$) of GK-X2 (**3**).
**Supplementary Fig. 10** $^{13}C$ NMR, DEPT-90 and DEPT-135 spectra (150 MHz, $D_2O$) of GK-X2 (**3**).
**Supplementary Fig. 11** HSQC spectrum (600 MHz, $D_2O$) of GK-X2 (**3**).
**Supplementary Fig. 12** $^{1}H$–$^{1}H$ COSY spectrum (600 MHz, $D_2O$) of GK-X2 (**3**).
**Supplementary Fig. 13** HMBC spectrum (600 MHz, $D_2O$) of GK-X2 (**3**).
**Supplementary Fig. 14** NOESY spectrum (600 MHz, $D_2O$) of GK-X2 (**3**).
**Supplementary Fig. 15** Chemical structure, key $^{1}H$–$^{1}H$ COSY, HMBC and NOESY correlations of GK-418 (**4**).
**Supplementary Fig. 16** ESI-HRMS spectrum of GK-418 (**4**).
**Supplementary Fig. 17** $^{1}H$ NMR spectrum (600 MHz, $D_2O$) of GK-418 (**4**).
**Supplementary Fig. 18** $^{13}C$ NMR, DEPT-90 and DEPT-135 spectra (150 MHz, $D_2O$) of GK-418 (**4**).
**Supplementary Fig. 19** HSQC spectrum (600 MHz, $D_2O$) of GK-418 (**4**).
**Supplementary Fig. 20** $^{1}H$–$^{1}H$ COSY spectrum (600 MHz, $D_2O$) of GK-418 (**4**).

**Supplementary Fig. 21** HMBC spectrum (600 MHz, $D_2O$) of GK-418 (**4**).
**Supplementary Fig. 22** NOESY spectrum (600 MHz, $D_2O$) of GK-418 (**4**).
**Supplementary Fig. 23** Chemical structure, key $^1$H–$^1$H COSY and HMBC correlations of GK-C1a (**7**).
**Supplementary Fig. 24** ESI-HRMS spectrum of GK-C1a (**7**).
**Supplementary Fig. 25** $^1$H NMR spectrum (600 MHz, $D_2O$) of GK-C1a (**7**).
**Supplementary Fig. 26** $^{13}$C NMR, DEPT-90 and DEPT-135 spectra (150 MHz, $D_2O$) of GK-C1a (**7**).
**Supplementary Fig. 27** HSQC spectrum (600 MHz, $D_2O$) of GK-C1a (**7**).
**Supplementary Fig. 28** $^1$H–$^1$H COSY spectrum (600 MHz, $D_2O$) of GK-C1a (**7**).
**Supplementary Fig. 29** HMBC spectrum (600 MHz, $D_2O$) of GK-C1a (**7**).
**Supplementary Fig. 30** NOESY spectrum (600 MHz, $D_2O$) of GK-C1a (**7**).
**Supplementary Fig. 31** Chemical structure, key $^1$H–$^1$H COSY, HMBC and NOESY correlations of GK-C2a (**8**).
**Supplementary Fig. 32** ESI-HRMS spectrum of GK-C2a (**8**).
**Supplementary Fig. 33** $^1$H NMR spectrum (600 MHz, $D_2O$) of GK-C2a (**8**).
**Supplementary Fig. 34** $^{13}$C NMR, DEPT-90 and DEPT-135 spectra (150 MHz, $D_2O$) of GK-C2a (**8**).
**Supplementary Fig. 35** HSQC spectrum (600 MHz, $D_2O$) of GK-C2a (**8**).
**Supplementary Fig. 36** $^1$H–$^1$H COSY spectrum (600 MHz, $D_2O$) of GK-C2a (**8**).
**Supplementary Fig. 37** HMBC spectrum (600 MHz, $D_2O$) of GK-C2a (**8**).
**Supplementary Fig. 38** NOESY spectrum (600 MHz, $D_2O$) of GK-C2a (**8**).
**Supplementary Fig. 39** Chemical structure, key $^1$H–$^1$H COSY, HMBC and NOESY correlations of GK-C2 (**9**).
**Supplementary Fig. 40** ESI-HRMS spectrum of GK-C2 (**9**).
**Supplementary Fig. 41** $^1$H NMR spectrum (600 MHz, $D_2O$) of GK-C2 (**9**).
**Supplementary Fig. 42** $^{13}$C NMR, DEPT-90 and DEPT-135 spectra (150 MHz, $D_2O$) of GK-C2 (**9**).
**Supplementary Fig. 43** HSQC spectrum (600 MHz, $D_2O$) of GK-C2 (**9**).
**Supplementary Fig. 44** $^1$H–$^1$H COSY spectrum (600 MHz, $D_2O$) of GK-C2 (**9**).
**Supplementary Fig. 45** HMBC spectrum (600 MHz, $D_2O$) of GK-C2 (**9**).
**Supplementary Fig. 46** NOESY spectrum (600 MHz, $D_2O$) of GK-C2 (**9**).
**Supplementary Fig. 47** Chemical structure, key $^1$H–$^1$H COSY, HMBC and NOESY correlations of GK-C1 (**11**).
**Supplementary Fig. 48** ESI-HR MS spectrum of GK-C1 (**11**).
**Supplementary Fig. 49** $^1$H NMR spectrum (600 MHz, $D_2O$) of GK-C1 (**11**).
**Supplementary Fig. 50** $^{13}$C NMR, DEPT-90 and DEPT-135 spectra (150 MHz, $D_2O$) of GK-C1 (**11**).
**Supplementary Fig. 51** HSQC spectrum (600 MHz, $D_2O$) of GK-C1 (**11**).
**Supplementary Fig. 52** $^1$H–$^1$H COSY spectrum (600 MHz, $D_2O$) of GK-C1 (**11**).
**Supplementary Fig. 53** HMBC spectrum (600 MHz, $D_2O$) of GK-C1 (**11**).
**Supplementary Fig. 54** NOESY spectrum (600 MHz, $D_2O$) of GK-C1 (**11**).
**Supplementary Fig. 55** Representative fluorescent images of HC cluster in L1 neuromast of pLL under different compounds treatment.
**Supplementary Fig. 56** The startle response assays of zebrafish larvae (5 dpf) under treatment with four compounds of GK-C2a (**8**), C2a, kanamycin B and dibekacin.

#### Supplementary References

#### Materials and Methods

##### Bacterial strains, chemicals, and culture conditions

Bacterial strains used in this work are listed in the Supplementary Table 1. G418 was purchased from Sigma-Aldrich, gentamicin X2, gentamicin C2, gentamicin C2a, gentamicin C1a and gentamicin C1 were purchased from Toku-E. *M. echinospora* ATCC 15835 wild-type and mutants were grown in liquid ATCC172 medium (glucose 1%, soluble starch 2%, yeast extract 0.5%, N-Z amine type A 0.5%, $CaCO_3$ 0.1%) for genomic DNA isolation and preparation of mycelium. F50 medium (soybean powder 2.0%, peptone 0.1%, glucose 0.3%, soluble starch 3.0%, $(NH_4)_2SO_4$ 0.03%, CaCO3 0.3%, $KNO_3$ 0.03%, $CoCl_2$ 0.005‰) was used for metabolite production. Soybean powder was applied as boiled broth and filtered before addition to the F50 medium. *E. coli* strains were maintained in 2×TY media at 37°C with the appropriate antibiotic selection at a final concentration of 100 μg/mL ampicillin, 25 μg/mL chloramphenicol and 25 μg/mL kanamycin, respectively. ABB medium (soytone 0.5%, soluble starch 0.5%, $CaCO_3$ 0.3%, MOPS, 0.21%, $FeSO_4$ 0.0012%, Thiamine-HCl 0.001%, agar 3%) with addition of 10 mM $MgCl_2$ solution was used for conjugation. A medium (soluble starch 1%, corn steep power 0.25%, yeast extract 0.3%, $CaCO_3$ 0.3%, $FeSO_4$ 0.0012%, agar 3%, pH 7.0, adjusted with KOH) with addition of 10 mM $MgCl_2$ solution was used for nonselective culture of the exconjugants.

##### Mutants construction

**Construction of ΔgenM2::kanE, ΔgenS2ΔgenKΔgenM2::kanE, ΔgenQΔgenKΔgenM2::kanE and ΔgenD1ΔgenKΔgenM2::kanE.** About 2 kb flanking regions of *genM2* were amplified from the wild-type genomic DNA using Phusion DNA polymerase (New England Biolabs) and cloned into the *Streptomyces-E. coli* shuttle vector pYH7 between *Nde*I and *Hin*dIII sites[1] to obtain construct pWHU47. pWHU47 was introduced into the wild-type strain and two previously constructed mutants, ΔgenS2ΔgenK and ΔgenQΔgenK, respectively, by conjugation and mutants screening using the method described before[2] to generate ΔgenM2, ΔgenS2ΔgenKΔgenM2, and ΔgenQΔgenKΔgenM2. Similarly, pWHU2649 was constructed by cloning the ~ 2 kb flanking regions of *genM2* amplified from ΔgenD1ΔgenK genomic DNA into pYH7 between *Nde*I and *Hin*dIII sites and was introduced into previously constructed ΔgenD1ΔgenK[3] to generate ΔgenD1ΔgenKΔgenM2.

Full length *kanE* was amplified from the genomic DNA of kanamycin-producing strain *Streptomyces kanamyceticus* ATCC 12853 using Phusion DNA polymerase and cloned into pWHU77 (between the *Nde*I and *Eco*RI sites) to obtain pWHU155. Then pWHU155 was introduced into the above *genM2* in-frame deletion mutants respectively to generate ΔgenM2::kanE, ΔgenS2ΔgenKΔgenM2::kanE, ΔgenQΔgenKΔgenM2::kanE and ΔgenD1ΔgenKΔgenM2::kanE.

**Construction of ΔgenM2ΔgenQ::kanE, ΔgenM2ΔgenK::kanE and ΔgenM2ΔgenB3::kanE.** pYH286 (for *genQ* in-frame deletion), pWHU1 (for *genK* in-frame deletion) and pWHU5 (for *genB3* in-frame deletion) constructed previously were introduced into ΔgenM2::kanE respectively by conjugation and mutants screening using the method described before[1] to generate ΔgenM2ΔgenQ::kanE, ΔgenM2ΔgenK::kanE and ΔgenM2ΔgenB3::kanE.

All in-frame deletion mutants were verified by PCR using the checking primers (Supplementary Table 2) and Southern blot analysis. Complemented exconjugants were verified based on thiostrepton resistance and confirmed by PCR.

**Small scale production, extraction and LC-ESI-HRMS analysis.**

For small scale aminoglycoside production, extraction and analysis, two-stage lab condition production was used for *M. echinospora* ATCC 15835 and its mutants. The seed culture (in liquid ATCC172 medium) was incubated at 28°C and 220 rpm for 3 days and then used to initiate the 50 mL culture in fermentation medium (F50 medium) with 5% inoculum. For kanamycin feeding experiment 50 μg/mL kanamycin B was supplemented into the culture at this stage. The culture was incubated at 28°C and 220 rpm for 5 days. Each culture broth was adjusted to pH 2.0 with 6 M $H_2SO_4$, agitated for 4 h and then centrifuged at 5,000 g for 10 min at 4°C. The supernatant was filtered and applied to a column of DOWEX 50WX8-200 ion-exchange resin (2.5 g per 50 mL culture broth) preconditioned with acetonitrile followed by Milli-Q water. The column was then washed with Milli-Q water (6 column volume) and aminoglycosides were eluted with 1 M $NH_4OH$ (6 column volume). The eluate was freezedried, dissolved in 1 mL Milli-Q water, filtered through microporous membrane (0.2 μm) and subjected to LC-ESI-HRMS analysis. Each mutant was cultivated and analyzed in triplicate.

LC-ESI-HRMS analysis was performed on Thermo Electron LTQ-Orbitrap XL connected to a Thermo Scientific Accela HPLC system fitted with Phenomenex Luna C18 column (250×4.6 mm) at flow rate 0.4 mL/min using mobile phase of (A) 0.2% trifluoroacetic acid adjusted to pH 2.0 with $NH_4OH$ and (B) 100% acetonitrile. The column was eluted with gradient as: 0-14min 2% B to 6% B, 14-16min 6% B to 8% B, 16-25min 8% B-15% B, 25-26min 15% B to 90% B, 26-34min 90% B, 35-45min 2% B. The mass spectrometry was operated in positive mode with scan range of 300-600 *m/z*. Source conditions were vaporizer temperature at 350°C, capillary temperature at 275°C, voltage of 3.5 kV, sheath gas at 60 L/h, auxiliary gas at 10 L/h. MS/MS analyses were carried out in the positive ionization mode with collision energy of 35 eV.

**Large scale fermentation and mono component HPLC-ELSD purification**

2 L scale lab-condition fermentation was performed similarly as above while distributed in 50 mL across four 250 mL flasks. 30 L scale industrial fermentation was conducted as following: the seed culture (in liquid ATCC172 medium with 25 μg/mL thiostrepton) was incubated at 34°C and 220 rpm for 3 days 48-60 h until the bacterial culture becomes dense and exhibits a slightly reddish color prior inoculating into industrial fermentation medium. The initial fermentation conditions were set as: pH 7.6, rotational speed 245 rpm, airflow rate 2 L/min, temperature 34ºC, internal pressure in the vessel 0.06 MPa. Parameters were monitored during fermentation, maintaining pH stability between 7.2 and 7.6 using 1 M sodium hydroxide and 1 M hydrochloric acid. 2L sterile water was added to the fermentation vessel daily. 3 L medium was added to the fermentation vessel on the third day when the fermentation broth turns red. The fermentation halted on day 5 when fermentation broth turned to a deep purple color.

The fermentation culture was extracted as descripted above with additional purification through 717 cation exchange resin to further remove pigment impurities. Purification of GK mono component from crude extract was performed on evaporative light scattering detector (ELSD, Alltech 2000ES) connected with a Thermo Scientific HPLC (UltiMate 3000) fitted with and a Phenomenex Synergi Hydro-RP 80A (250×10 mm, 4 μm). The column was eluted at a flow rate of 4 mL/min using a mobile phase of (A) 0.2% trifluoroacetic acid (TFA) in $H_2O$ and (B) 100% $CH_3CN$ with a linear gradient of 2%-8% A over 20 minutes. The atomization and drift tube temperature were set at 108°C and gas flow of ELSD was set at 2.8 L/min. Detection and collection ratio was set as 1:3 post-column. Targeted fractions were collected and freeze dried.

**Structural characterization**

**GK-X2** (**3**) was obtained as an amorphous powder and its molecular formula was determined as $C_{20}H_{40}N_4O_{11}$, with three degrees of unsaturation based on its ESI-HRMS data ($[M + H]^+$ *m/z* calcd., 513.2766; found, 513.2782) (Supplementary Fig. 8). The $^1$H NMR spectrum (Supplementary Fig. 9 and Supplementary Table 3) showed signals for two singlet methyl groups ($\delta_H$ 1.42 and 2.93). The $^{13}$C NMR and DEPT spectra (Supplementary Fig.10 and Supplementary Table 3) exhibited 20 carbon signals, including two methyls ($\delta_C$ 20.2 and 34.8), three methylenes ($\delta_C$ 29.2, 59.2 and 60.3), fourteen methines ($\delta_C$ 48.7, 49.4, 54.2, 64.1, 66.1, 69.4, 69.4, 73.5, 73.9, 74.7, 81.9, 84.4, 97.4, and 100.0), and one quaternary carbons ($\delta_C$ 71.3). Without the degrees of unsaturation generated by carboxyl, carbonyl, aldehyde or olefinic groups, the three degrees of unsaturation manifested that **3** had a tricyclic structure. Detailed analysis of 1D and 2D NMR spectra of **3** demonstrated its structure to be similar to aminoglycosides. HMBC correlations (Supplementary Fig. 13) from $H_2$-2 ($\delta_H$ 2.47, 1.83) to C-1 ($\delta_C$ 49.4), from H-5 ($\delta_H$ 3.87) to C-4 ($\delta_C$ 81.9) and C-6 ($\delta_C$ 84.4), in combination with $^1$H–$^1$H COSY correlations (Supplementary Fig. 12) of H-6 ($\delta_H$ 3.80)/H-1 ($\delta_H$ 3.52)/H-2b ($\delta_H$ 1.83)/H-3 ($\delta_H$ 3.44) demonstrated the existence of core aminocyclitol moiety 2-deoxystreptamine, which was decorated by sugar substituents at the C-4 and C-6 positions in **3**. In addition, HMBC correlations (Supplementary Fig.13) from H-1′ ($\delta_H$ 5.62) to C-3′ ($\delta_C$ 69.4) and C-5′ ($\delta_C$ 73.5), from H-2′ ($\delta_H$ 3.40) to C-3′, and from H-5′ ($\delta_H$ 3.87) to C-6′ ($\delta_C$ 60.3), in combination with $^1$H–$^1$H COSY correlations (Supplementary Fig.12) of H-1′/H-2′/H-3′ ($\delta_H$ 3.92) and H-4′ ($\delta_H$ 3.49)/H-5′, and $J_{H1'-H2'}$ = 4.0 Hz (Supplementary Table 3) revealed the existence of *α*-D-glucosamine in **3**. Furthermore, HMBC correlations (Supplementary Fig. 13) from H-1″ ($\delta_H$ 5.21) to C-3″ ($\delta_C$ 64.1) and C-5″ ($\delta_C$ 74.7), from H-2″ ($\delta_H$ 4.22) to C-3″, from $H_2$-6″ ($\delta_H$ 3.97,

3.69) to C-5″, from $H_3$-7″ ($\delta_H$ 1.42) to C-3″, C-4″ ($\delta_C$ 71.3) and C-5″, and from $H_3$-8″ ($\delta_H$ 2.93) to C-3″, in combination with $^1H$–$^1H$ COSY correlations (Supplementary Fig.12) of H-1″/H-2″/H-3″ ($\delta_H$ 3.49) and H-5″ ($\delta_H$ 4.10)/$H_2$-6″, and $J_{H1''\text{-}H2''}$ = 3.9 Hz (Supplementary Table 3) revealed the existence of 3-deoxy-3-(methylamino)-4-methyl-*β*-L-glucopyranose in **3**. The location of *α*-D-glucosamine and 3-deoxy-3-(methylamino)-4-methyl-*β*-L-glucopyranose moieties was deduced to be at C-4 and C-6, respectively, by the key HMBC correlations (Supplementary Fig.13) from H-1′ to C-4 and from H-1″ to C-6. Thus, the structure of **3** was determined as depicted in Supplementary Fig. 7.

**GK-418** (**4**) was obtained as an amorphous powder and its molecular formula was determined as $C_{21}H_{42}N_4O_{11}$, with three degrees of unsaturation based on its ESI-HRMS data ($[M + H]^+$ *m/z* calcd., 527.2923; found, 527.2947) (Supplementary Fig. 16), being 14 mass units greater than that of **3**. The $^1H$ NMR spectrum (Supplementary Fig. 17 and Supplementary Table 4) showed signals for three singlet methyl groups ($\delta_H$ 1.23, 1.43 and 2.94). The $^{13}C$ NMR and DEPT spectra (Supplementary Fig. 18 and Supplementary Table 4) exhibited 21 carbon signals, including three methyls ($\delta_C$ 14.4, 20.2 and 34.8), two methylenes ($\delta_C$ 27.8 and 59.2), fifteen methines ($\delta_C$ 48.9, 49.2, 54.2, 64.0, 65.2, 66.1, 69.4, 70.2, 73.6, 74.7, 75.2, 81.6, 83.6, 97.7 and 100.1), and one quaternary carbons ($\delta_C$ 71.3). Analysis of its 1D and 2D NMR spectra indicated that its structure was closely similar to that of **3**, differing only in a methyl group ($\delta_C$ 14.4) at C-6′ ($\delta_C$ 65.2), verified by HMBC correlations (Supplementary Fig. 21) from H-7′ ($\delta_H$ 1.23) to C-5′ ($\delta_C$ 75.2) and C-6′ ($\delta_C$ 65.2), together with $^1H$–$^1H$ COSY correlations (Supplementary Fig. 20) of H-6′ ($\delta_H$ 3.80)/H-7′. The configuration of C-6′ was assigned to be 6′*R* according to the crucial NOESY correlation (Supplementary Fig. 22) of H-4′/$H_3$-7′ and the knowledge of its biosynthetic pathway. Therefore, the structure of **4** was proposed as shown in Supplementary Fig. 15.

**GK-C1a** (**7**) was obtained as an amorphous powder and its molecular formula was determined as $C_{20}H_{41}N_5O_8$, with three degrees of unsaturation based on its ESI-HRMS data ($[M + H]^+$ *m/z* calcd., 480.3028; found, 480.3049) (Supplementary Fig. 24). The $^1H$ NMR spectrum (Supplementary Fig. 25 and Supplementary Table 5) showed signals for two singlet methyl groups ($\delta_H$ 1.41 and 2.93). $^{13}C$ NMR and DEPT spectra (Supplementary Fig.26 and Supplementary Table 5) exhibited 20 carbon signals, including two methyls ($\delta_C$ 20.2 and 34.8), four methylenes ($\delta_C$ 20.5, 25.6, 28.4 and 59.2), thirteen methines ($\delta_C$ 48.6, 48.7, 49.5, 64.0, 65.9, 66.1, 74.5, 74.7, 77.0, 84.1, 94.7, and 100.0), and one quaternary carbons ($\delta_C$ 71.3). Without degrees of unsaturation generated by carboxyl, carbonyl, aldehyde or olefinic groups, the three degrees of unsaturation manifested that **7** had a tricyclic structure. Analysis of its 1D and 2D NMR spectra indicated that **7** has structural features closely similar to those of **3**, differing only in the *α*-D-glucosamine. HMBC correlations (Supplementary Fig. 29) from H-1′ ($\delta_H$ 5.82) to C-3′ ($\delta_C$ 20.5) and C-5′ ($\delta_C$ 65.9), from H-2′ ($\delta_H$ 3.55) to C-3′, from H-3′a ($\delta_H$ 2.03) to C-4′ ($\delta_C$ 25.6), and from $H_2$-6′b ($\delta_H$ 3.07) to C-4′ and C-5′, in combination with $^1$H–$^1$H COSY correlations (Supplementary Fig. 28) of H-1′/H-2′/H-3′a/H-4′b ($\delta_H$ 1.58)/H-5′ ($\delta_H$ 4.18) /H-6′b revealed the change of *α*-D-glucosamine in **7**. Thus, the structure of **7** could be determined as depicted in Supplementary Fig. 23.

**GK-C2a** (**8**) was obtained as an amorphous powder and its molecular formula was determined as $C_{21}H_{43}N_5O_8$, with three degrees of unsaturation based on its ESI-HRMS data ($[M + H]^+$ *m/z* calcd., 494.3184; found, 494.3206) (Supplementary Fig. 32), being 14 mass units greater than that of **7**. The $^1$H NMR spectrum (Supplementary Fig. 33 and Supplementary Table 6) showed signals for three singlet methyl groups ($\delta_H$ 1.33, 1.42 and 2.94). $^{13}$C NMR and DEPT spectra (Supplementary Fig. 34 and Supplementary Table 6) exhibited 21 carbon signals, including three methyls ($\delta_C$ 14.1, 20.2 and 34.8), four methylenes ($\delta_C$ 20.3, 25.3, 27.6 and 59.2), thirteen methines ($\delta_C$ 48.6, 48.7, 49.4, 51.1, 64.0, 66.1, 70.1, 74.5, 74.7, 75.7, 83.9, 94.2 and 100.1), and one quaternary carbons ($\delta_C$ 71.3). Analysis of its 1D and 2D NMR spectra indicated that **8** has structural features closely similar to those of **7**, differing only in a methyl group ($\delta_C$ 14.1) at C-6′ ($\delta_C$ 51.1) proved by the HMBC correlation (Supplementary Fig. 37) from H-7′ ($\delta_H$ 1.33) to C-5′ ($\delta_C$ 70.1) and C-6′ ($\delta_C$ 51.1), as well as the $^1$H–$^1$H COSY correlations (Supplementary Fig. 36) of H-5′ ($\delta_H$ 3.87)/H-6′ ($\delta_H$ 3.35)/H-7′. The configuration of C-6′ was assigned to be 6′*S* according to the crucial NOESY correlation (Supplementary Fig. 38) of H-4′/H-6′ and the knowledge of its biosynthetic pathway. Accordingly, the structure of **8** was proposed as shown in Supplementary Fig. 31.

**GK-C2** (**9**) was obtained as an amorphous powder and its molecular formula was determined as $C_{21}H_{43}N_5O_8$, with three degrees of unsaturation based on its ESI-HRMS data ($[M + H]^+$ *m/z* calcd., 494.3184; found, 494.3201) (Supplementary Fig. 40). It had the same molecular formula as that of **8**. The $^1$H NMR spectrum (Supplementary Fig. 41 and Supplementary Table 7) showed signals for three singlet methyl groups ($\delta_H$ 1.30, 1.42 and 2.94). $^{13}$C NMR and DEPT spectra (Supplementary Fig. 42 and Supplementary Table 7) exhibited 21 carbon signals, including three methyls ($\delta_C$ 12.2, 20.2 and 34.8), four methylenes ($\delta_C$ 20.6, 23.0, 27.7 and 59.2), thirteen methines ($\delta_C$ 48.6, 48.8, 49.4, 49.5, 64.0, 66.1, 68.8, 74.4, 74.7, 76.3, 83.9, 94.9 and 100.1), and one quaternary carbons ($\delta_C$ 71.3). Comparative analysis of their 1D and 2D NMR spectra indicated that **9** and **8** were a pair of C-6′ epimers. The configuration of C-6′ was assigned to be 6′*R* according to the crucial NOESY correlation (Supplementary Fig. 46) of H-4′/$H_3$-7′ and the knowledge of its biosynthetic pathway. Thus, the structure of **9** was assigned as shown Supplementary Fig. 39.

**GK-C1** (**11**) was obtained as an amorphous powder and its molecular formula was determined as $C_{22}H_{45}N_5O_8$, with three degrees of unsaturation based on its ESI-HRMS data ([M + H]$^+$ *m/z* calcd., 508.3340; found, 508.3361) (Supplementary Fig. 48), being 14 mass units greater than that of **9**. The $^1$H NMR spectrum (Supplementary Fig. S49 and Supplementary Table 8) showed signals for four singlet methyl groups ($\delta_H$ 1.31, 1.42, 2.76 and 2.94). $^{13}$C NMR and DEPT spectra (Supplementary Fig. 50 and Supplementary Table 8) exhibited 22 carbon signals, including four methyls ($\delta_C$ 9.4, 20.2, 31.1 and 34.8), four methylenes ($\delta_C$ 20.6, 23.1, 27.7 and 59.2), thirteen methines ($\delta_C$ 48.6, 48.7, 49.4, 57.5, 64.0, 66.1, 69.1, 74.4, 74.7, 76.1, 83.9, 94.8 and 100.1), and one quaternary carbons ($\delta_C$ 71.3). Analysis of its 1D and 2D NMR spectra indicated that **11** has structural features closely similar to those of **9**, differing only in a methyl group ($\delta_C$ 31.1) at *N* atom of C-6′, revealed by crucial HMBC correlation (Supplementary Fig. 53) from H-8′ ($\delta_H$ 2.76) to C-6′ ($\delta_C$ 57.5). The configuration of C-6′ was assigned to be 6′*R* according to the crucial NOESY correlation (Supplementary Fig. 54) of H-4′/$H_3$-7′ and the knowledge of its biosynthetic pathway. Thus, the structure of **11** was proposed as shown in Supplementary Fig. 47.

#### GKs sulfate preparation

TFA was removed from the HPLC-ELSD purified samples by multiple time freeze drying and then products were finally dissolved in water and the pH of the solution was adjusted to neutral using $H_2SO_4$ to give GKs sulfate that used for test of antimicrobial activity and toxicity.

**Antimicrobial activity test**

Antibacterial activity of the aminoglycosides was evaluated by minimal inhibition concentration (MIC) against indicator strains. MIC value was determined using broth microdilution method according to standard protocols of antimicrobial susceptibility test (Clinical and Laboratory Standards Institute, CLSI, 2014). The inoculum was prepared by resuspending the isolated colonies of indicator strains from an 18-24 h growth on blood agar plate in Mueller Hinton Broth (MHB) (Qingdao Hope Bio-Technology Co., Ltd) for *Enterococcus faecium* ATCC 19434, *Staphylococcus aureus* ATCC 29213, *Klebsiella pneumoniae* ATCC 700603, *Acinetobacter baumannii* ATCC 19606, and *Enterobacter cloacae* ATCC 13047 or cation-adjusted MHB (CAMHB, with final concentration of $Mg^{2+}$ and $Ca^{2+}$ adjusted to 12.5 mg/L and 25 mg/L with $MgCl_2$ and $CaCl_2$, respectively) for *Pseudomonas aeruginosa* ATCC 27853, and adjusting the suspension to achieve a turbidity equivalent to a 0.5 McFarland standard with MHB or CAMHB. This inoculum was then diluted 100 times with MHB or CAMHB to reach the concentration of approximately $1\times10^6$ CFU/mL just before the assay. A 2-fold serial dilution of each aminoglycoside sulfate solution stock (100 μL) in MHB or CAMHB was prepared in 96-well microplates except the last column, which served as negative controls (bacterial inoculum and MHB without aminoglycoside). 100 μL of prepared bacterial suspensions were added to each well to reach a final concentration of approximately $5\times10^5$ CFU/mL. After 20-24 h of incubation at 37°C, MIC was determined as the lowest concentration of the aminoglycosides inhibiting visible bacterial growth. All experiments were performed in triplicates.

**Toxicity characterization**

**Zebrafish strains and maintenance.** Zebrafish (*Danio rerio*) adults and embryos were raised and maintained in the Zebrafish Center of Nantong University under conditions described as our previous protocols[4]. The transgenic line *Tg(Brn3c:mGFP)*[5,6] and wild-type AB strain[7] were used in ototoxicity test, in which, the membrane-localized green fluorescent protein (GFP) is expressed specifically in the hair cells (HCs). 4-5 days post fertilization (dpf) larvae were used in this study. Animal Care and Use Committee of Nantong University and Wuhan University approved all animal procedures.

**Drugs treatment and microinjection in zebrafish.** Compounds of GK-C1, C1, GK-C1a, C1a, GK-C2, C2, GK-C2a, C2a, kanamycin B and dibekacin were directly diluted in E3 embryo medium[8] to prepare the working solutions with appropriate concentrations. For assessing compounds toxicity to HCs in poster lateral line (pLL), ten healthy *Tg(Brn3c:mGFP)* zebrafish larvae at 5 dpf/well were placed in a 24-well plate and exposed to aminoglycoside compounds at different concentrations (0.5, 1, 2.5, 5, 10, 25, 50, 100 μM). Compound solutions were removed after 6 h and larvae were washed three times with embryo medium for subsequently imaging or startle response assay. To evaluate the toxicity of drugs to HCs in otic vesicle, Texas Red labeled dextran (Invitrogen, United States) was added to prepare drug solution with a final concentration of 10 mg/mL. About 2 nL mixture was microinjected into semicircular canal of larvae at 5 dpf, and lasting for 6 h to observe the phenotypes of crista HCs in otic vesicle.

**Vital dye staining.** The vital dye FM® 4-64 (n-(3-triethylammoniumpropyl)-4-(6-(4-(diethylamino) phenyl) hexatrienyl) pyridinium dibromide, Invitrogen Molecular Probes, Eugene, OR) was used to specifically label functional HCs in neuromasts of pL[9]. Free swimming larvae were immersed in 3 μM FM® 4-64 in embryo medium for 45 s at room temperature in the dark, followed by three rinses in E3 embryo medium.

**Startle response test.** Twenty normal larvae were put in a thin layer of E3 embryo medium in a petri dish attached to a mini vibrator which generated sound stimulus (a tone burst 9 dB re $ms^{-2}$, 600 Hz, for 30 ms). Twenty times stimuli were repeated and the response of larvae to every sound stimulus was recorded by an infrared camera over a 6 s period. Subsequently, movement trajectories were extracted from the recording movies, and the movement typical parameters of mean distance and peak velocity were used to quantify the startle response of larvae to sound stimuli.

**Images acquisition and statistical analysis.** The HCs phenotypes in our experiments were scanned by a confocal microscopy (Nikon, A1-DUT). For microscopic imaging of zebrafish, embryos were anaesthetized with MS-222 (Sigma) and embedded in 0.6% low-melting point agarose. Confocal images analysis was performed by Imaris X64 software (version 9.0.1). All data were presented as mean ± standard error of the mean (SEM), and all experiments were repeated at least three times. Two-tailed, unpaired student's t-test was used to identify the significance difference between groups, with $P < 0.05$ considered statistically significant.

#### Supplementary Tables

**Supplementary Table 1.** The bacterial strains and plasmids used in this study.

| Strains/Plasmids | Description | Reference |
|---|---|---|
| ***Escherichia coil*** | | |
| DH10B | Host for general cloning | Invitrogen |
| ET12567/pUZ8002 | Donor strain for conjugation between *E. coli* and *Streptomyces* | 10 |
| ***Micromonospora echinospora*** | | |
| ATCC 15835 | Gentamicin-producing wild-type strain | 11 |
| ΔgenM2 | Gentamicin production abolished mutant | This work |
| ΔgenS2ΔgenK | Gentamicin A2 accumulating mutant | 12 |
| ΔgenD1ΔgenK | Gentamicin A accumulating mutant | 3 |
| ΔgenQΔgenK | Gentamicin X2 accumulating mutant | 1 |
| ΔgenQ | G418 accumulating mutant | 1 |
| ΔgenB3 | JI-20B accumulating mutant | 1 |
| ΔgenK | Gentamicins “left-hand” branch products accumulating mutant | 1 |
| ΔgenS2ΔgenKΔgenM2 | Gentamicin production abolished mutant | This work |
| ΔgenD1ΔgenKΔgenM2 | Gentamicin production abolished mutant | This work |
| ΔgenQΔgenKΔgenM2 | Gentamicin production abolished mutant | This work |
| ΔgenM2::kanE | GKs producing mutant | This work |
| ΔgenS2ΔgenKΔgenM2::kanE | GK-A2 accumulating mutant | This work |
| ΔgenD1ΔgenKΔgenM2::kanE | GK-A accumulating mutant | This work |
| ΔgenQΔgenKΔgenM2::kanE | GK-X2 accumulating mutant | This work |
| ΔgenM2ΔgenQ::kanE | GK-418 accumulating mutant | This work |
| ΔgenM2ΔgenB3::kanE | GK-JI-20B accumulating mutant | This work |

| | | |
|---|---|---|
| ΔgenM2ΔgenK::kanE | GKs "left-hand" branch products accumulating mutant | This work |
| ***Streptomyces kanamyceticus*** | | |
| ATCC 12853 | Kanamycin-producing wild-type strain | ATCC |
| **Antimicrobial activity indicator strains** | | |
| *Enterococcus Faecium* ATCC 19434 | Standard pathogenic bacteria strain | ATCC |
| *Staphylococcus aureus* ATCC 29213 | Standard pathogenic bacteria strain | ATCC |
| *Acinetobacter baumannii* ATCC 19606 | Standard pathogenic bacteria strain | ATCC |
| *Pseudomonas aeruginosa* ATCC 27853 | Standard pathogenic bacteria strain | ATCC |
| *Klebsiella pneumoniae* ATCC 700603 | Standard pathogenic bacteria strain | ATCC |
| *Enterobacter Cloacae* ATCC 13047 | Standard pathogenic bacteria strain | ATCC |
| **Plasmids** | | |
| pYH7 | *E. coli - Streptomyces* shuttle vector | 2 |
| pWHU47 | Construct for *genM2* in-frame deletion | This work |
| pWHU2649 | Construct for *genM2* in-frame deletion based on ΔgenD1ΔgenK | This work |
| pYH286 | Construct for *genQ* in-frame deletion | 1 |
| pWHU1 | Construct for *genK* in-frame deletion | 1 |
| pWHU5 | Construct for *genB3* in-frame deletion | 1 |
| pWHU77 | Vector for complementation | 1 |
| pWHU155 | Construct for *kanE* complementation | This work |

**Supplementary Table 2.** Oligonucleotide primers used in this study.

| **Primer** | **Oligonucleotide sequences (5' to 3')** | **Restriction site** |
|---|---|---|
| kanE-F | AGGG**CATATG**ACCGAGCCTGCCAAG | *Nde*I |
| kanE-R | AGACGAG**GAATTC**CCTCACAGCCCG | *Eco*RI |
| genM2-L-F | GGT**CATATG**GACCCGTGACGAAGGG | *Nde*I |
| genM2-L-R | GCC**GAATTC**GTGCATGCCGCCCATC | *Eco*RI |
| genM2-R-F | ACC**GAATTC**CTGGTGCCGCCCTTCG | *Eco*RI |
| genM2-R-R | CCG**AAGCTT**GCCAGTTTCTGCGTCA | *Hin*dIII |
| genM2 in ΔgenD1ΔgenK-L-F | GTG**CATATG**GGTTGTGCGTATGGAG | *Nde*I |
| genM2 in ΔgenD1ΔgenK-L-R | CCC**ACTAGT**GTGCATGCCGCCCATCG | *Spe*I |
| genM2 in ΔgenD1ΔgenK-R-F | ACC**ACTAG**TCTGGTGCCGCCCTTCG | *Spe*I |
| genM2 in ΔgenD1ΔgenK-R-R | TGA**AAGCTT**AGCGGAAGCTGGACTT | *Hin*dIII |
| kanE-CK-F | GATGTGCTGCAAGGCGATTAAGTTGGG | |
| kanE-CK-R | GCTTTACACTTTATGCTTCCGGCTCGTATGTT | |
| genM2-CK-F | CCCCGAGTGGGAGTGGAT | |
| genM2-CK-R | TGATGAAGGGCTTGGTGAA | |
| genM2 in ΔgenD1ΔgenK-CK-F | TCCGCAACAGCGGATACGTGCAC | |
| genM2 in ΔgenD1ΔgenK-CK-R | ACAACGACGGTGGGAATGCGACA | |
| genQ-CK-F | CCTCCTCGTCACCGTGG | |
| genQ-CK-R | CAGGTGCTCAGCGTCCG | |
| genB3-CK-F | CGCGTTACGGAAAGTAAAATCAC | |
| genB3-CK-R | CATCGAGGGCCACCACC | |
| genK-CK-F | CGGGCGAACCTTCGGGATA | |
| genK-CK-R | CCGTCAGCGTTGGCAATAA | |

**Supplementary Table 3.** $^{1}H$ NMR (600 MHz, $D_2O$) and $^{13}C$ NMR (150 MHz, $D_2O$) data of GK-X2 (**3**).

| **No.** | $\delta_C$ | $\delta_H$ |
|---|---|---|
| 1 | 49.4, CH | 3.52, m |
| 2a | 29.2, $CH_2$ | 2.47, dt (12.8, 4.4) |
| 2b | | 1.83, q (12.8) |
| 3 | 48.7, CH | 3.44, m |
| 4 | 81.9, CH | 3.81, overlap |
| 5 | 73.9, CH | 3.87, overlap |
| 6 | 84.4, CH | 3.80, overlap |
| 1′ | 97.4, CH | 5.62, d (4.0) |
| 2′ | 54.2, CH | 3.40, ddd (10.8, 4.0, 1.5) |
| 3′ | 69.4, CH | 3.92, overlap |
| 4′ | 69.4, CH | 3.49, overlap |
| 5′ | 73.5, CH | 3.87, overlap |
| 6′a | 60.3, $CH_2$ | 3.92, overlap |
| 6′b | | 3.76, m |
| 1″ | 100.0, CH | 5.21, d (3.9) |
| 2″ | 66.1, CH | 4.22, ddd (11.1, 3.9, 1.2) |
| 3″ | 64.1, CH | 3.49, overlap |
| 4″ | 71.3, C | |
| 5″ | 74.7, CH | 4.10, dd (8.3, 2.8) |
| 6″a | 59.2, $CH_2$ | 3.97, dd (12.0, 2.8) |
| 6″b | | 3.69, ddd (12.0, 8.5, 1.4) |
| 7″ | 20.2, $CH_3$ | 1.42, s |
| 8″ | 34.8, $CH_3$ | 2.93, s |

**Supplementary Table 4.** $^{1}H$ NMR (600 MHz, $D_2O$) and $^{13}C$ NMR (150 MHz, $D_2O$) data of GK-418 (**4**).

| No. | $\delta_C$ | $\delta_H$ |
|---|---|---|
| 1 | 49.2, CH | 3.64, overlap |
| 2a | 27.8, $CH_2$ | 2.60, d (12.2) |
| 2b | | 2.01, overlap |
| 3 | 48.9, CH | 3.64, overlap |
| 4 | 81.6, CH | 3.96, overlap |
| 5 | 73.6, CH | 3.93, overlap |
| 6 | 83.6, CH | 3.92, overlap |
| 1′ | 97.7, CH | 5.62, t (3.4) |
| 2′ | 54.2, CH | 3.48, dd (10.9, 4.0) |
| 3′ | 69.4, CH | 3.95, overlap |
| 4′ | 70.2, CH | 3.45, t (9.6) |
| 5′ | 75.2, CH | 3.94, overlap |
| 6′ | 65.2, CH | 4.25, overlap |
| 7′ | 14.4, $CH_3$ | 1.23, d (6.6) |
| 1″ | 100.1, CH | 5.24, d (3.8) |
| 2″ | 66.1, CH | 4.25, overlap |
| 3″ | 64.0, CH | 3.52, d (11.0) |
| 4″ | 71.3, C | |
| 5″ | 74.7, CH | 4.12, dd (8.6, 2.6) |
| 6″a | 59.2, $CH_2$ | 3.99, dd (12.0, 2.6) |
| 6″b | | 3.70, dd (12.0, 8.6) |
| 7″ | 20.2, $CH_3$ | 1.43, s |
| 8″ | 34.8, $CH_3$ | 2.94, s |

**Supplementary Table 5.** $^{1}H$ NMR (600 MHz, $D_2O$) and $^{13}C$ NMR (150 MHz, $D_2O$) data of GK-C1a (**7**).

| No. | $\delta_C$ | $\delta_H$ |
|---|---|---|
| 1 | 49.5, CH | 3.59, m |
| 2a | 28.4, $CH_2$ | 2.52, dt (12.6, 4.4) |
| 2b | | 2.03, overlap |
| 3 | 48.6, CH | 3.52, m |
| 4 | 77.0, CH | 4.03, m |
| 5 | 74.5, CH | 3.91, t (8.9) |
| 6 | 84.1, CH | 3.87, t (9.5) |
| 1′ | 94.7, CH | 5.82, d (3.6) |
| 2′ | 48.7, CH | 3.55, m |
| 3′a | 20.5, $CH_2$ | 2.03, overlap |
| 3′b | | 2.03, overlap |
| 4′a | 25.6, $CH_2$ | 1.94, dd (13.9, 3.4) |
| 4′b | | 1.58, m |
| 5′ | 65.9, CH | 4.18, m |
| 6′a | 42.7, $CH_2$ | 3.25, dd (13.4, 3.0) |
| 6′b | | 3.07, dd (13.4, 8.5) |
| 1″ | 100.0, CH | 5.22, d (4.0) |
| 2″ | 66.1, CH | 4.23, dd (11.1, 4.0) |
| 3″ | 64.0, CH | 3.50, overlap |
| 4″ | 71.3, C | |
| 5″ | 74.7, CH | 4.10, dd (8.5, 2.8) |
| 6″a | 59.2, $CH_2$ | 3.97, dd (11.9, 2.8) |
| 6″b | | 3.68, dd (11.9, 8.5) |
| 7″ | 20.2, $CH_3$ | 1.41, s |
| 8″ | 34.8, $CH_3$ | 2.93, s |

**Supplementary Table 6.** $^{1}H$ NMR (600 MHz, $D_2O$) and $^{13}C$ NMR (150 MHz, $D_2O$) data of GK-C2a (**8**).

| **No.** | $\delta_C$ | $\delta_H$ |
|---|---|---|
| 1 | 49.4, CH | 3.64, m |
| 2a | 27.6, $CH_2$ | 2.57, dt (12.6, 4.4) |
| 2b | | 2.15, q (12.6) |
| 3 | 48.7, CH | 3.60, m |
| 4 | 75.7, CH | 4.13, m |
| 5 | 74.5, CH | 3.95, d (8.9) |
| 6 | 83.9, CH | 3.92, m |
| 1′ | 94.2, CH | 5.93, d (3.6) |
| 2′ | 48.6, CH | 3.56, m |
| 3′a | 20.3, $CH_2$ | 2.03, overlap |
| 3′b | | 2.04, overlap |
| 4′a | 25.3, $CH_2$ | 2.06, overlap |
| 4′b | | 1.54, dt (14.0, 9.6) |
| 5′ | 70.1, CH | 3.87, m |
| 6′ | 51.1, CH | 3.35, m |
| 7′ | 14.1, $CH_3$ | 1.33, d (6.8) |
| 1″ | 100.1, CH | 5.23, d (3.9) |
| 2″ | 66.1, CH | 4.24, dd (11.1, 3.9) |
| 3″ | 64.0, CH | 3.51, d (11.1) |
| 4″ | 71.3, C | |
| 5″ | 74.7, CH | 4.10, m |
| 6″a | 59.2, $CH_2$ | 3.98, m |
| 6″b | | 3.68, dd (12.0, 8.5) |
| 7″ | 20.2, $CH_3$ | 1.42, s |
| 8″ | 34.8, $CH_3$ | 2.94, s |

**Supplementary Table 7.** $^{1}H$ NMR (600 MHz, $D_2O$) and $^{13}C$ NMR (150 MHz, $D_2O$) data of GK-C2 (**9**).

| No. | $\delta_C$ | $\delta_H$ |
|---|---|---|
| 1 | 49.4, CH | 3.63, overlap |
| 2a | 27.7, $CH_2$ | 2.59, dt (12.6, 4.4) |
| 2b | | 2.11, m |
| 3 | 48.8, CH | 3.55, overlap |
| 4 | 76.3, CH | 4.07, dd (10.3, 8.9) |
| 5 | 74.4, CH | 3.96, m |
| 6 | 83.9, CH | 3.91, t (9.5) |
| 1′ | 94.9, CH | 5.89, d (3.7) |
| 2′ | 48.6, CH | 3.55, overlap |
| 3′a | 20.6, $CH_2$ | 2.06, overlap |
| 3′b | | 2.06, overlap |
| 4′a | 23.0, $CH_2$ | 1.91, dd (14.0, 3.1) |
| 4′b | | 1.64, m |
| 5′ | 68.8, CH | 4.14, m |
| 6′ | 49.5, CH | 3.63, overlap |
| 7′ | 12.2, $CH_3$ | 1.30, d (6.9) |
| 1″ | 100.1, CH | 5.23, d (3.9) |
| 2″ | 66.1, CH | 4.24, dd (11.0, 3.9) |
| 3″ | 64.0, CH | 3.52, d (11.0) |
| 4″ | 71.3, C | |
| 5″ | 74.7, CH | 4.11, dd (8.5, 2.6) |
| 6″a | 59.2, $CH_2$ | 3.98, m |
| 6″b | | 3.69, overlap |
| 7″ | 20.2, $CH_3$ | 1.42, s |
| 8″ | 34.8, $CH_3$ | 2.94, s |

**Supplementary Table 8.** $^{1}H$ NMR (600 MHz, $D_2O$) and $^{13}C$ NMR (150 MHz, $D_2O$) data of GK-C1 (**11**).

| No. | $\delta_C$ | $\delta_H$ |
|---|---|---|
| 1 | 49.4, CH | 3.62, m |
| 2a | 27.7, $CH_2$ | 2.58, dt (12.7, 4.5) |
| 2b | | 2.11, q (12.7) |
| 3 | 48.7, CH | 3.57, m |
| 4 | 76.1, CH | 4.08, d (9.8) |
| 5 | 74.4, CH | 3.95, m |
| 6 | 83.9, CH | 3.90, t (9.5) |
| 1′ | 94.8, CH | 5.91, d (3.7) |
| 2′ | 48.6, CH | 3.54, m |
| 3′a | 20.6, $CH_2$ | 2.05, overlap |
| 3′b | | 2.05, overlap |
| 4′a | 23.1, $CH_2$ | 1.92, dd (13.8, 3.2) |
| 4′b | | 1.63, m |
| 5′ | 69.1, CH | 4.18, dt (12.2, 2.8) |
| 6′ | 57.5, CH | 3.46, qd (6.8, 2.8) |
| 7′ | 9.4, $CH_3$ | 1.31, d (6.8) |
| 8′ | 31.1, $CH_3$ | 2.76, s |
| 1″ | 100.1, CH | 5.23, d (4.0) |
| 2″ | 66.1, CH | 4.24, dd (11.0, 4.0) |
| 3″ | 64.0, CH | 3.51, d (11.0) |
| 4″ | 71.3, C | |
| 5″ | 74.7, CH | 4.10, dd (8.5, 2.9) |
| 6″a | 59.2, $CH_2$ | 3.98, m |
| 6″b | | 3.68, dd (11.9, 8.5) |
| 7″ | 20.2, $CH_3$ | 1.42, s |
| 8″ | 34.8, $CH_3$ | 2.94, s |

#### Supplementary Figures

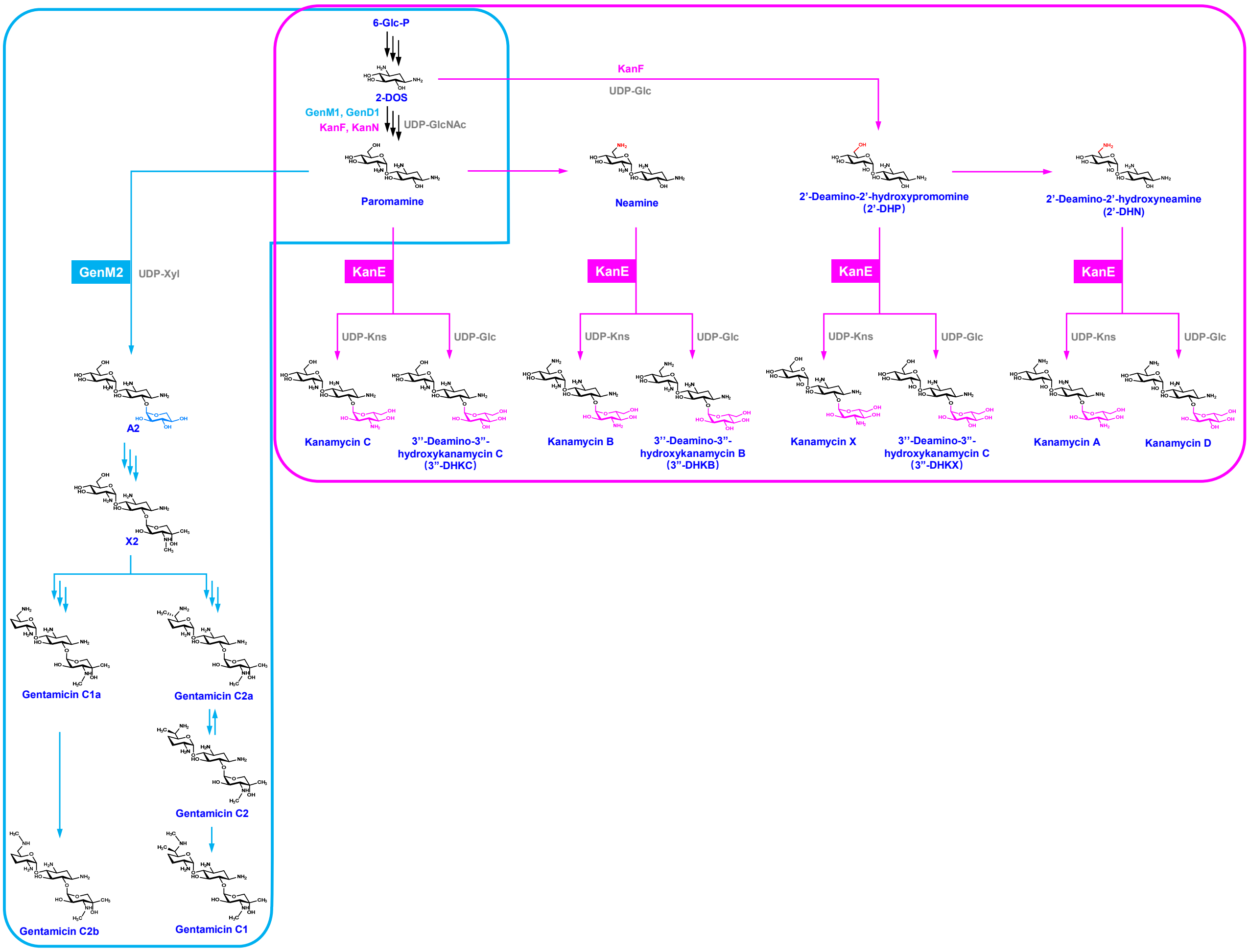

**Supplementary Fig. 1** Comprehensive biosynthetic pathway of gentamicin and kanamycin complex. The gentamicin and kanamycin biosynthetic pathway and their involved glycotransferase are presented in the blue and red, respectively. The shared pathway is indicated by black arrows. 6-Glc-P, 6-Phosphate-D-glucose; 2-DOS, 2-Deoxystreptamine; UDP-GlcNAc, Uridine 5'-diphospho-D-2-N-acetylglucosamine; UDP-Xyl, UDP-xylose; UDP-Kns, UDP-kanosamine; UDP-Glc, UDP-D-glucose; 2'-DHP, 2'-Diamino-2'-hydroxypromomine; 2'-DHN, 2'-Diamino-2'-hydroxyneamine; 3"-DHKC, 3''-Diamino-3"-hydroxykanamycin C; 3"-DHKB, 3''-Diamino-3"-hydroxykanamycin B; 3"-DHKX, 3''-Diamino-3"-hydroxykanamycin C.

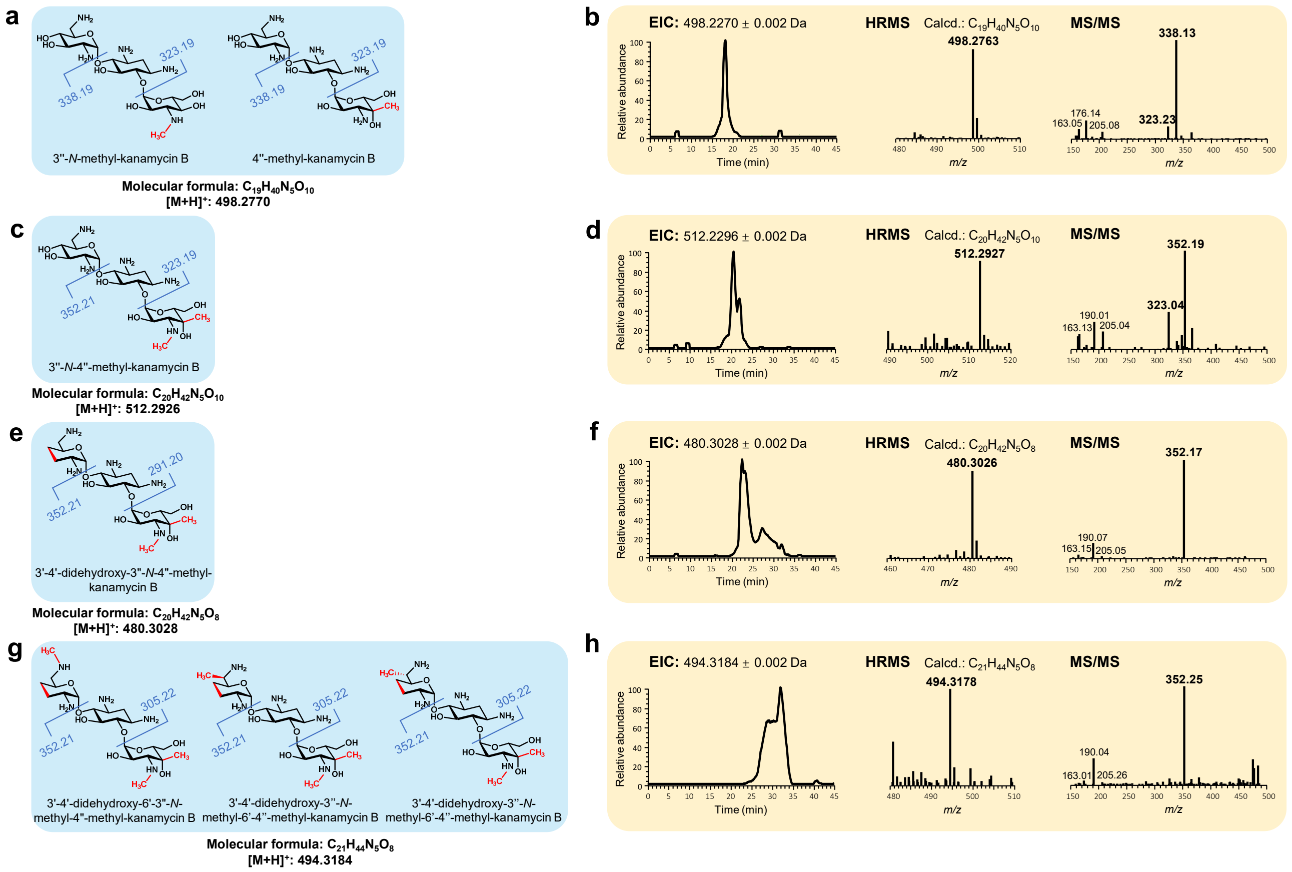

**Supplementary Fig. 2** Predicted structures (**a**, **c**, **e**, **g**) and their observed ions (**b**, **d**, **f**, **h**) by LC-ESI-HRMS and MS/MS analysis of fermentation extract of ΔgenM2 fed with kanamycin B. Structural differences compared to kanamycin B are highlighted in red.

(1) ΔgenM2

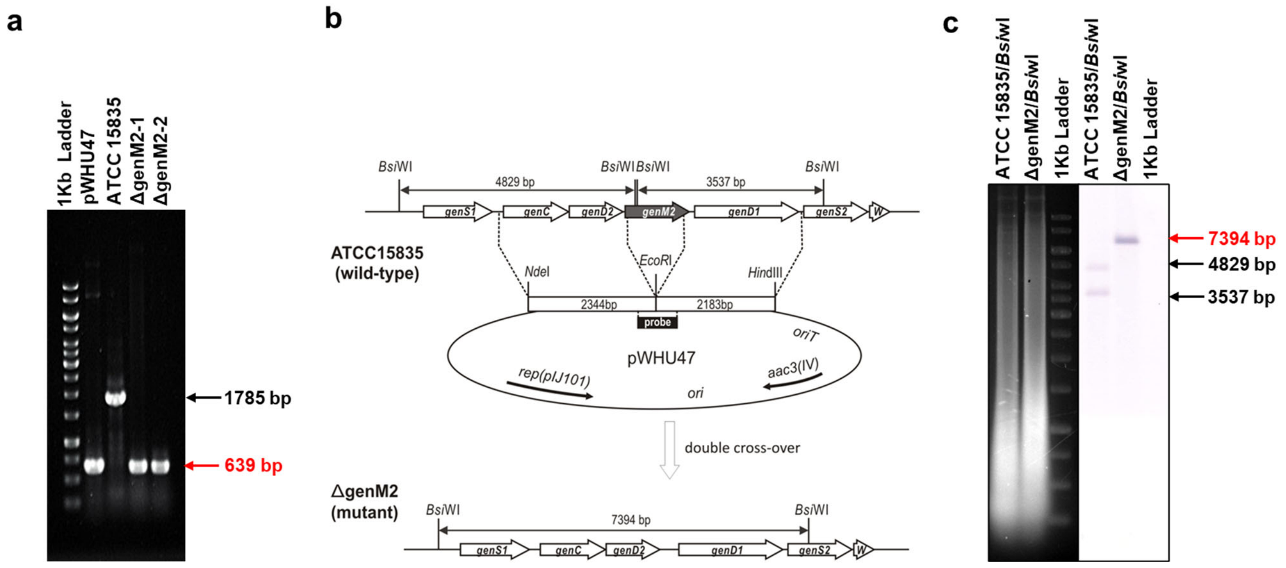

(2) ΔgenM2ΔgenK::kanE

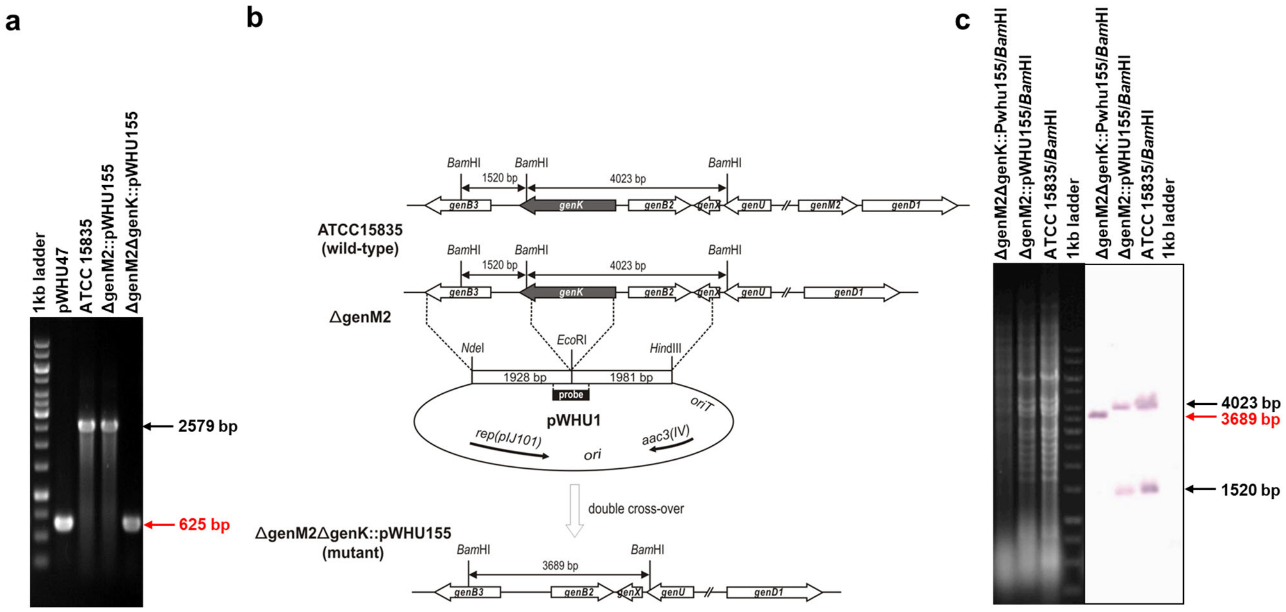

**Supplementary Fig. 3** Genetic confirmation of in-frame gene deletion and *kanE* complementation mutants. PCR confirmation (**a**) schematic representation (**b**) and Southern blot confirmation (**c**) of in-frame gene deletion mutants.

(3) ΔgenS2ΔgenKΔgenM2

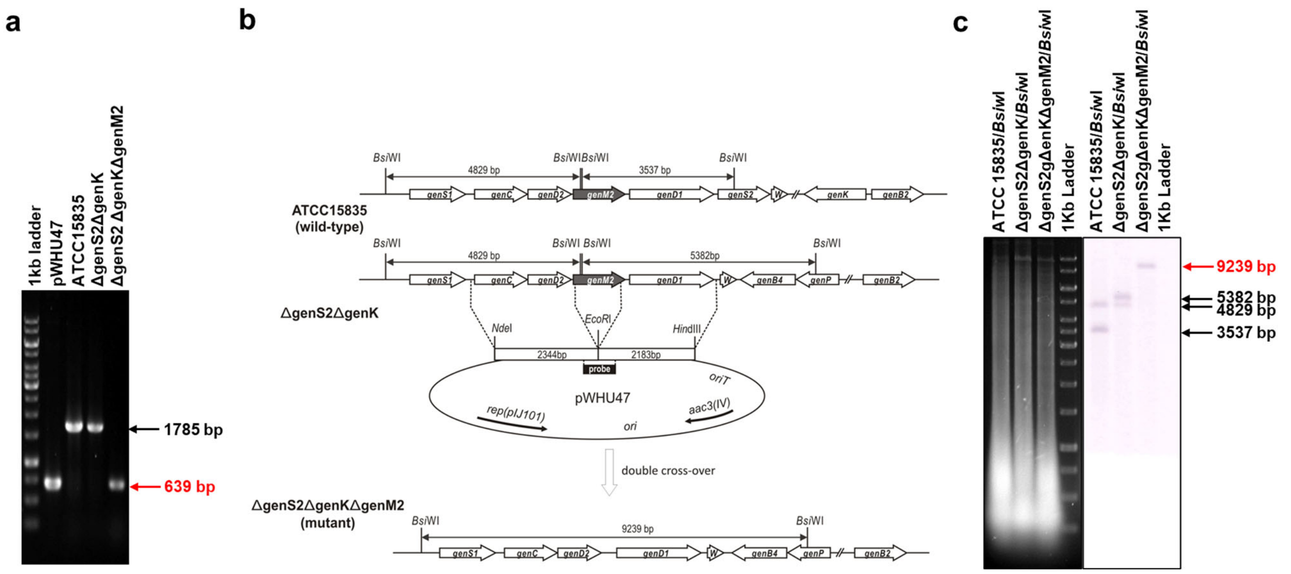

(4) ΔgenD1ΔgenKΔgenM2

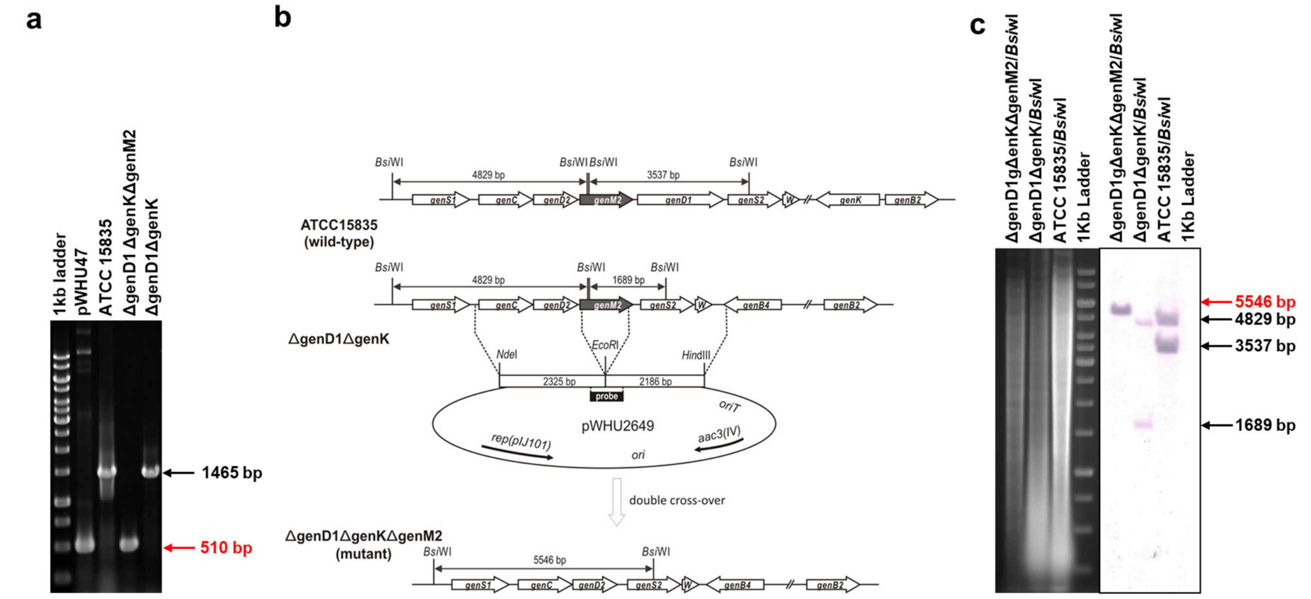

**Supplementary Fig. 3 (continued)** Genetic confirmation of in-frame gene deletion and *kanE* complementation mutants. PCR confirmation (**a**), schematic representation (**b**) and Southern blot confirmation (**c**) of in-frame gene deletion mutants.

(5) ΔgenQΔgenKΔgenM2

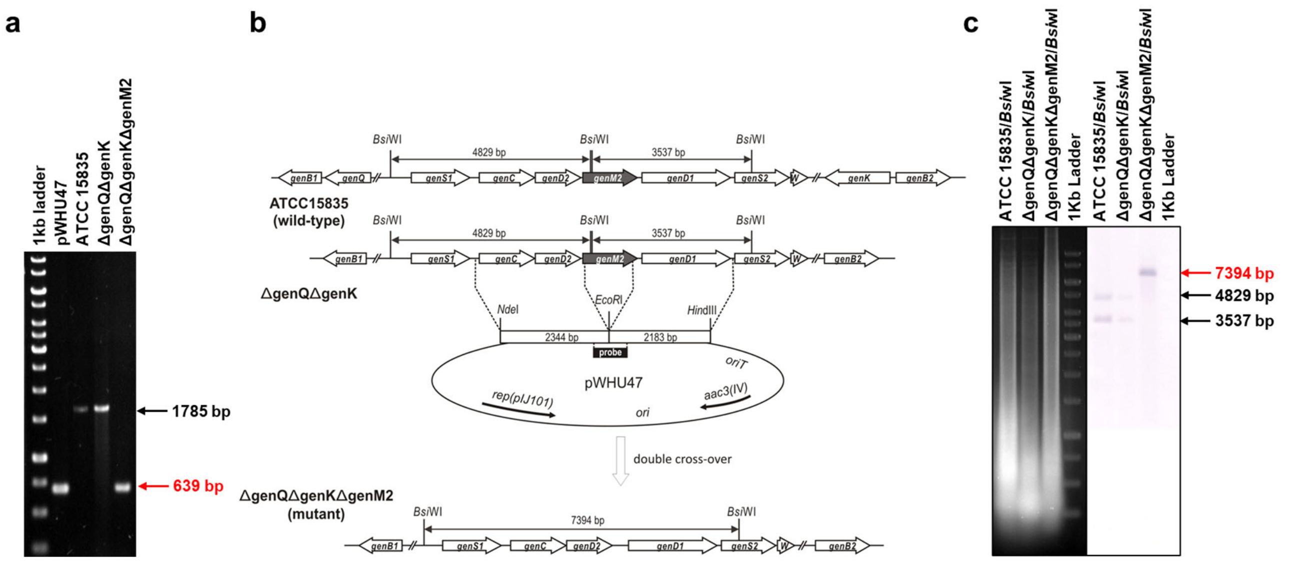

(6) ΔgenM2ΔgenQ::kanE

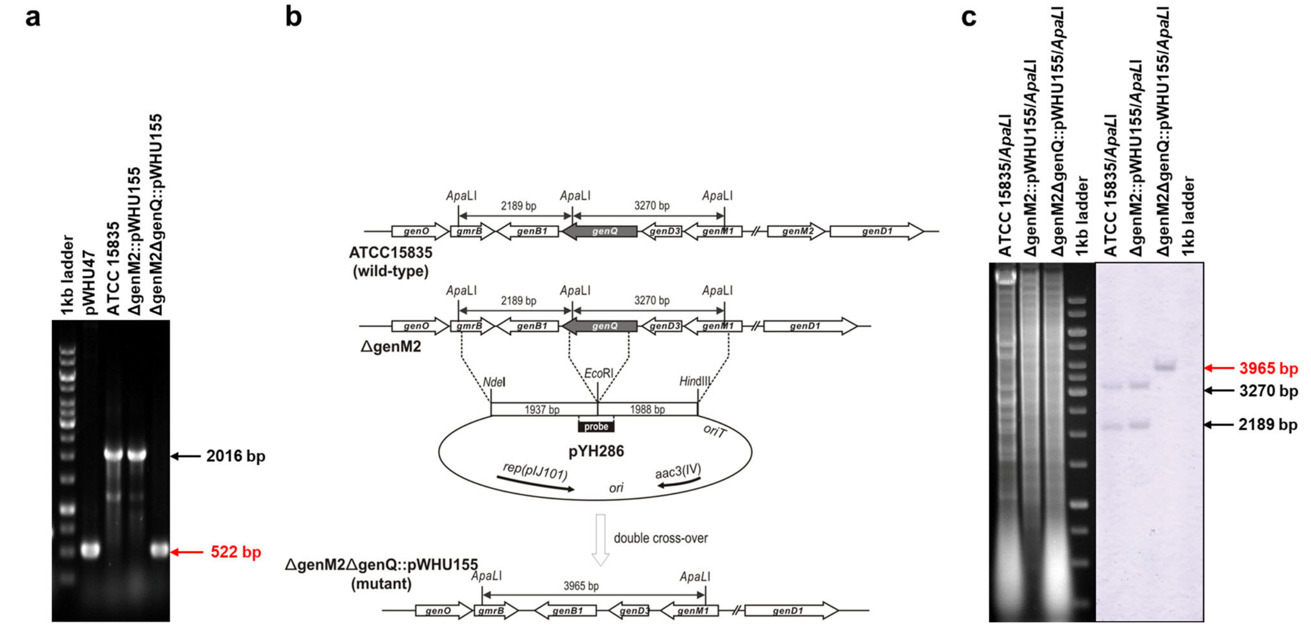

**Supplementary Fig. 3 (continued)** Genetic confirmation of in-frame gene deletion and *kanE* complementation mutants. PCR confirmation (**a**), schematic representation (**b**) and Southern blot confirmation (**c**) of in-frame gene deletion mutants.

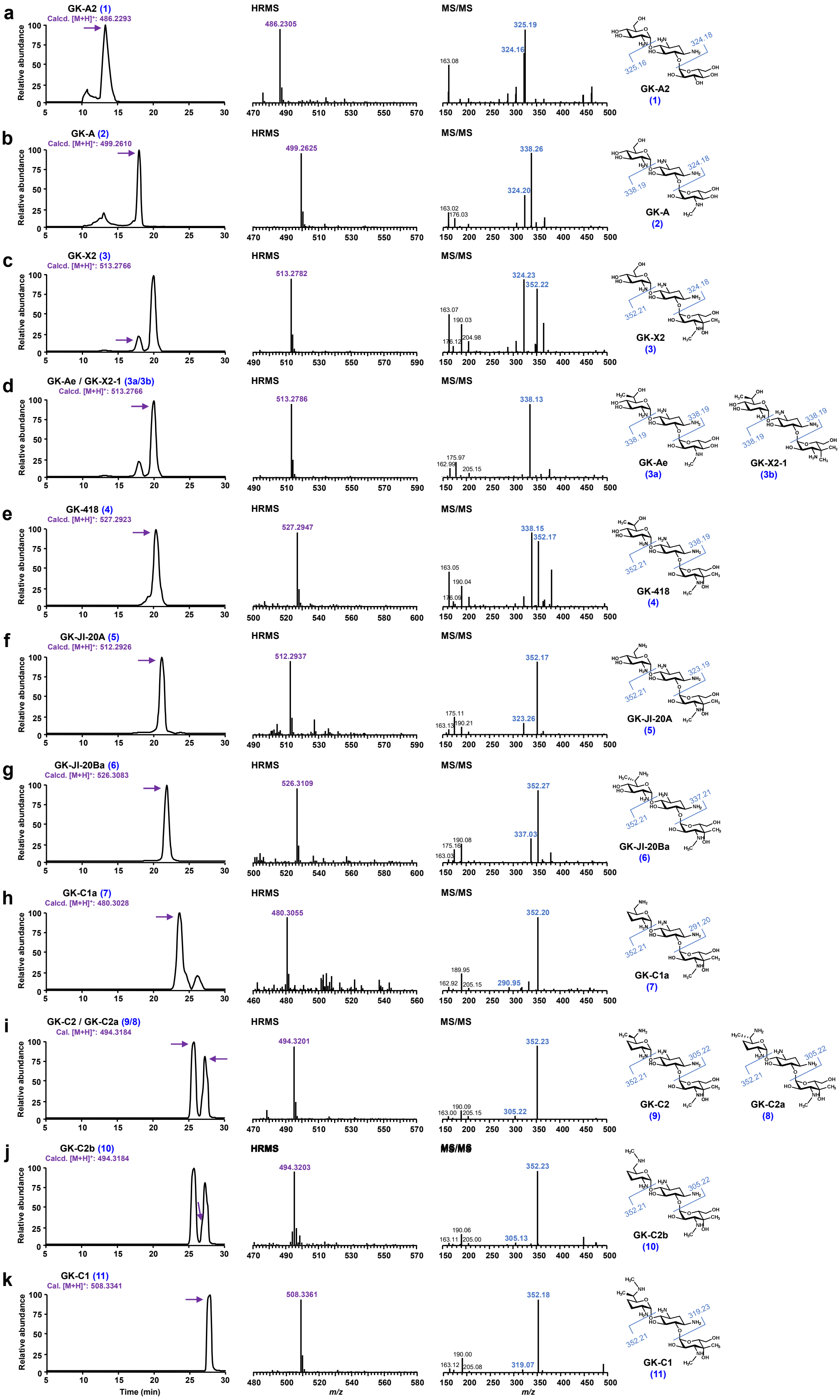

**Supplementary Fig. 4** LC-ESI-HRMS and MS/MS analysis of GK C-complex components and intermediates in ΔgenM2::kanE.

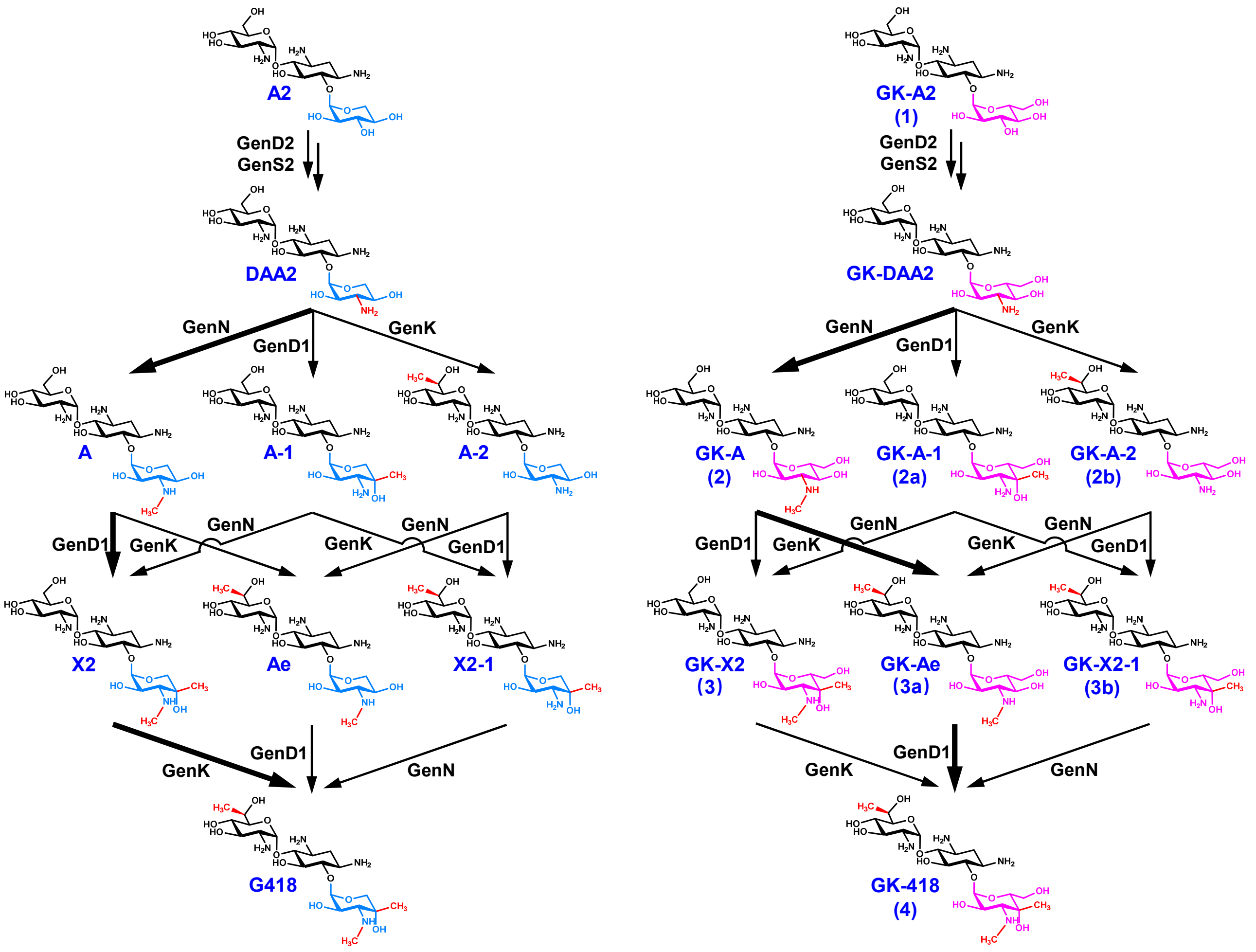

**Supplementary Fig. 5** Proposed methylation pathway in gentamicin and GK biosynthesis. The main pathways are showed in bold arrows.

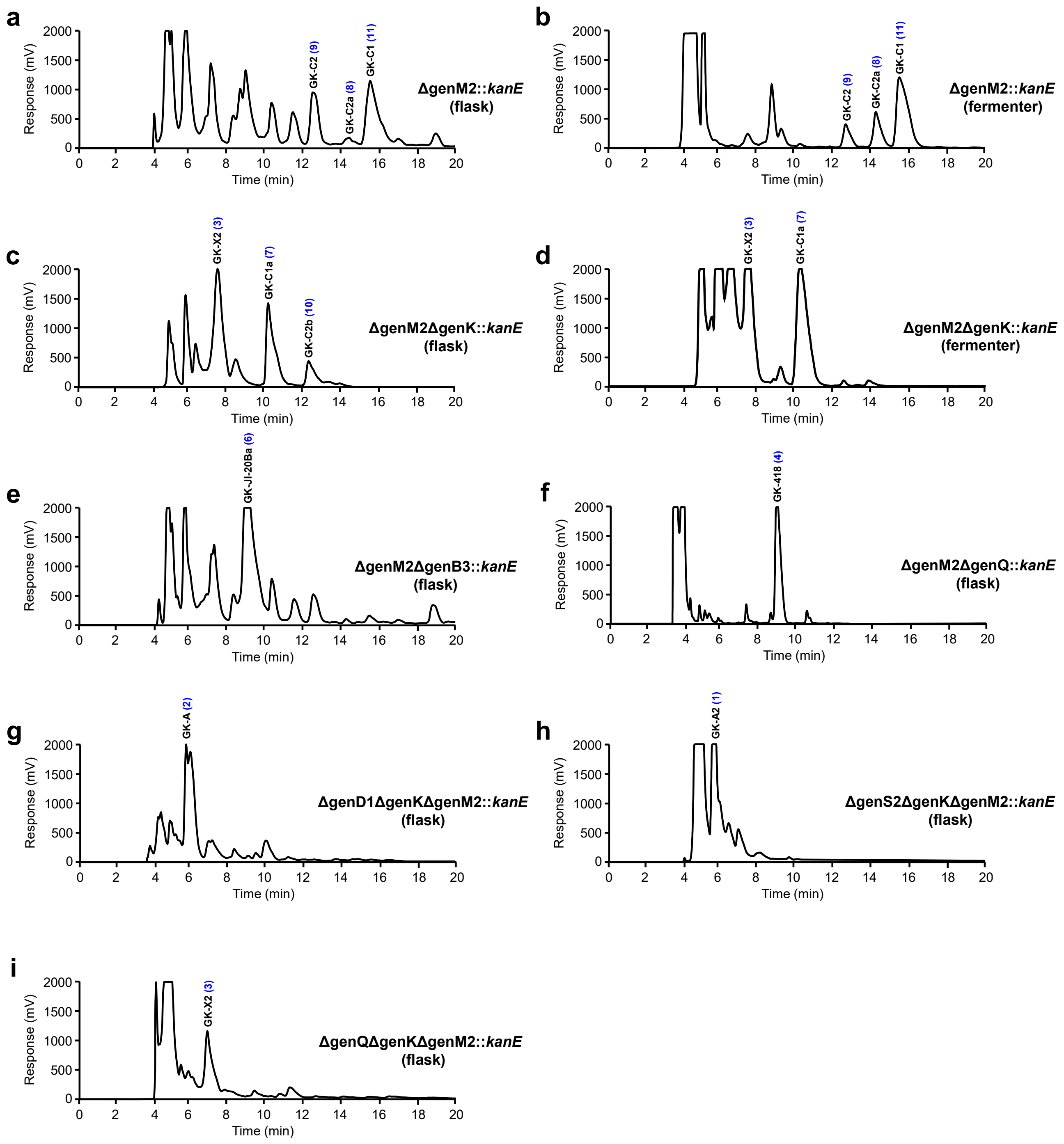

**Supplementary Fig. 6** HPLC-ELSD chromatogram of culture extracts of GK producing and accumulating mutants.

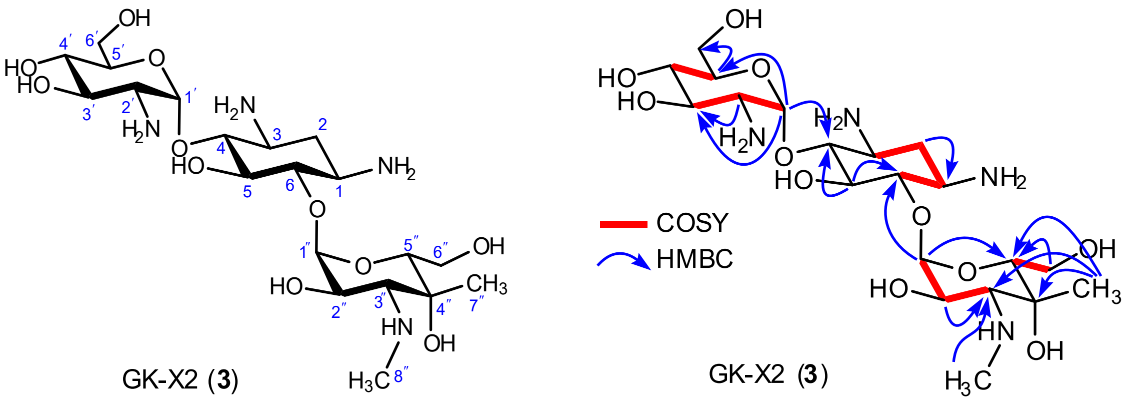

**Supplementary Fig. 7** Chemical structure (left), key [1]H–[1]H COSY and HMBC correlations (right) of GK-X2 (**3**).

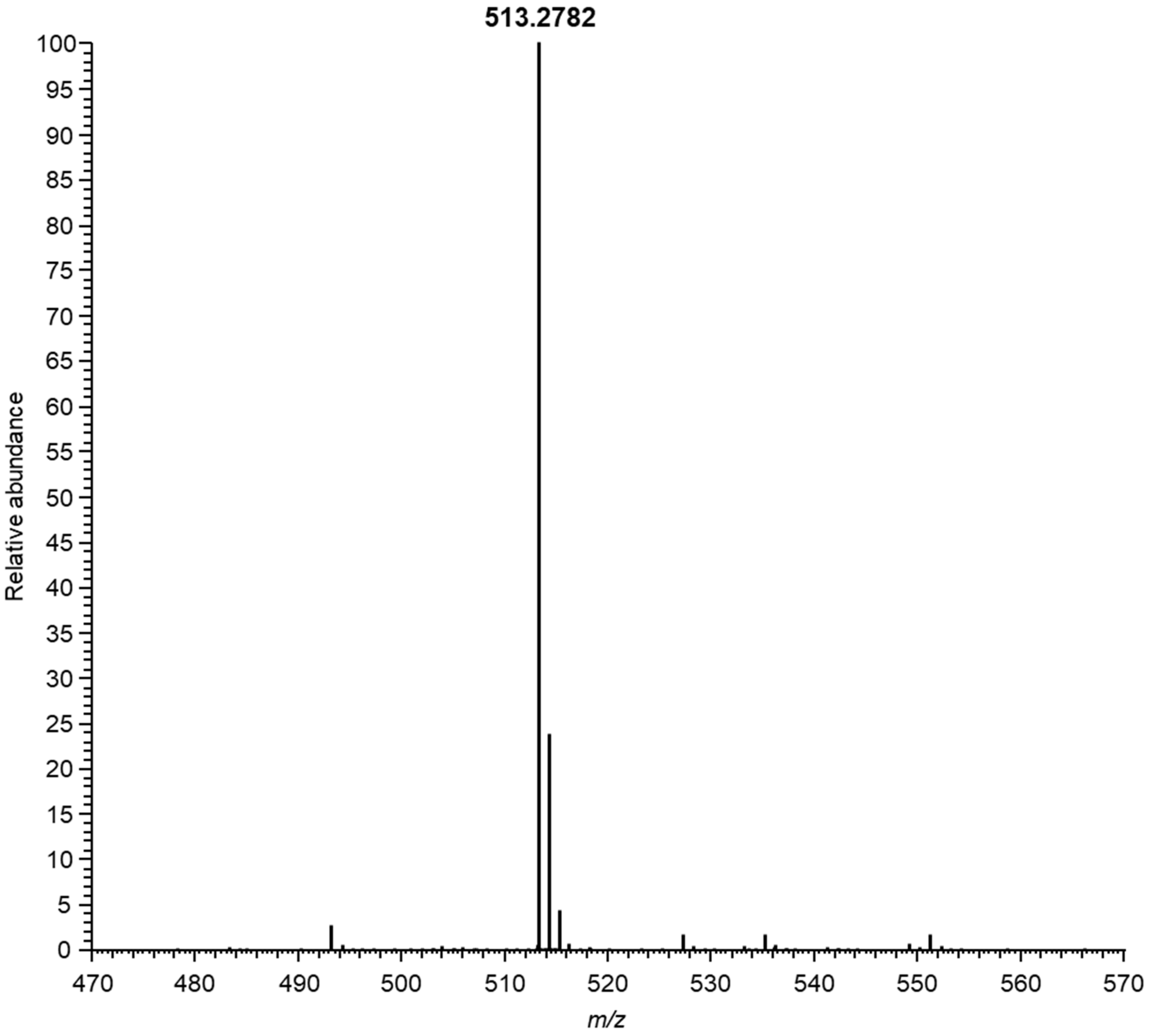

**Supplementary Fig. 8** ESI-HRMS spectrum of GK-X2 (**3**).

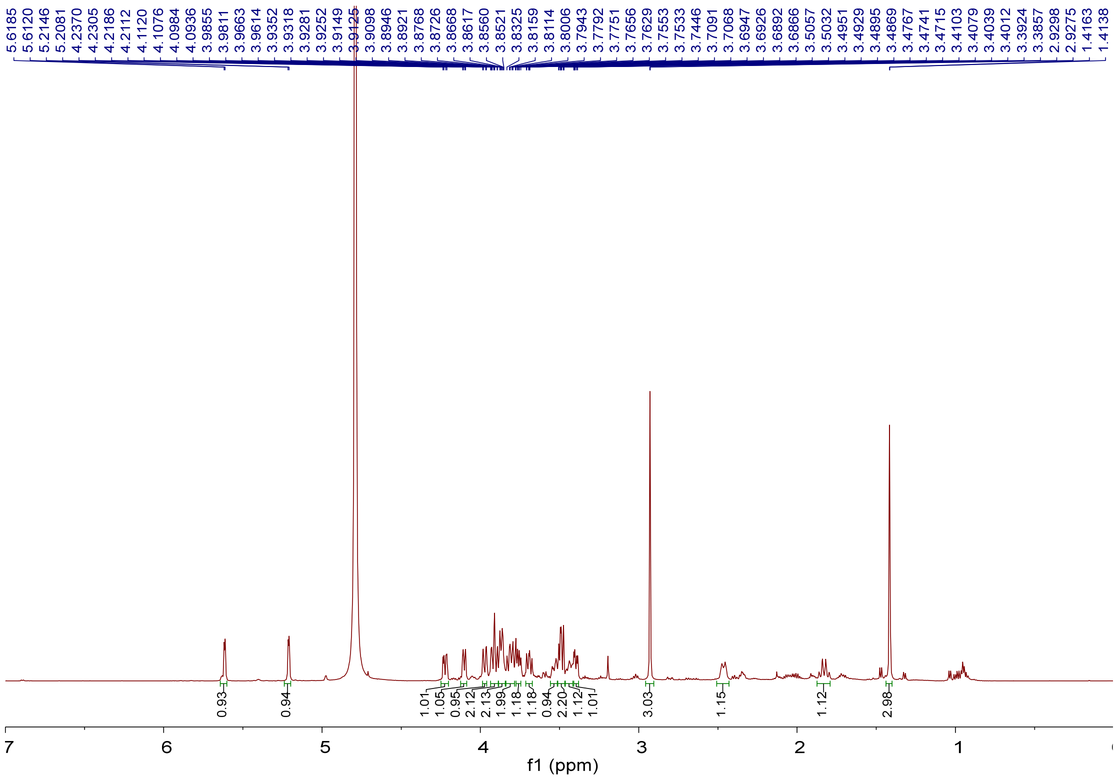

**Supplementary Fig. 9** $^{1}H$ NMR spectrum (600 MHz, $D_2O$) of GK-X2 (**3**).

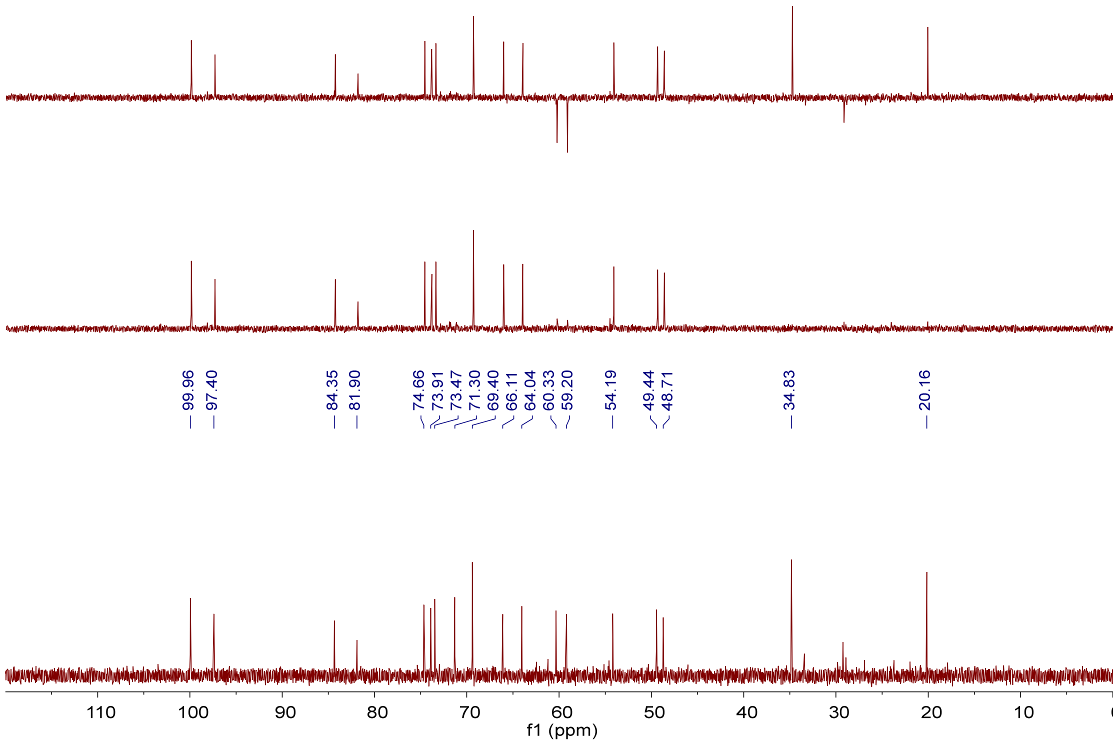

**Supplementary Fig. 10** $^{13}C$ NMR, DEPT-90 and DEPT-135 spectra (150 MHz, $D_2O$) of GK-X2 (**3**).

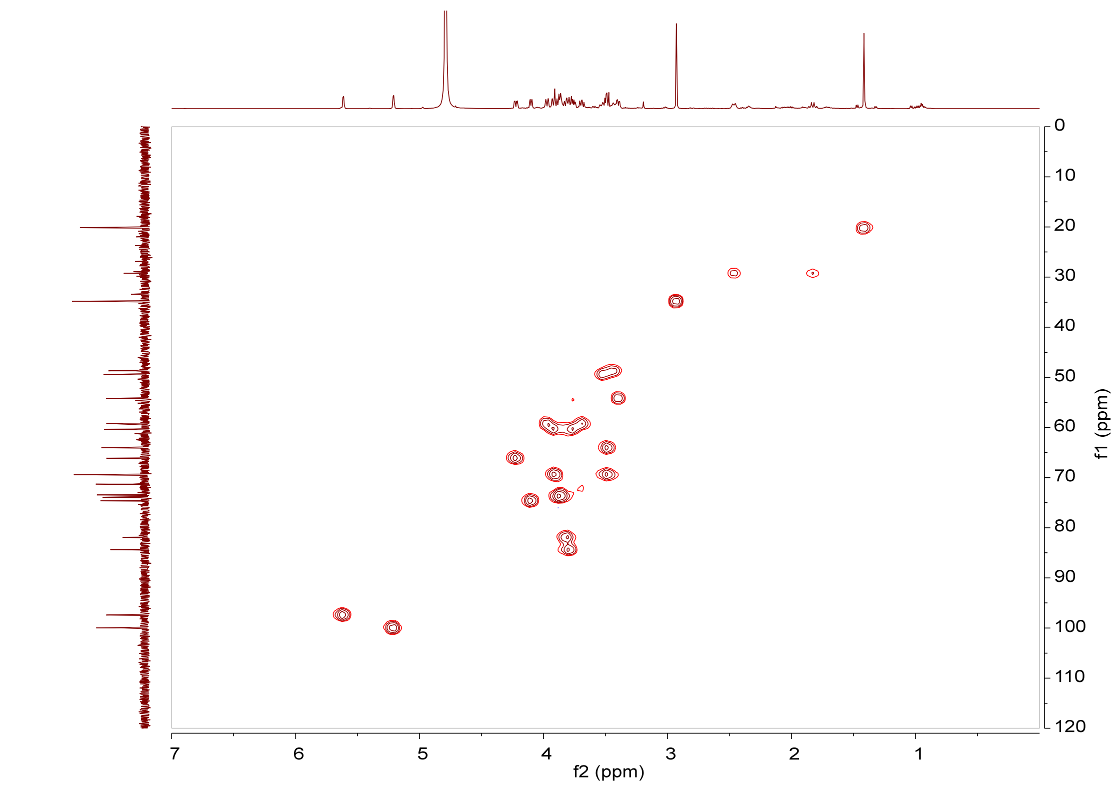

**Supplementary Fig. 11** HSQC spectrum (600 MHz, $D_2O$) of GK-X2 (**3**).

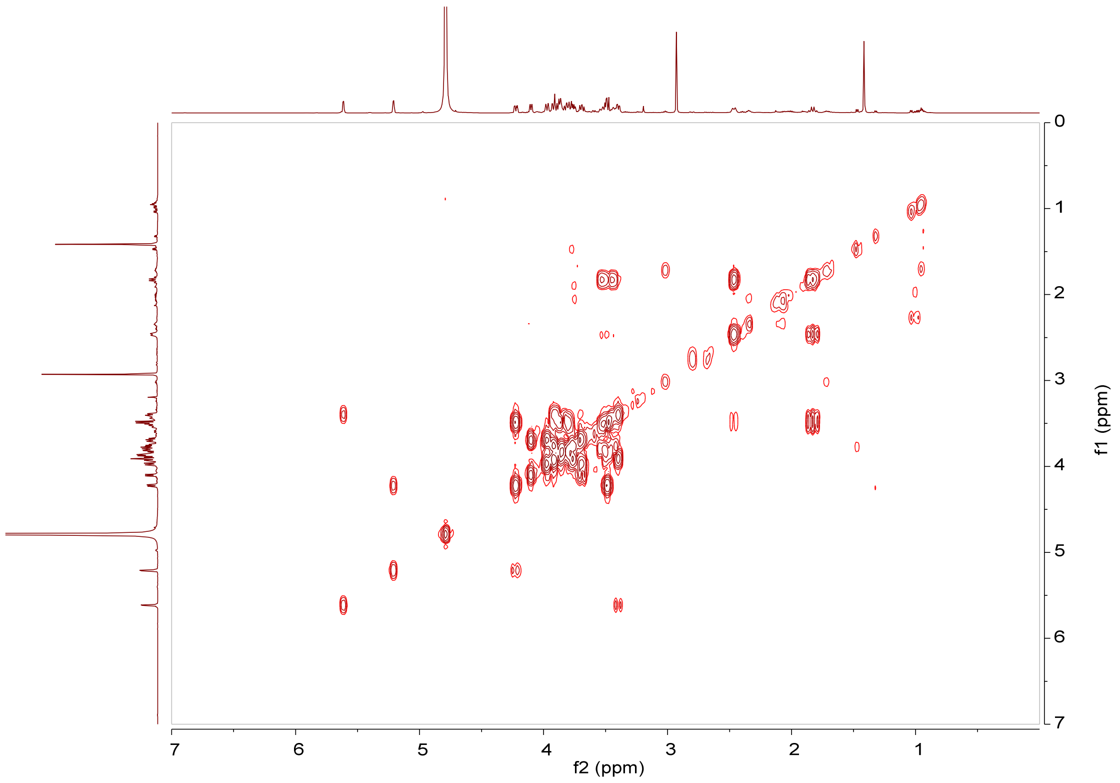

**Supplementary Fig. 12** $^1H$–$^1H$ COSY spectrum (600 MHz, $D_2O$) of GK-X2 (**3**).

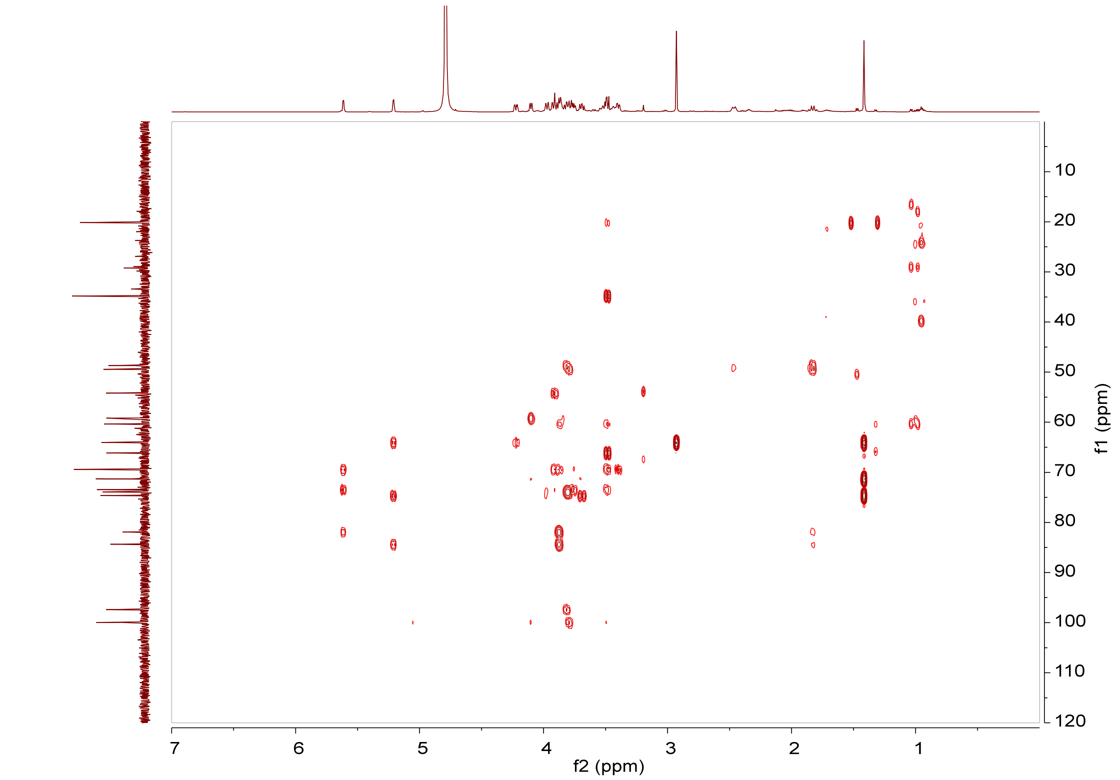

**Supplementary Fig. 13** HMBC spectrum (600 MHz, $D_2O$) of GK-X2 (**3**).

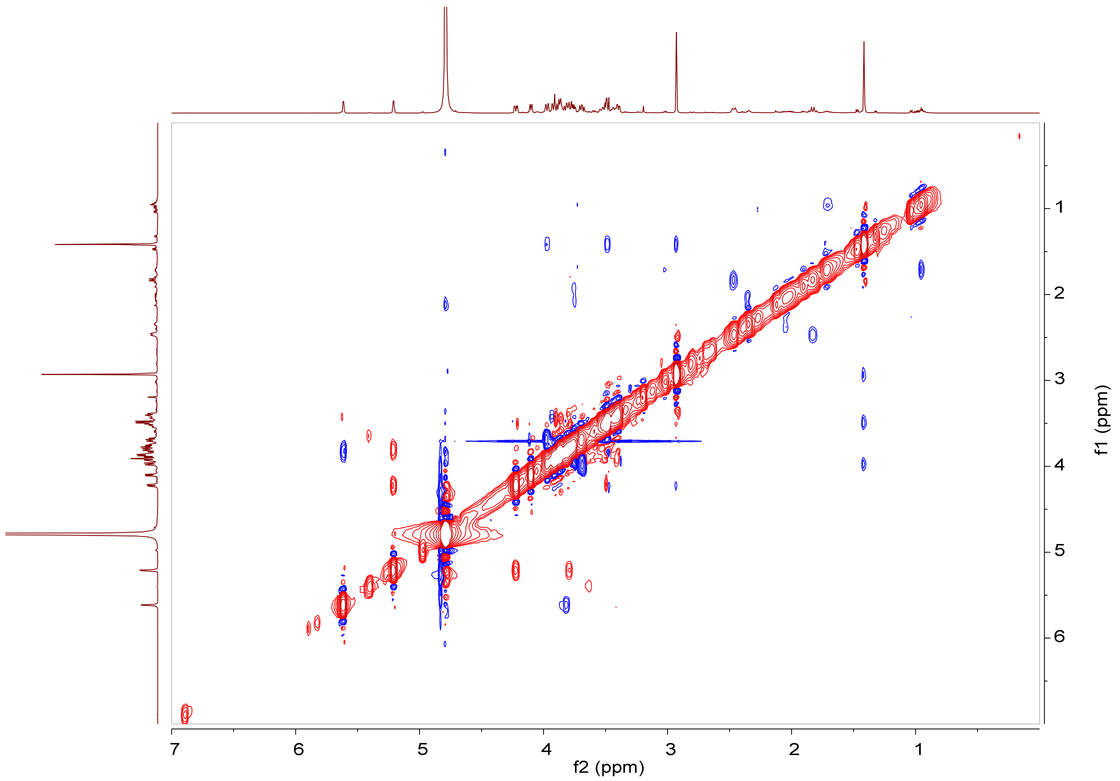

**Supplementary Fig. 14** NOESY spectrum (600 MHz, $D_2O$) of GK-X2 (**3**).

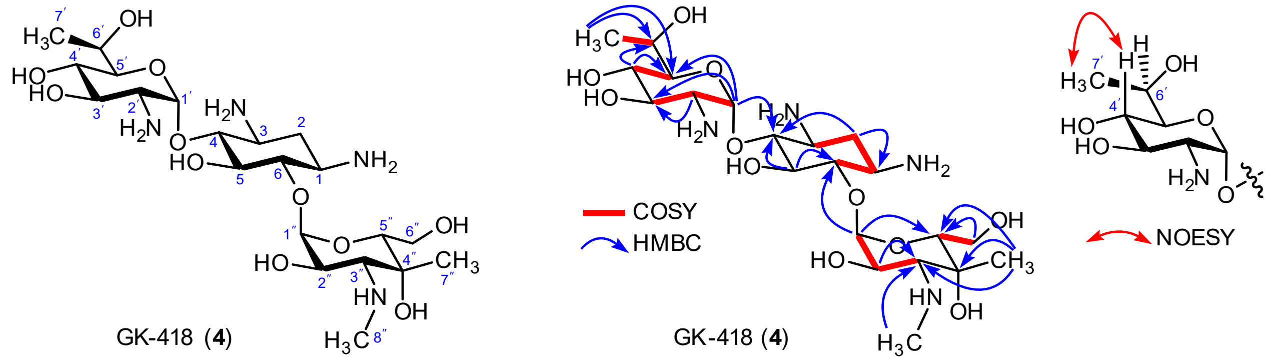

**Supplementary Fig. 15** Chemical structure (left), key $^{1}$H–$^{1}$H COSY, HMBC and NOESY correlations of GK-418 (**4**) (right).

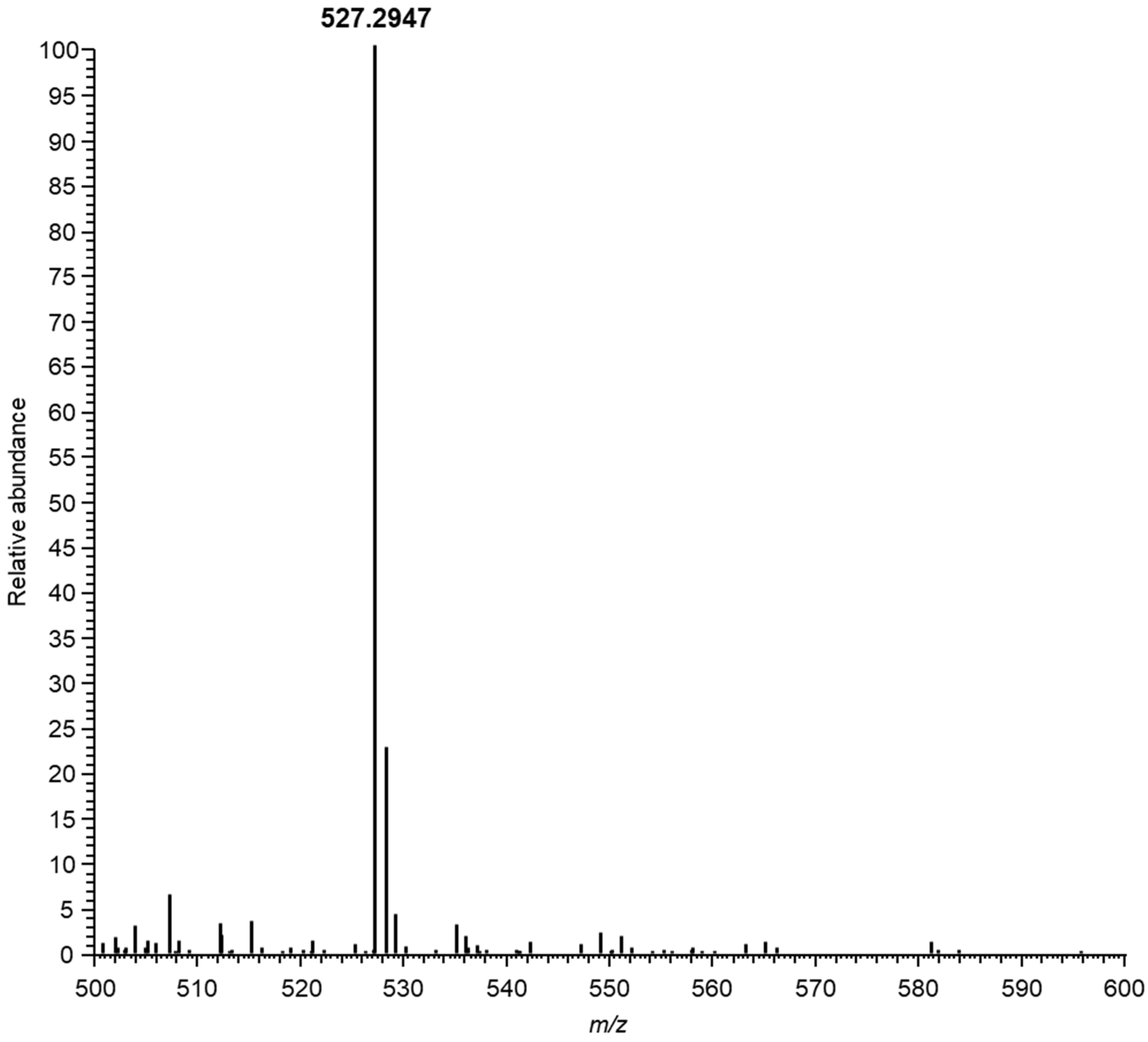

**Supplementary Fig. 16** ESI-HRMS spectrum of GK-418 (**4**).

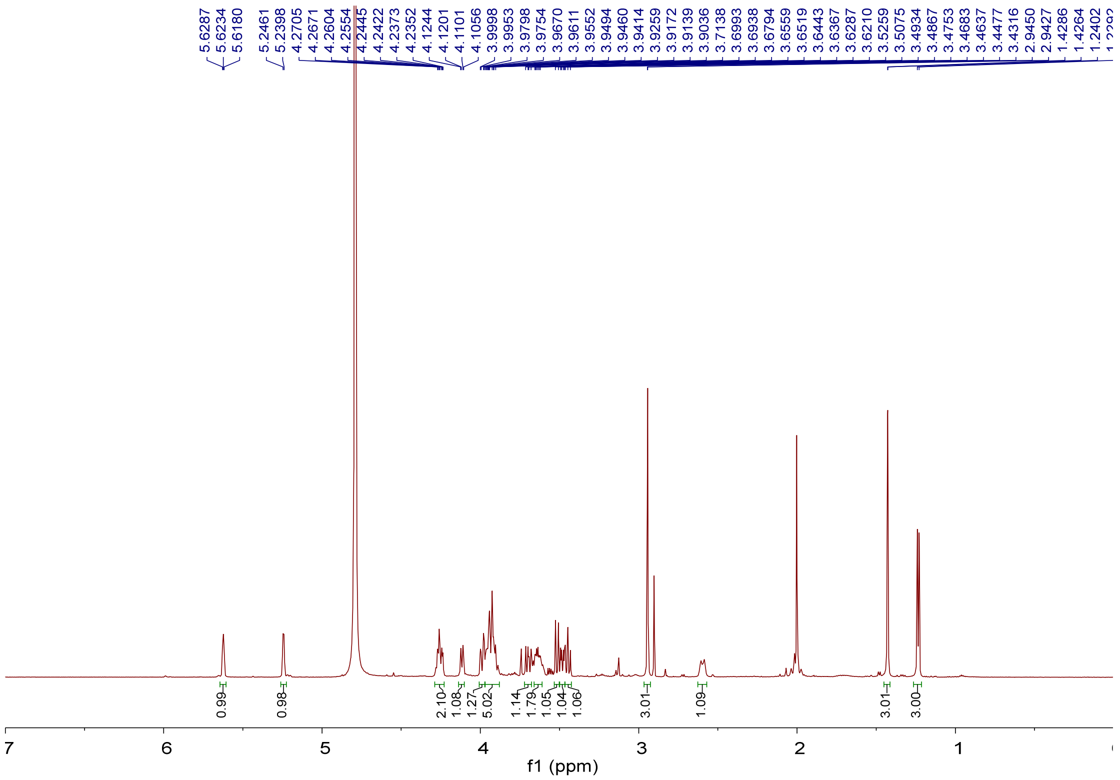

**Supplementary Fig. 17** ¹H NMR spectrum (600 MHz, $D_2O$) of GK-418 (**4**).

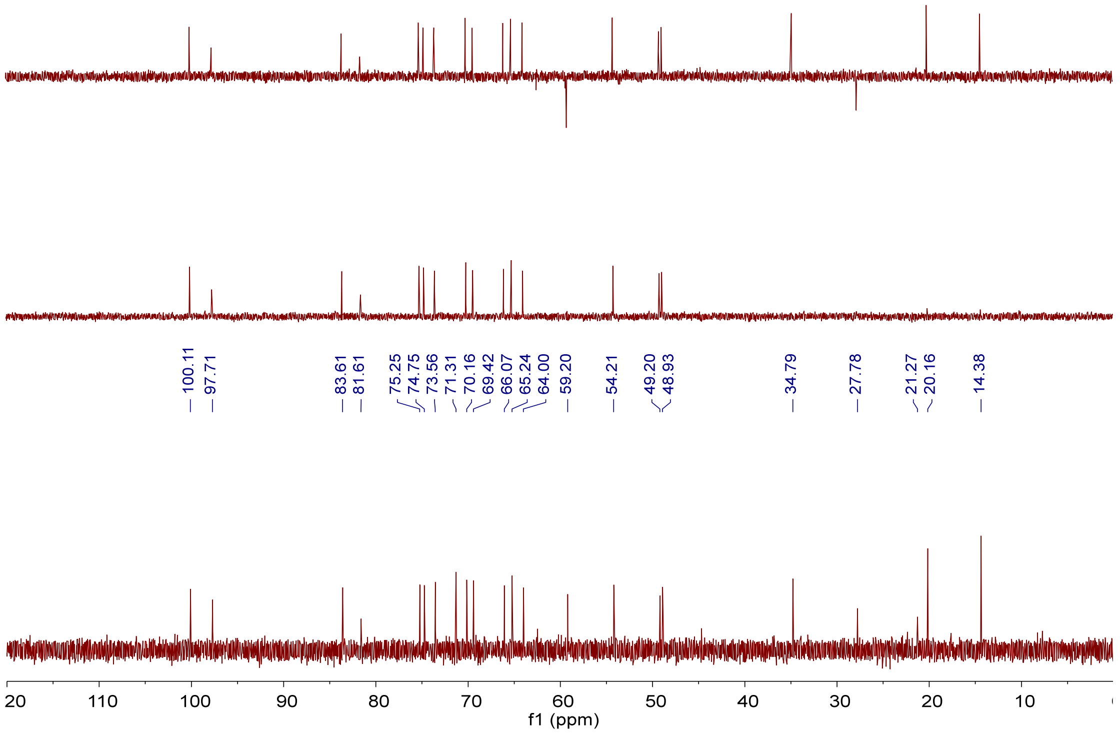

**Supplementary Fig. 18** ¹³C NMR, DEPT-90 and DEPT-135 spectra (150 MHz, $D_2O$) of GK-418 (**4**).

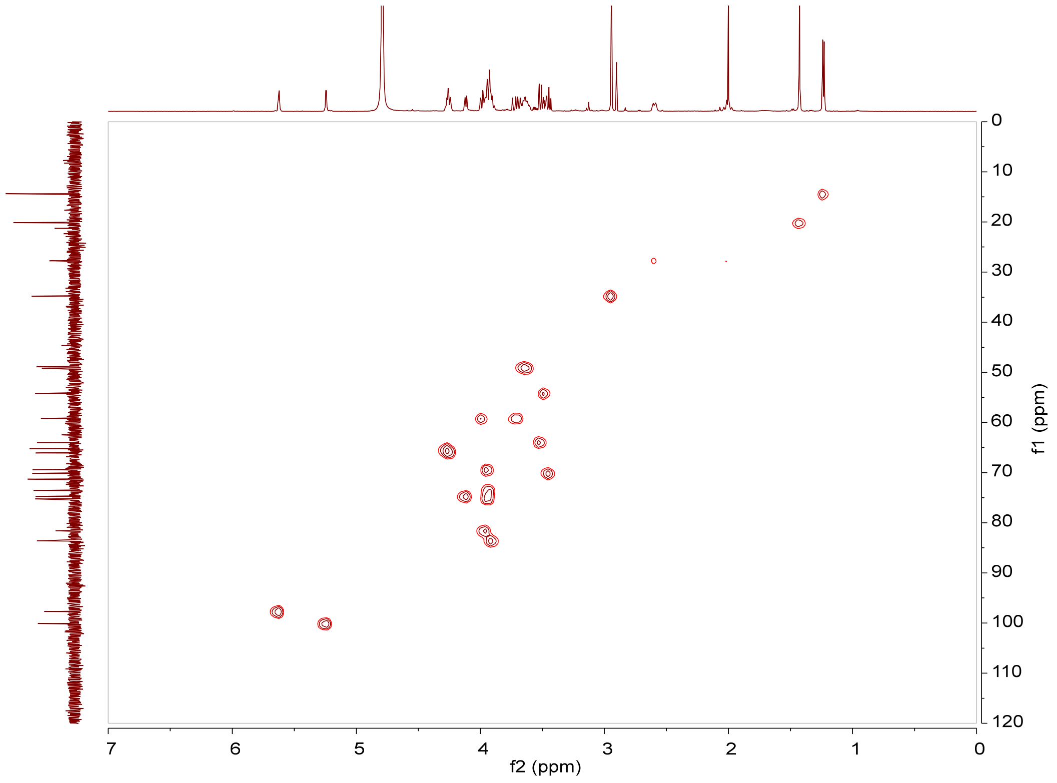

**Supplementary Fig. 19** HSQC spectrum (600 MHz, $D_2O$) of GK-418 (**4**).

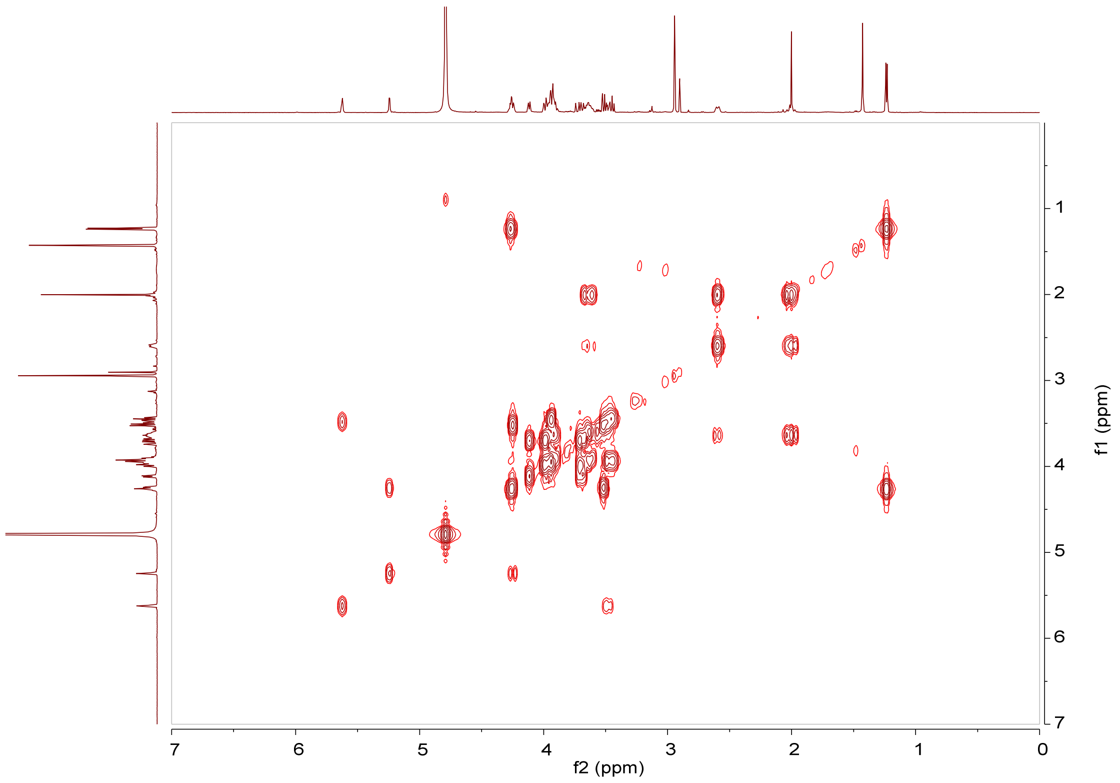

**Supplementary Fig. 20** $^1H$–$^1H$ COSY spectrum (600 MHz, $D_2O$) of GK-418 (**4**).

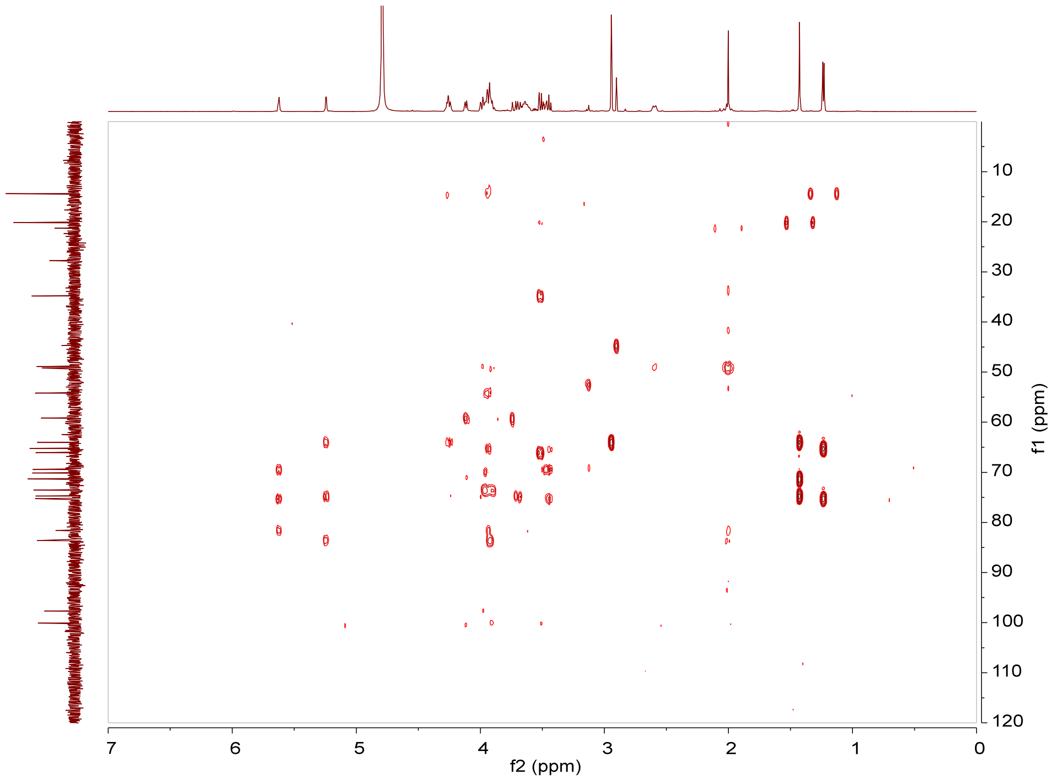

**Supplementary Fig. 21** HMBC spectrum (600 MHz, $D_2O$) of GK-418 (**4**).

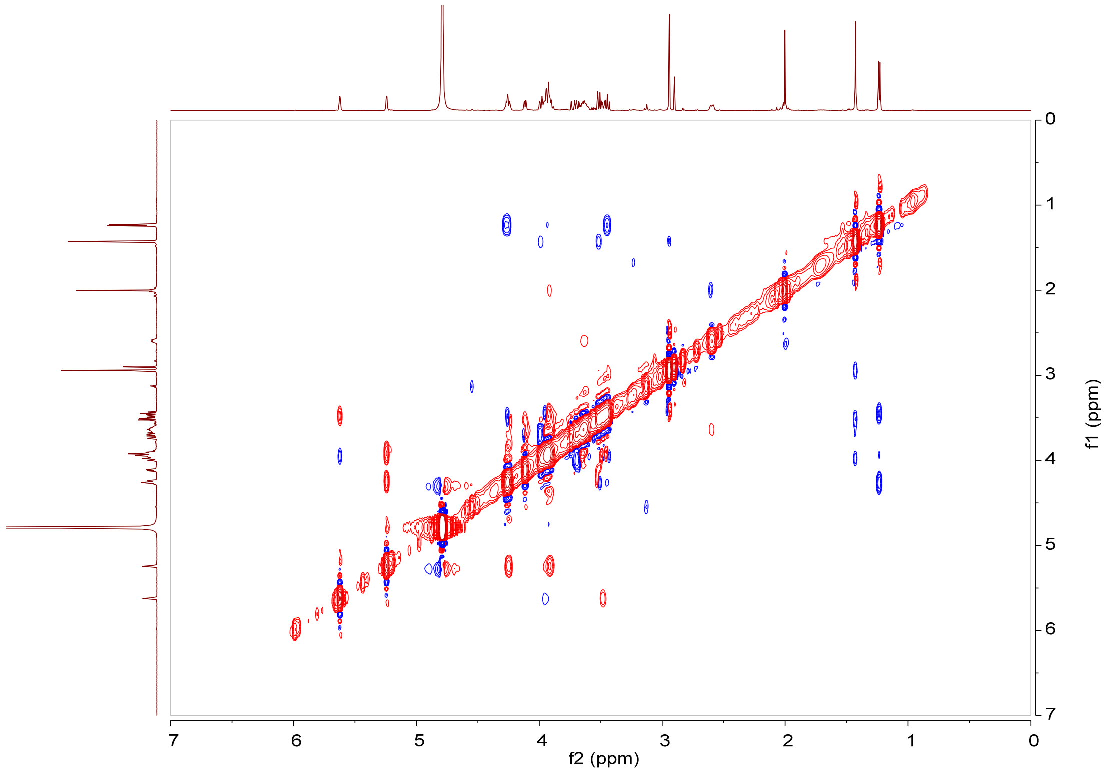

**Supplementary Fig. 22** NOESY spectrum (600 MHz, $D_2O$) of GK-418 (**4**).

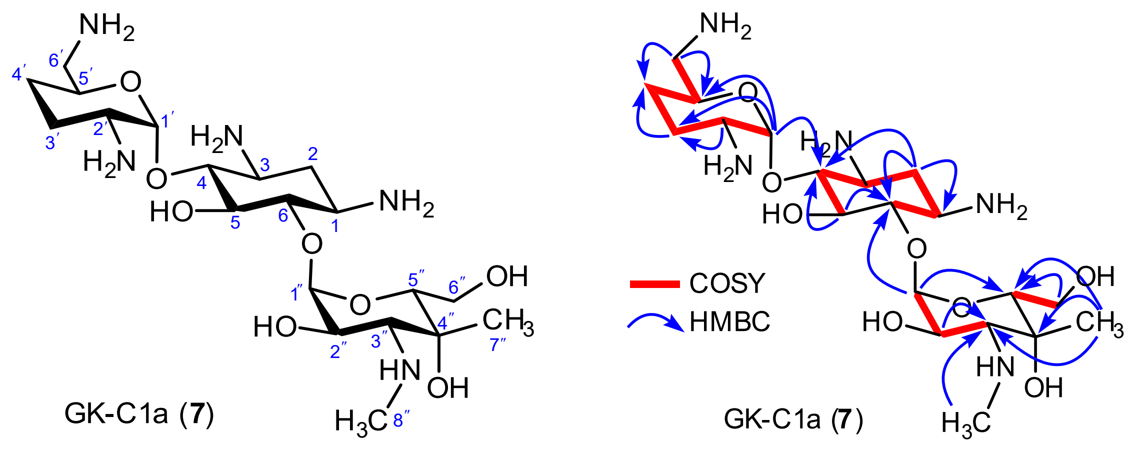

**Supplementary Fig. 23** Chemical structure (left), key $^{1}H$–$^{1}H$ COSY and HMBC correlations of GK-C1a (**7**) (right).

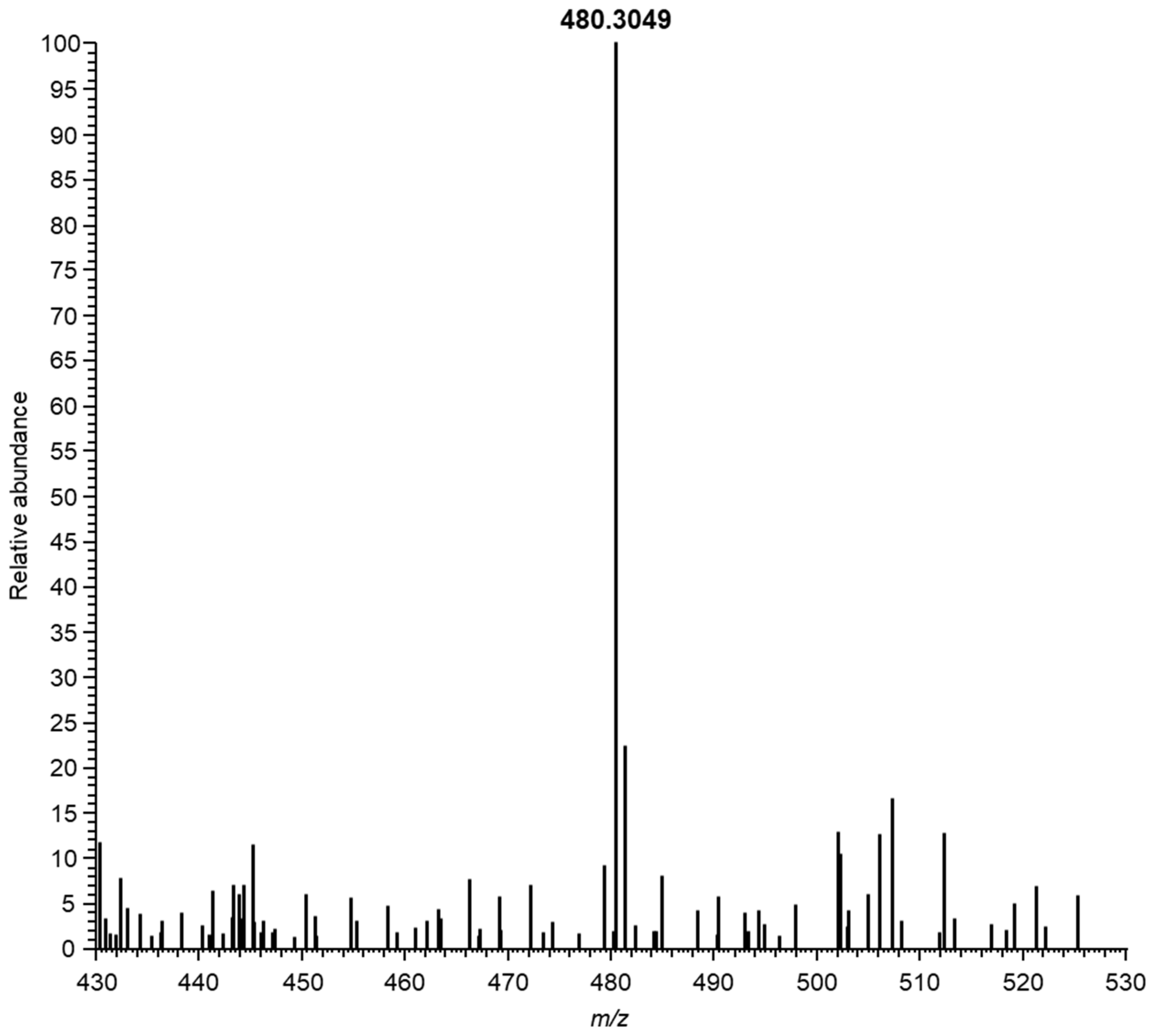

**Supplementary Fig. 24** ESI-HRMS spectrum of GK-C1a (**7**).

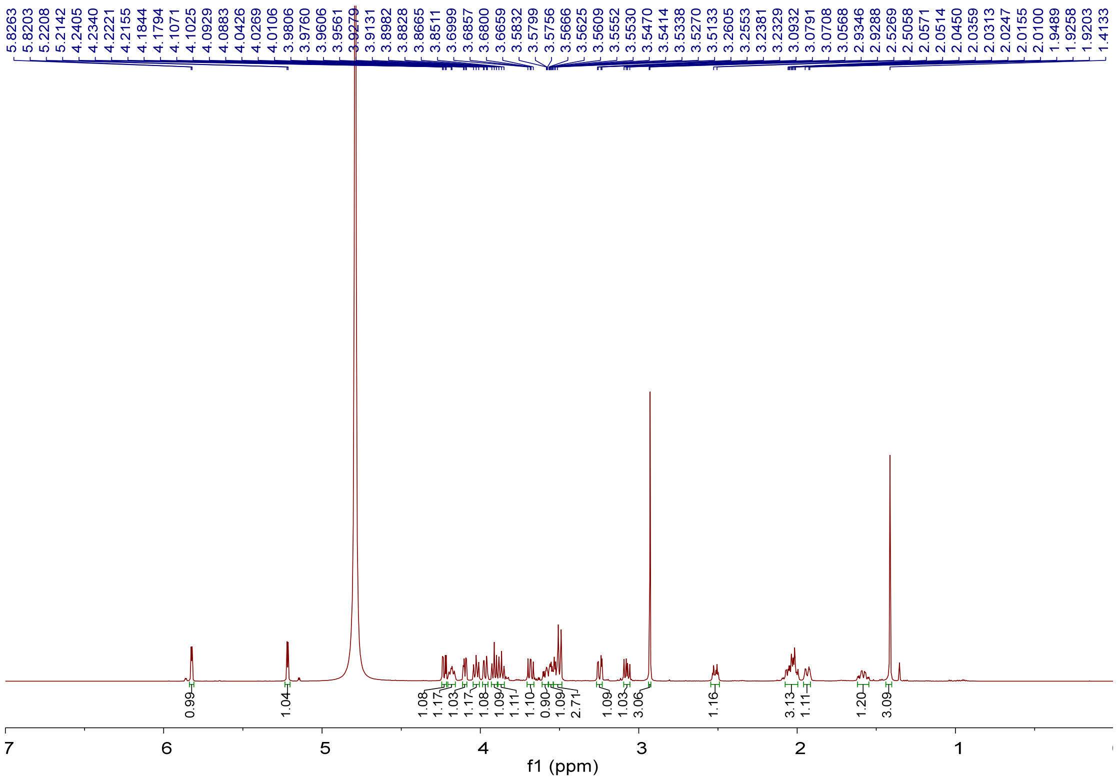

**Supplementary Fig. 25** $^1$H NMR spectrum (600 MHz, $D_2O$) of GK-C1a (**7**).

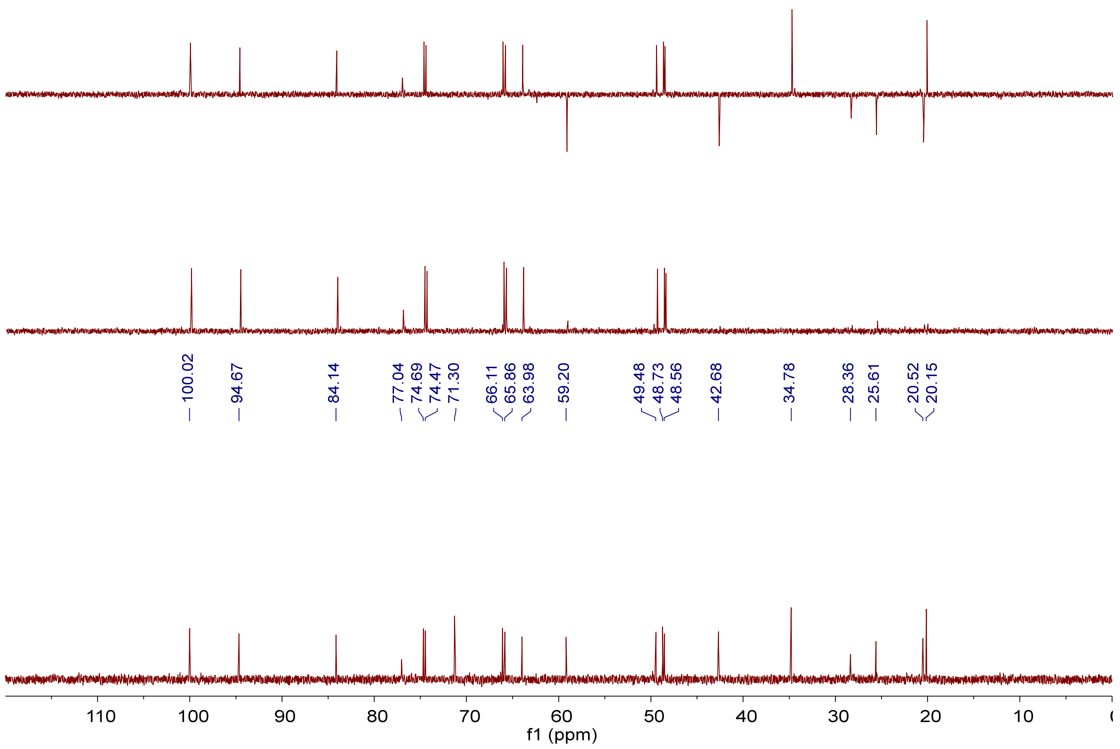

**Supplementary Fig. 26** $^{13}$C NMR, DEPT-90 and DEPT-135 spectra (150 MHz, $D_2O$) of GK-C1a (**7**).

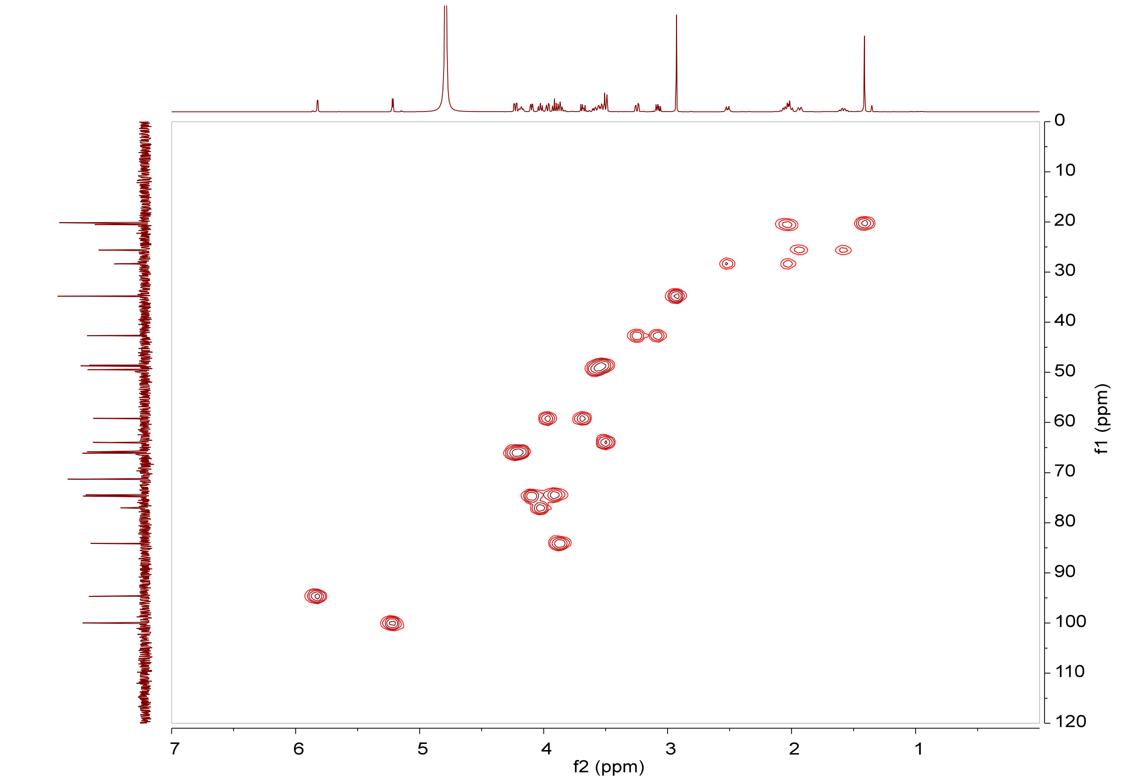

**Supplementary Fig. 27** HSQC spectrum (600 MHz, $D_2O$) of GK-C1a (**7**).

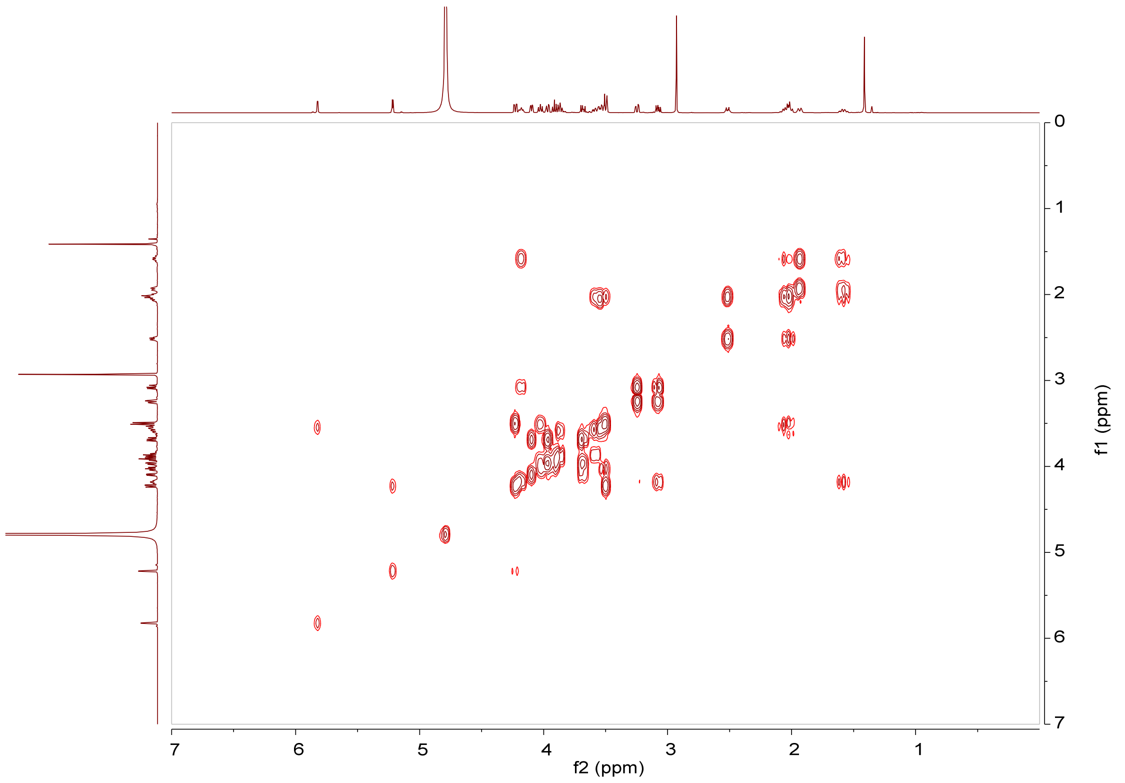

**Supplementary Fig. 28** $^1H$–$^1H$ COSY spectrum (600 MHz, $D_2O$) of GK-C1a (**7**).

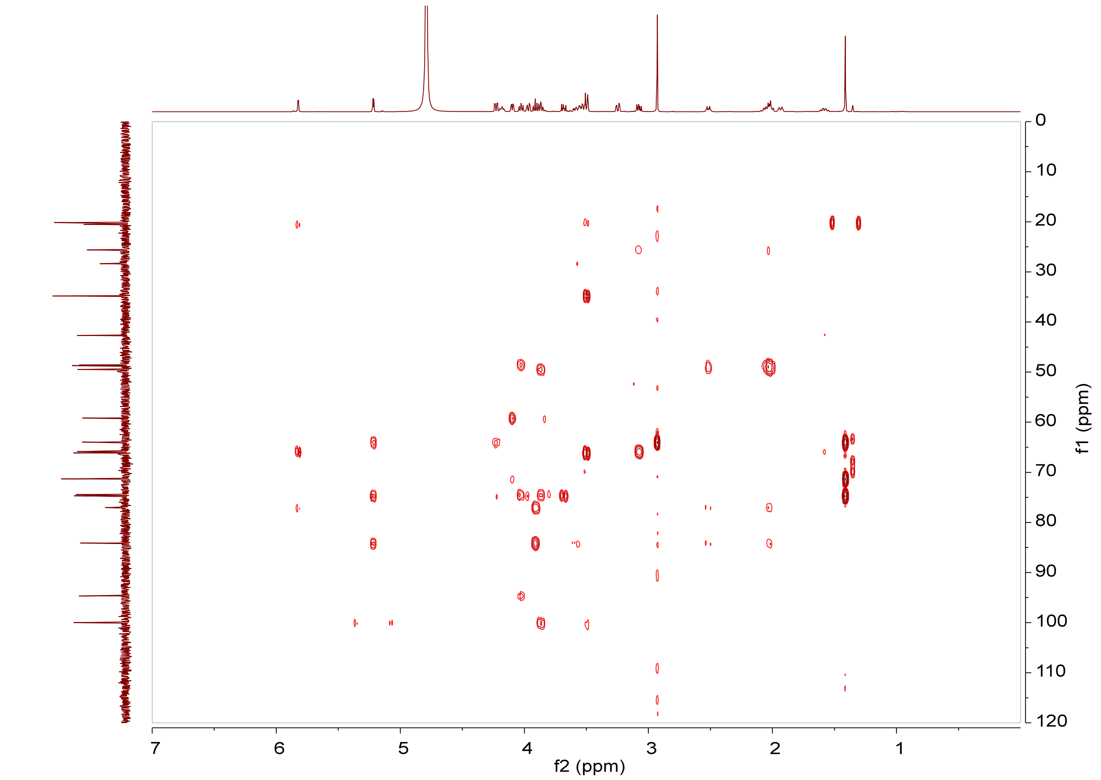

**Supplementary Fig. 29** HMBC spectrum (600 MHz, $D_2O$) of GK-C1a (**7**).

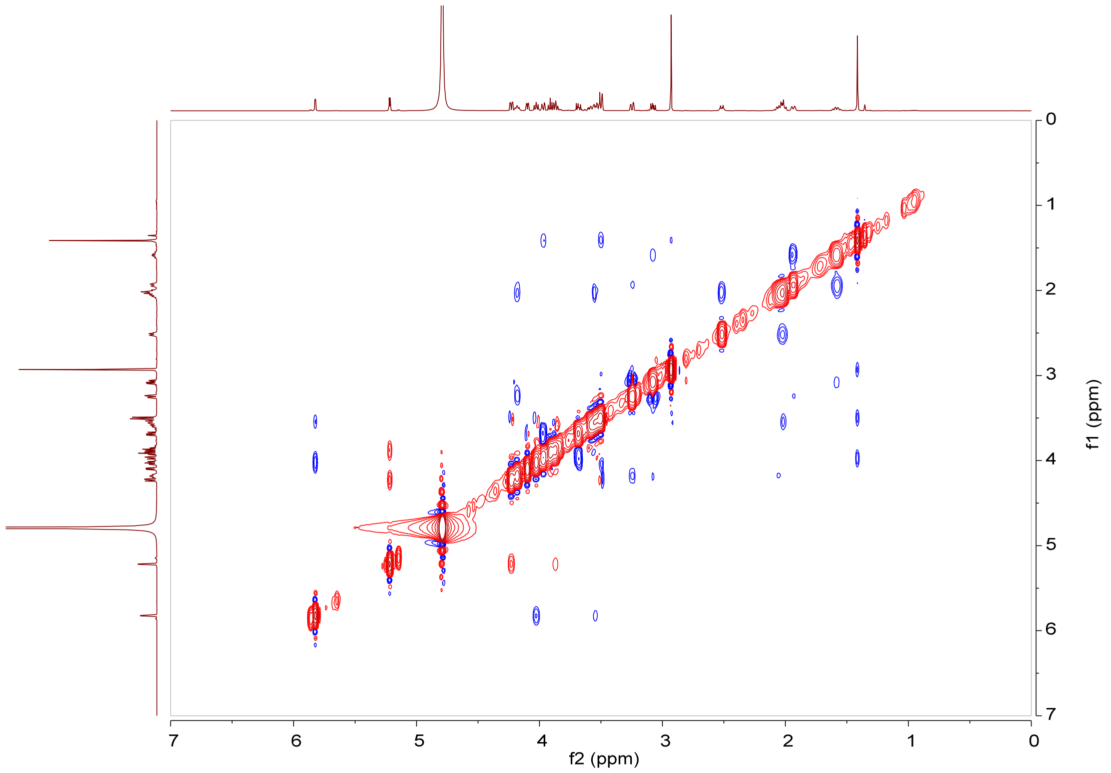

**Supplementary Fig. 30** NOESY spectrum (600 MHz, $D_2O$) of GK-C1a (**7**).

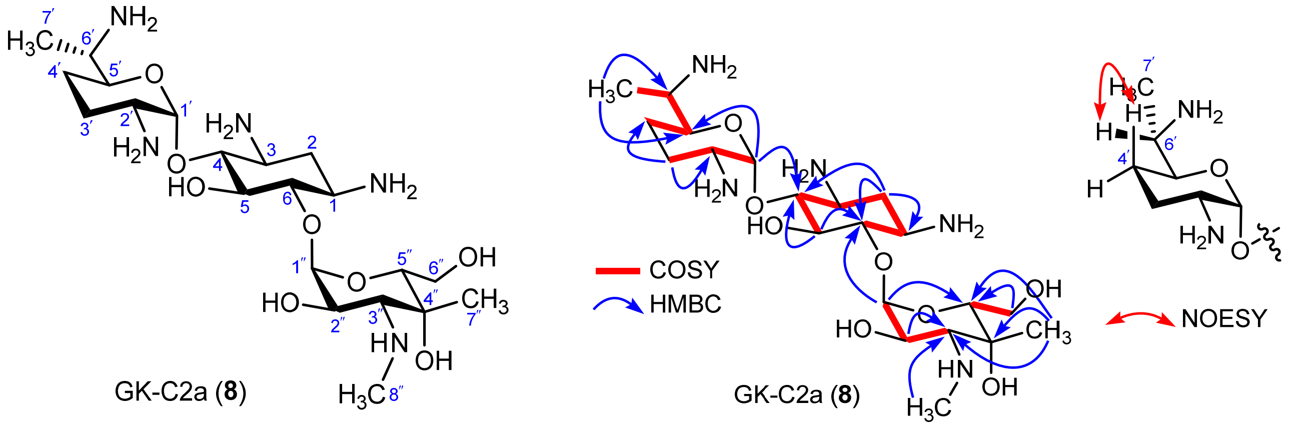

**Supplementary Fig. 31** Chemical structure (left), key $^{1}H$–$^{1}H$ COSY, HMBC and NOESY correlations of GK-C2a (**8**) (right).

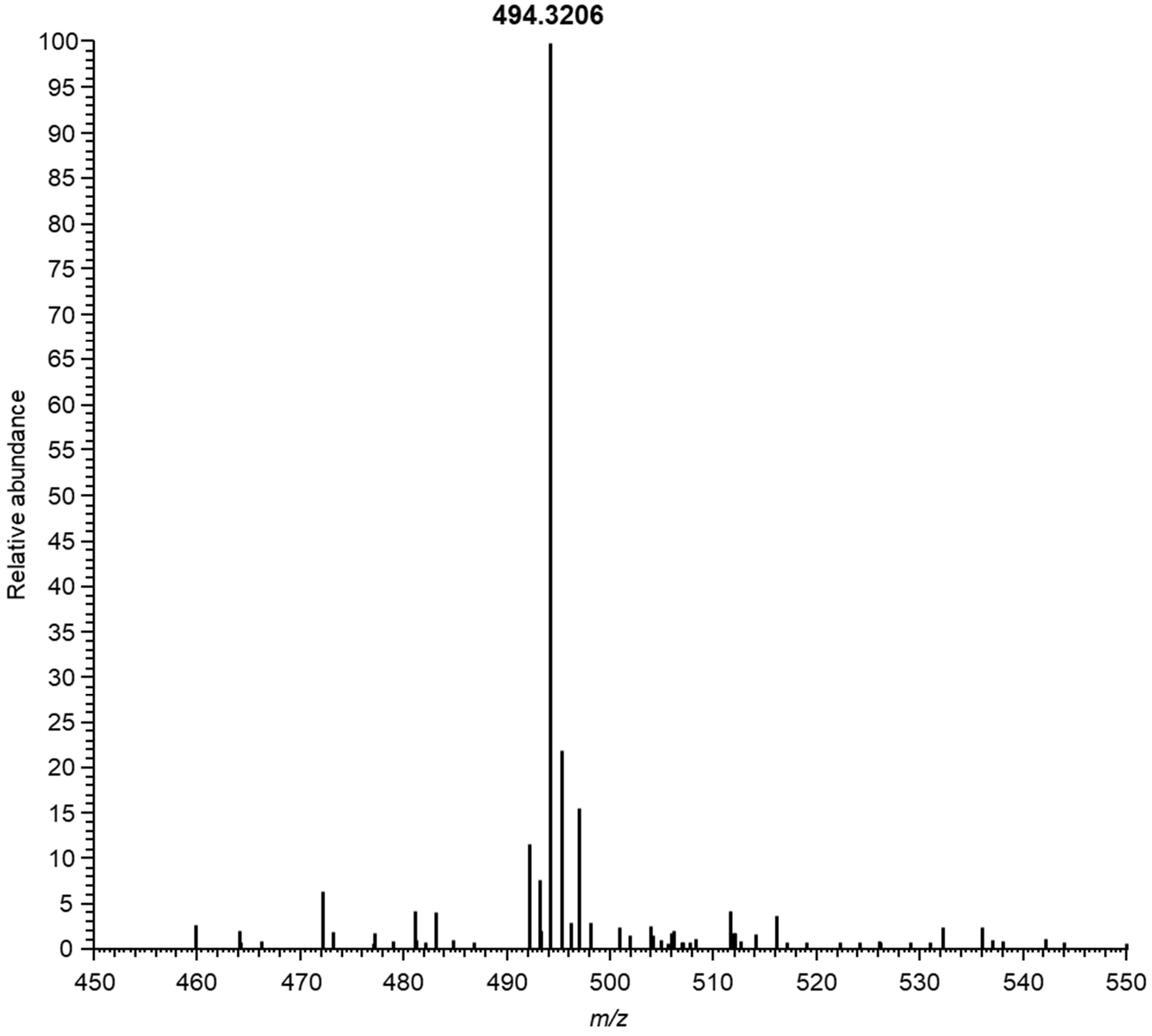

**Supplementary Fig. 32** ESI-HRMS spectrum of GK-C2a (**8**).

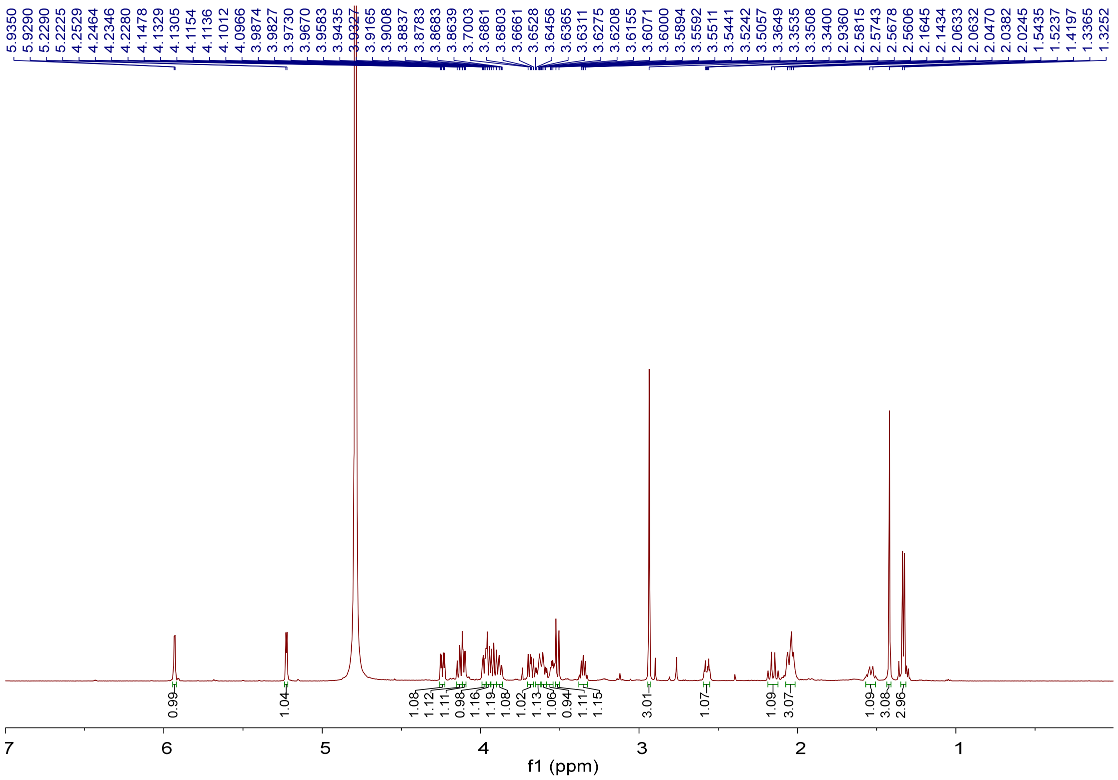

**Supplementary Fig. 33** $^{1}H$ NMR spectrum (600 MHz, $D_2O$) of GK-C2a (**8**).

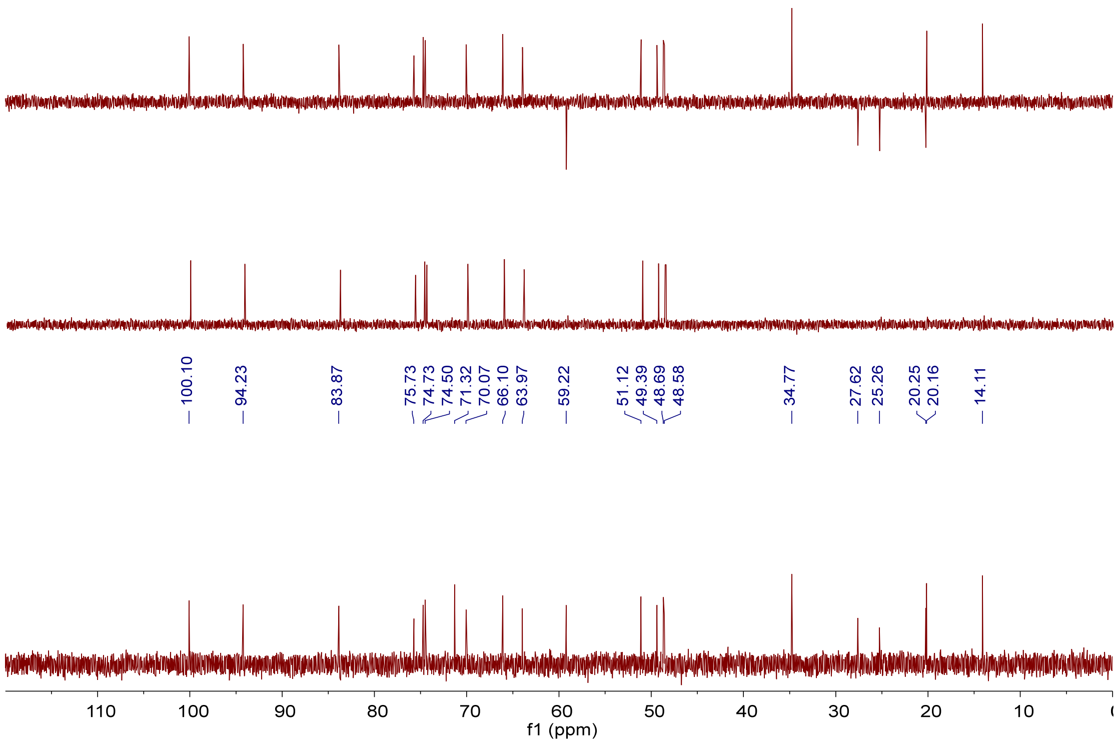

**Supplementary Fig. 34** $^{13}C$ NMR, DEPT-90 and DEPT-135 spectra (150 MHz, $D_2O$) of GK-C2a (**8**).

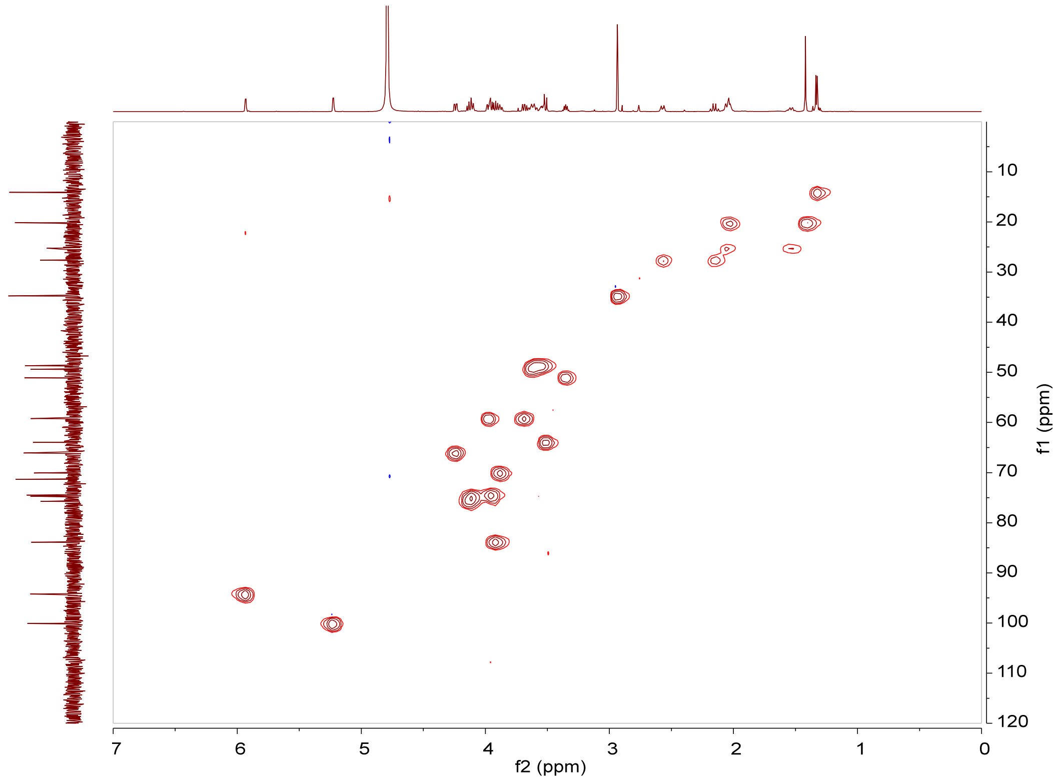

**Supplementary Fig. 35** HSQC spectrum (600 MHz, $D_2O$) of GK-C2a (**8**).

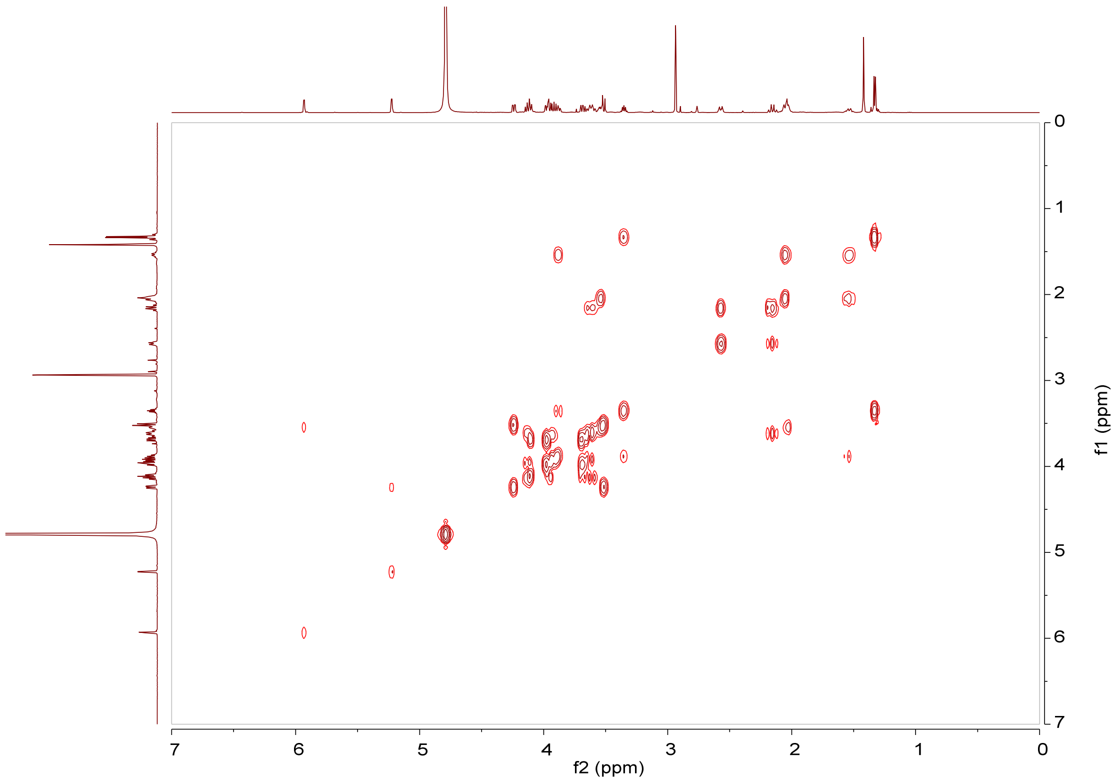

**Supplementary Fig. 36** $^1H$–$^1H$ COSY spectrum (600 MHz, $D_2O$) of GK-C2a (**8**).

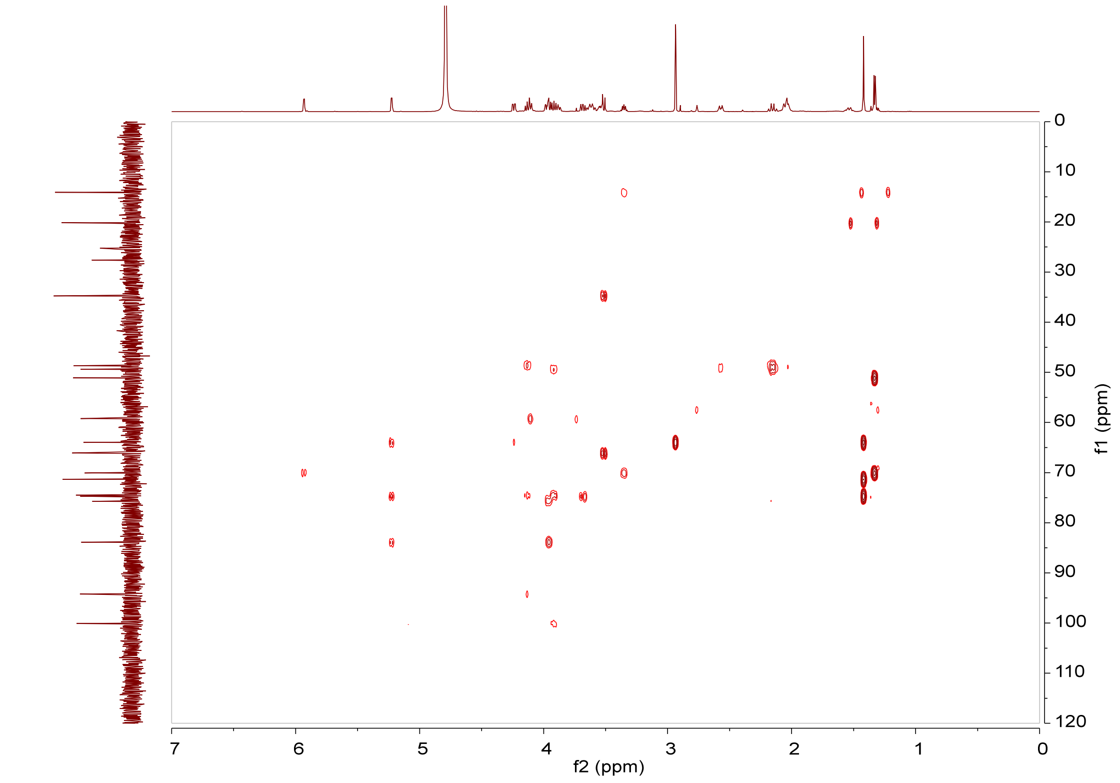

**Supplementary Fig. 37** HMBC spectrum (600 MHz, $D_2O$) of GK-C2a (**8**).

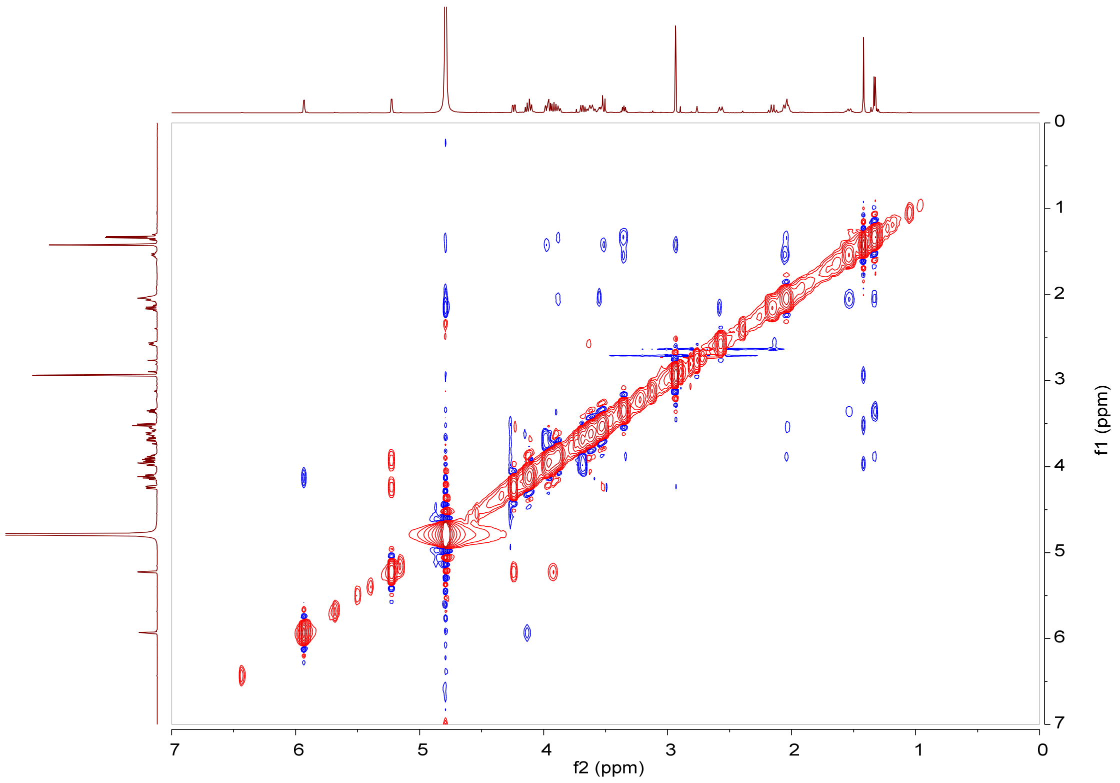

**Supplementary Fig. 38** NOESY spectrum (600 MHz, $D_2O$) of GK-C2a (**8**).

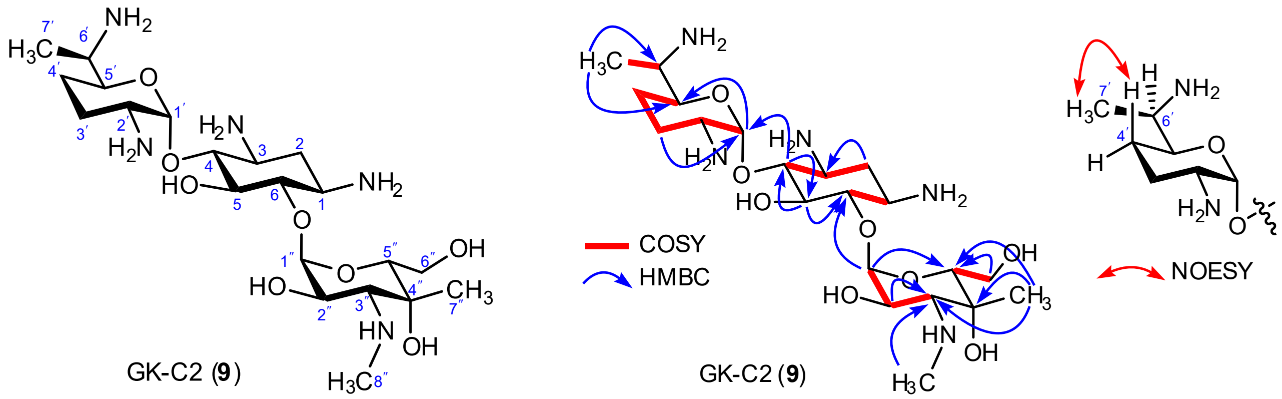

**Supplementary Fig. 39** Chemical structure (left), key $^{1}H$–$^{1}H$ COSY, HMBC and NOESY correlations of GK-C2 (**9**) (right).

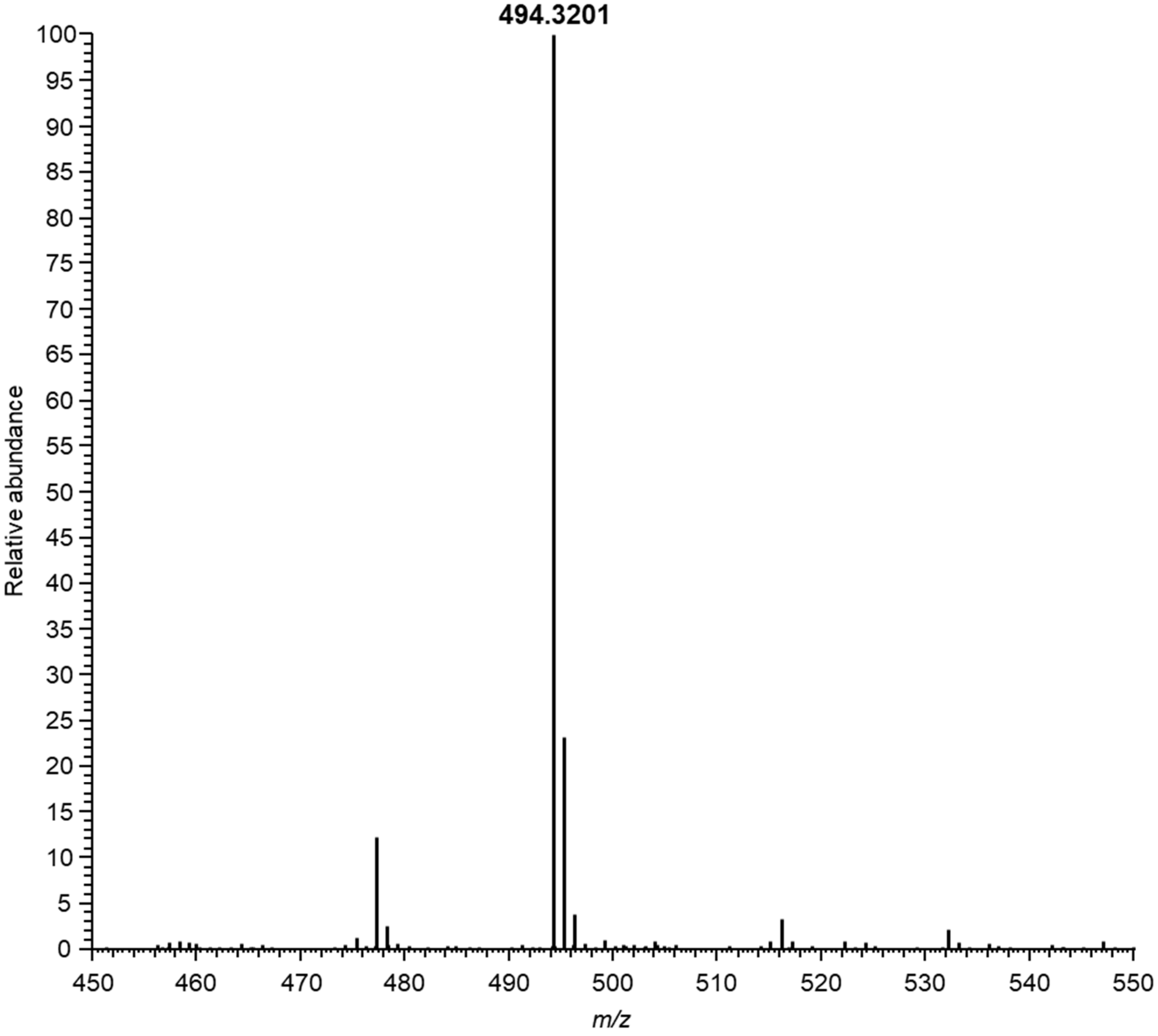

**Supplementary Fig. 40** ESI-HRMS spectrum of GK-C2 (**9**).

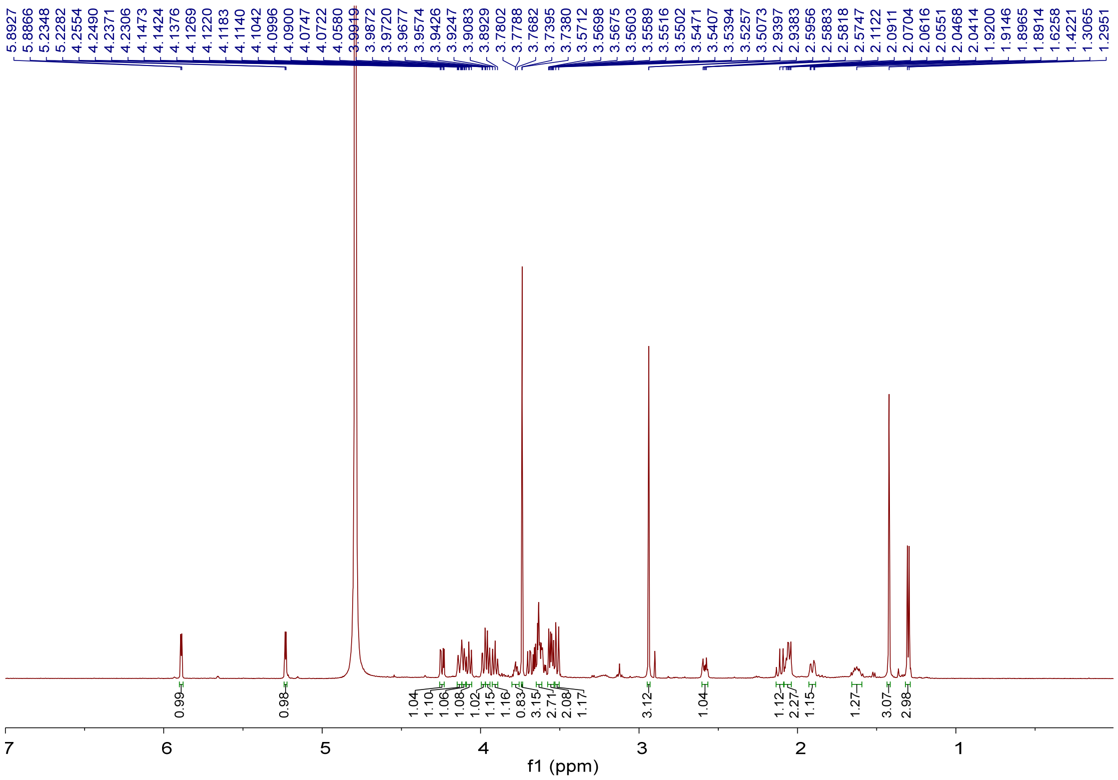

**Supplementary Fig. 41** $^{1}H$ NMR spectrum (600 MHz, $D_2O$) of GK-C2 (**9**).

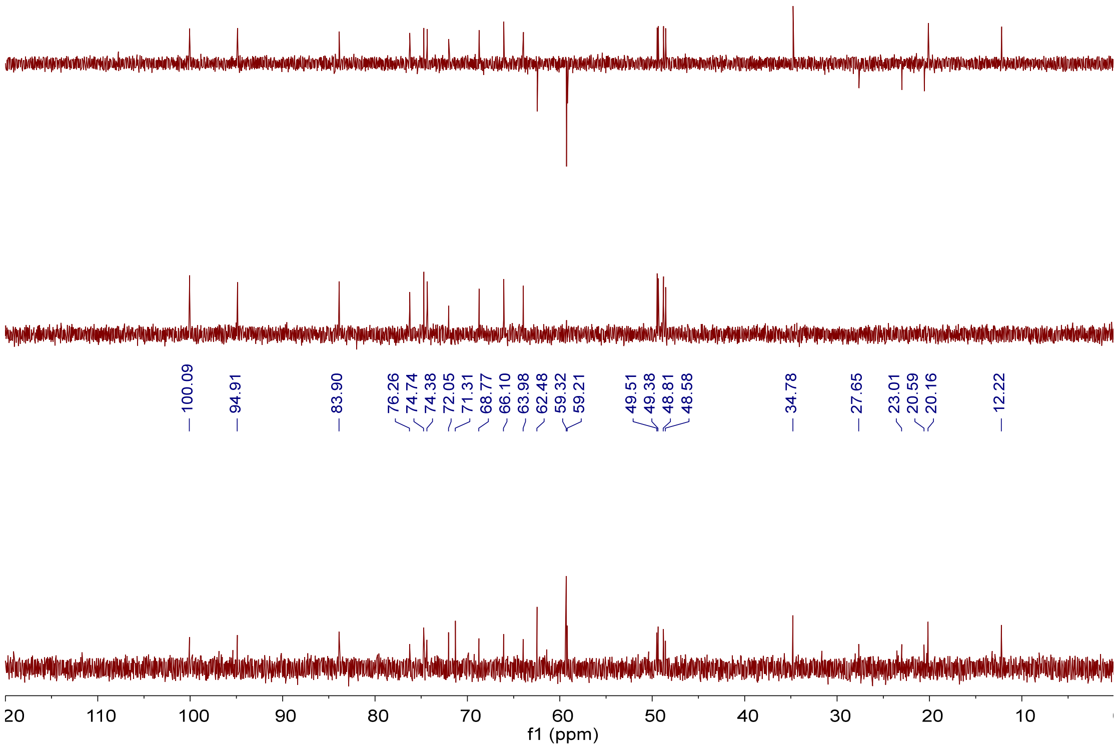

**Supplementary Fig. 42** $^{13}C$ NMR, DEPT-90 and DEPT-135 spectra (150 MHz, $D_2O$) of GK-C2 (**9**).

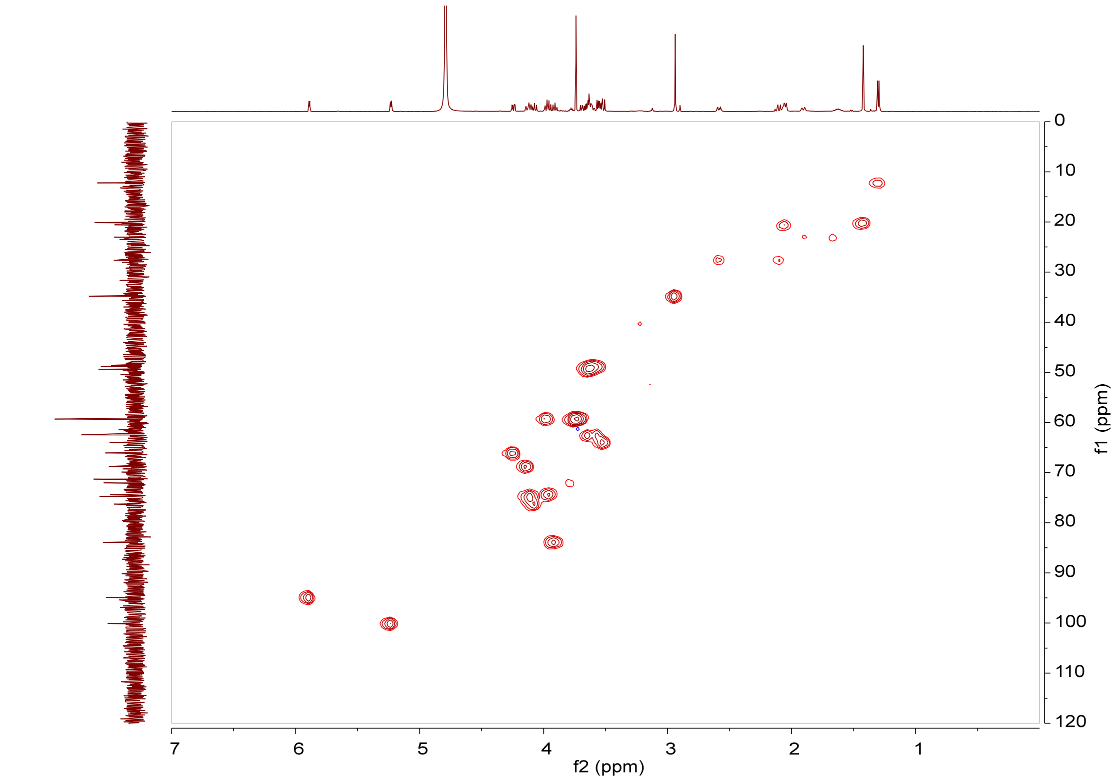

**Supplementary Fig. 43** HSQC spectrum (600 MHz, $D_2O$) of GK-C2 (**9**).

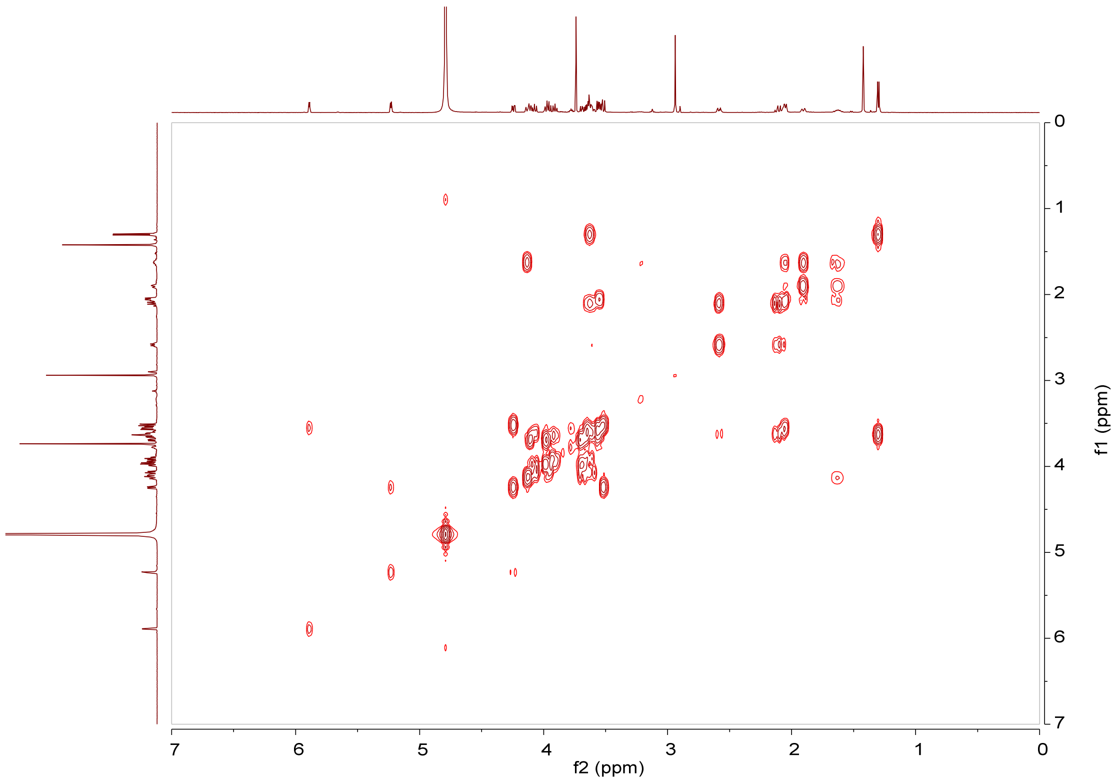

**Supplementary Fig. 44** $^1H$–$^1H$ COSY spectrum (600 MHz, $D_2O$) of GK-C2 (**9**).

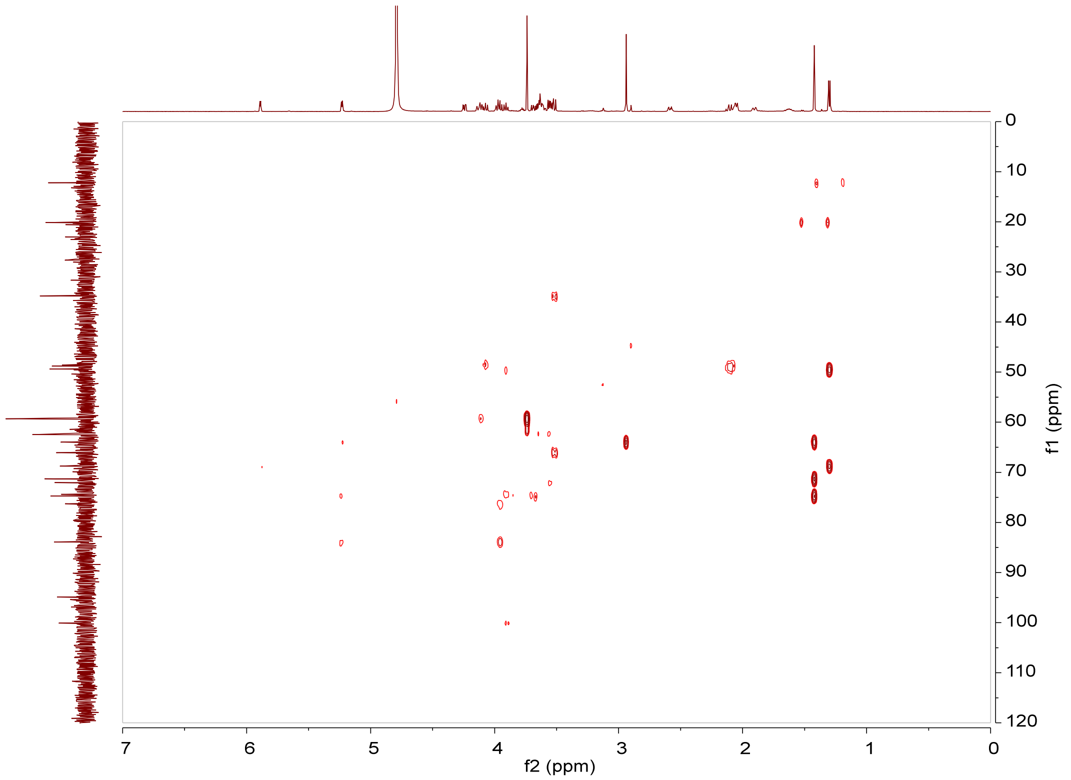

**Supplementary Fig. 45** HMBC spectrum (600 MHz, $D_2O$) of GK-C2 (**9**).

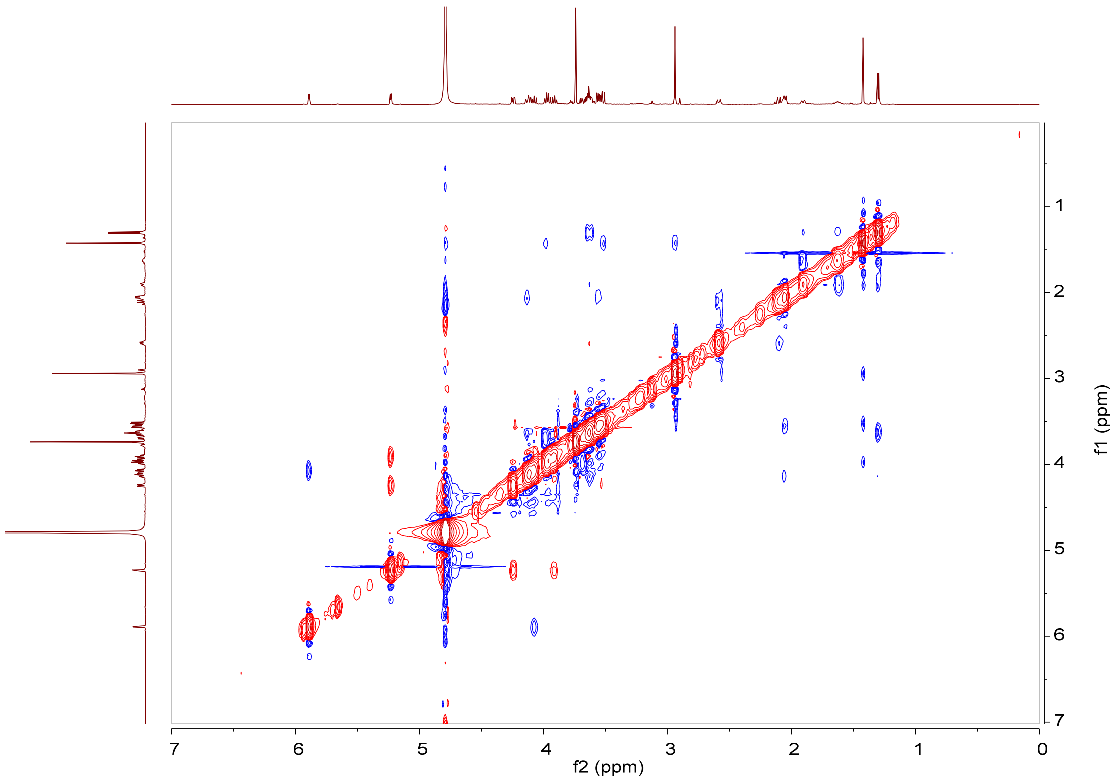

**Supplementary Fig. 46** NOESY spectrum (600 MHz, $D_2O$) of GK-C2 (**9**).

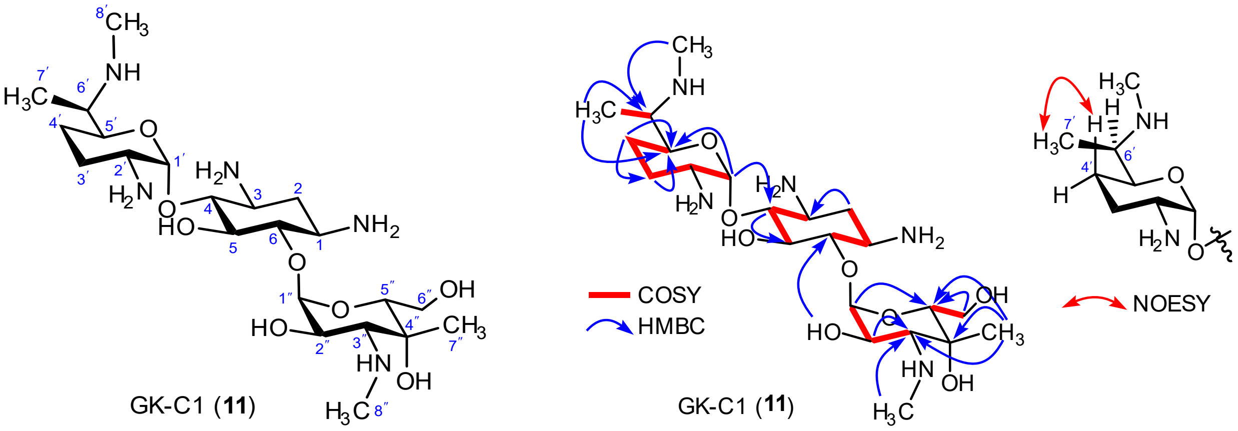

**Supplementary Fig. 47** Chemical structure (left), key $^{1}H$–$^{1}H$ COSY, HMBC and NOESY correlations of GK-C1 (**11**) (right).

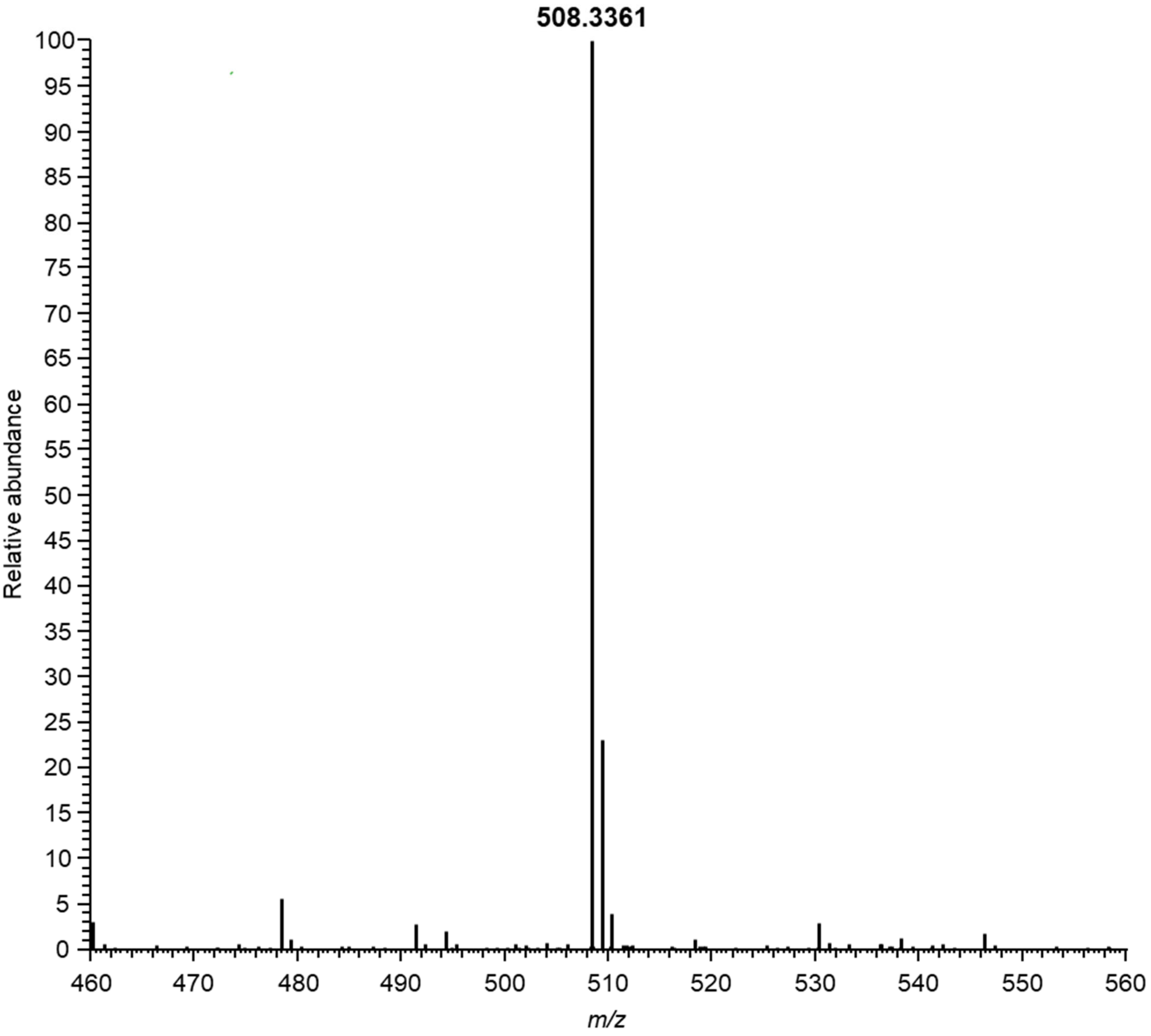

**Supplementary Fig. 48** ESI-HR MS spectrum of GK-C1 (**11**).

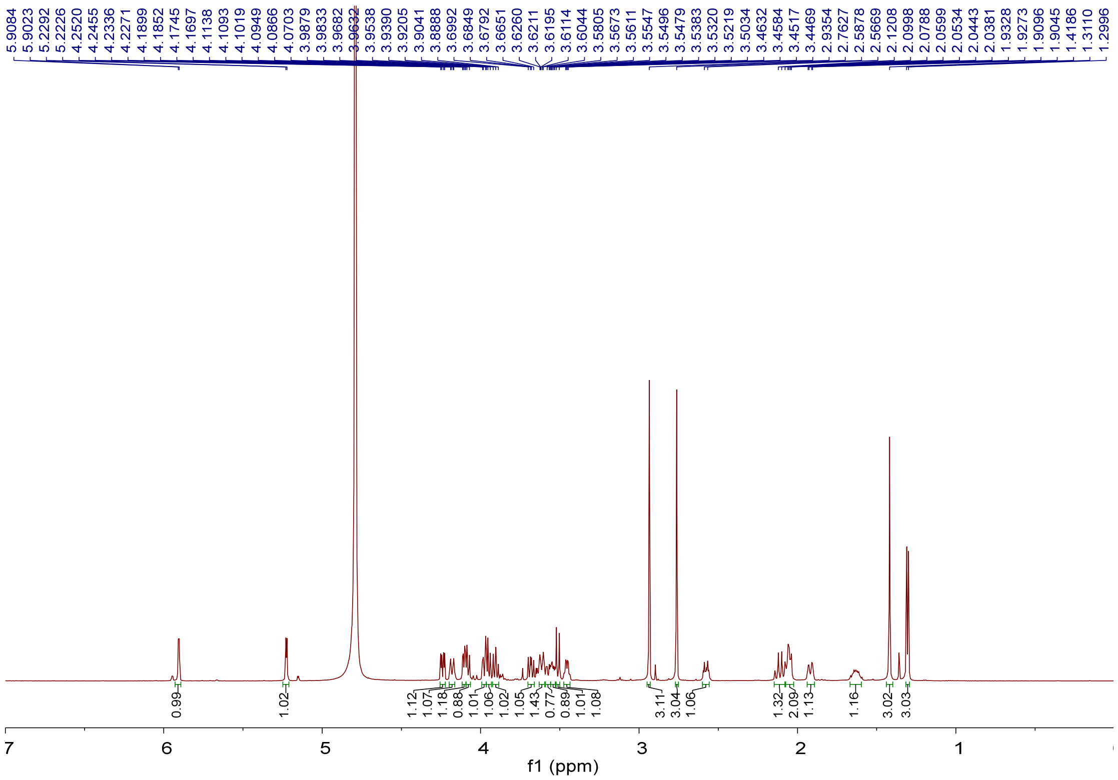

**Supplementary Fig. 49** $^{1}H$ NMR spectrum (600 MHz, $D_2O$) of GK-C1 (**11**).

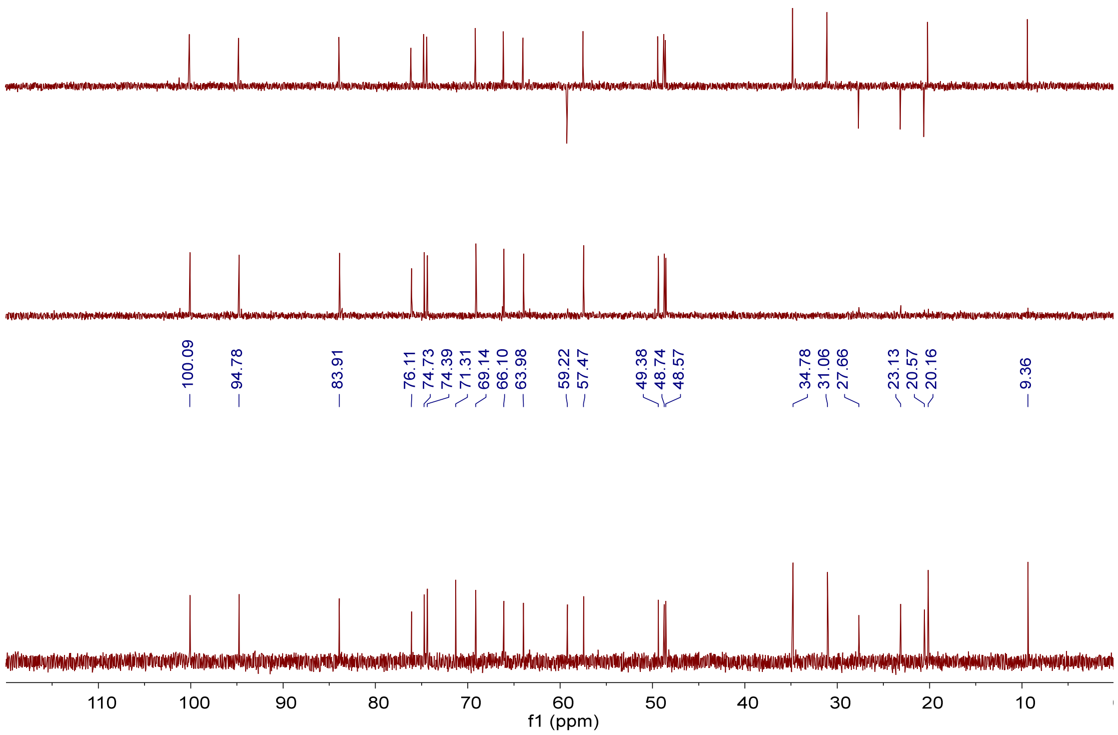

**Supplementary Fig. 50** $^{13}C$ NMR, DEPT-90 and DEPT-135 spectra (150 MHz, $D_2O$) of GK-C1 (**11**).

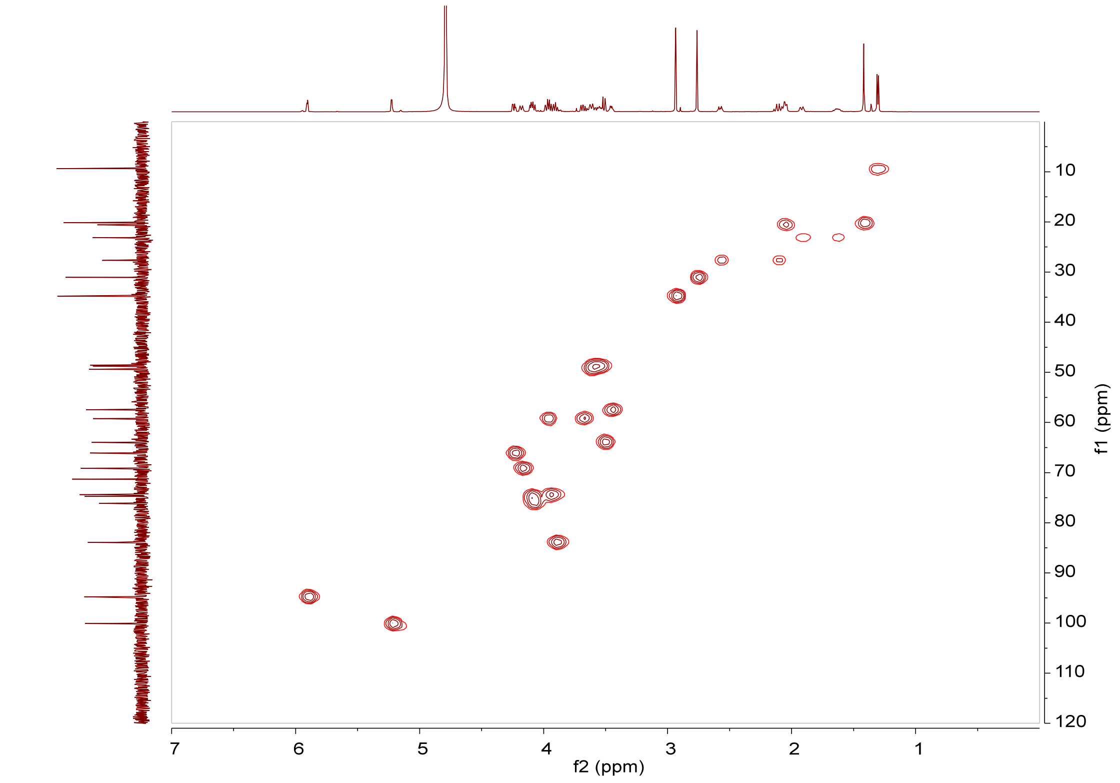

**Supplementary Fig. 51** HSQC spectrum (600 MHz, $D_2O$) of GK-C1 (**11**).

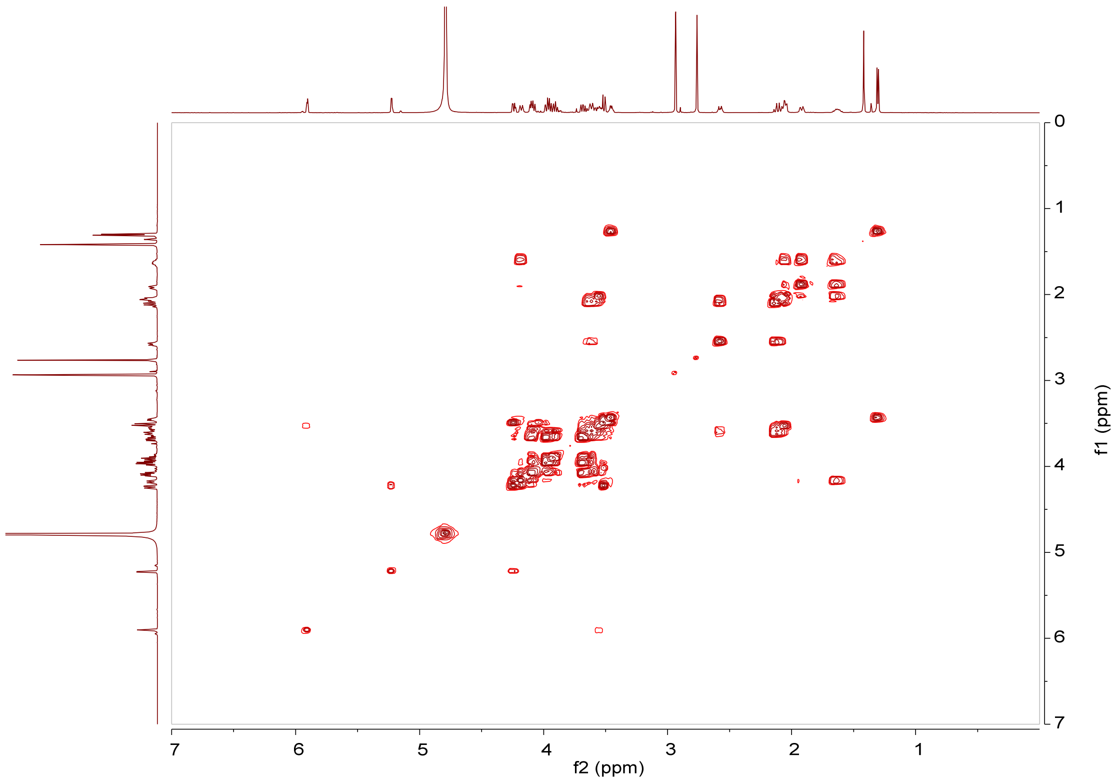

**Supplementary Fig. 52** $^1H$–$^1H$ COSY spectrum (600 MHz, $D_2O$) of GK-C1 (**11**).

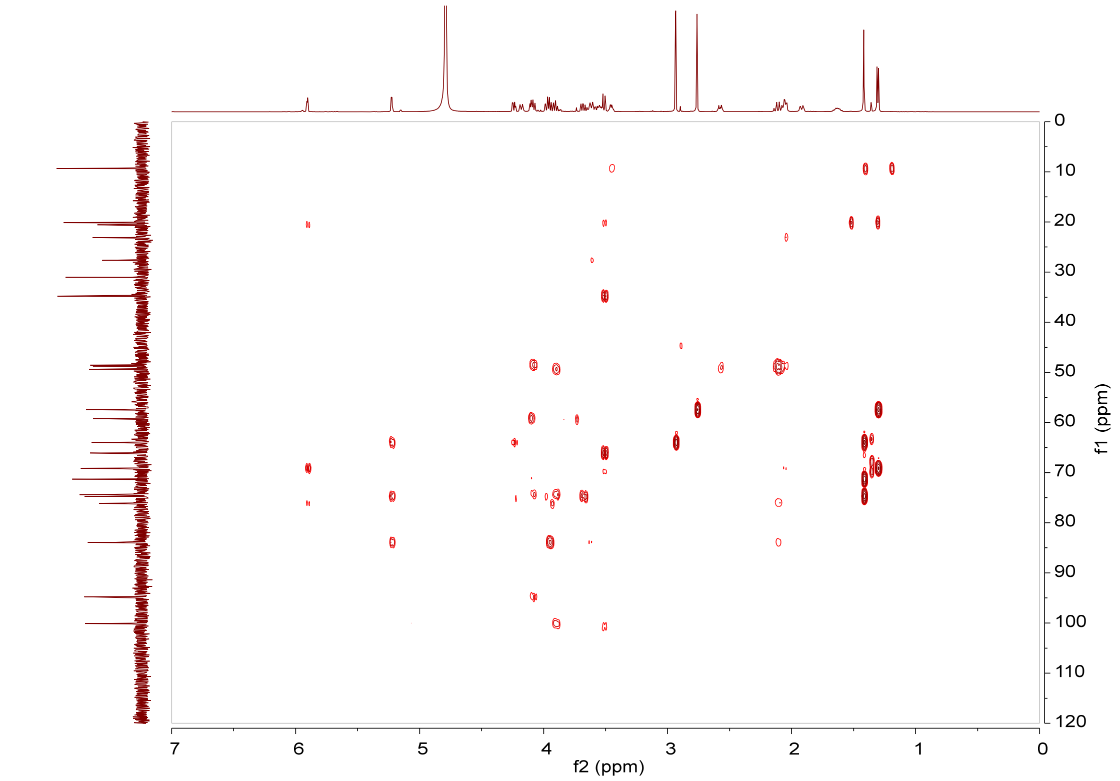

**Supplementary Fig. 53** HMBC spectrum (600 MHz, $D_2O$) of GK-C1 (**11**).

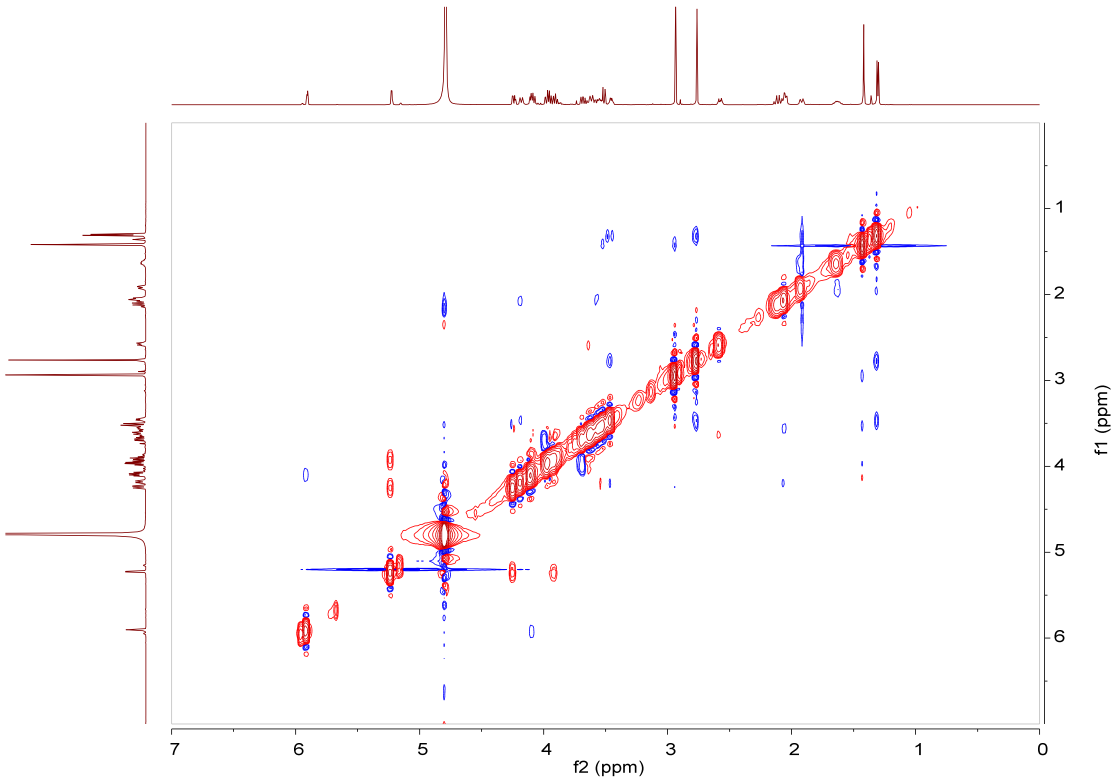

**Supplementary Fig. 54** NOESY spectrum (600 MHz, $D_2O$) of GK-C1 (**11**).

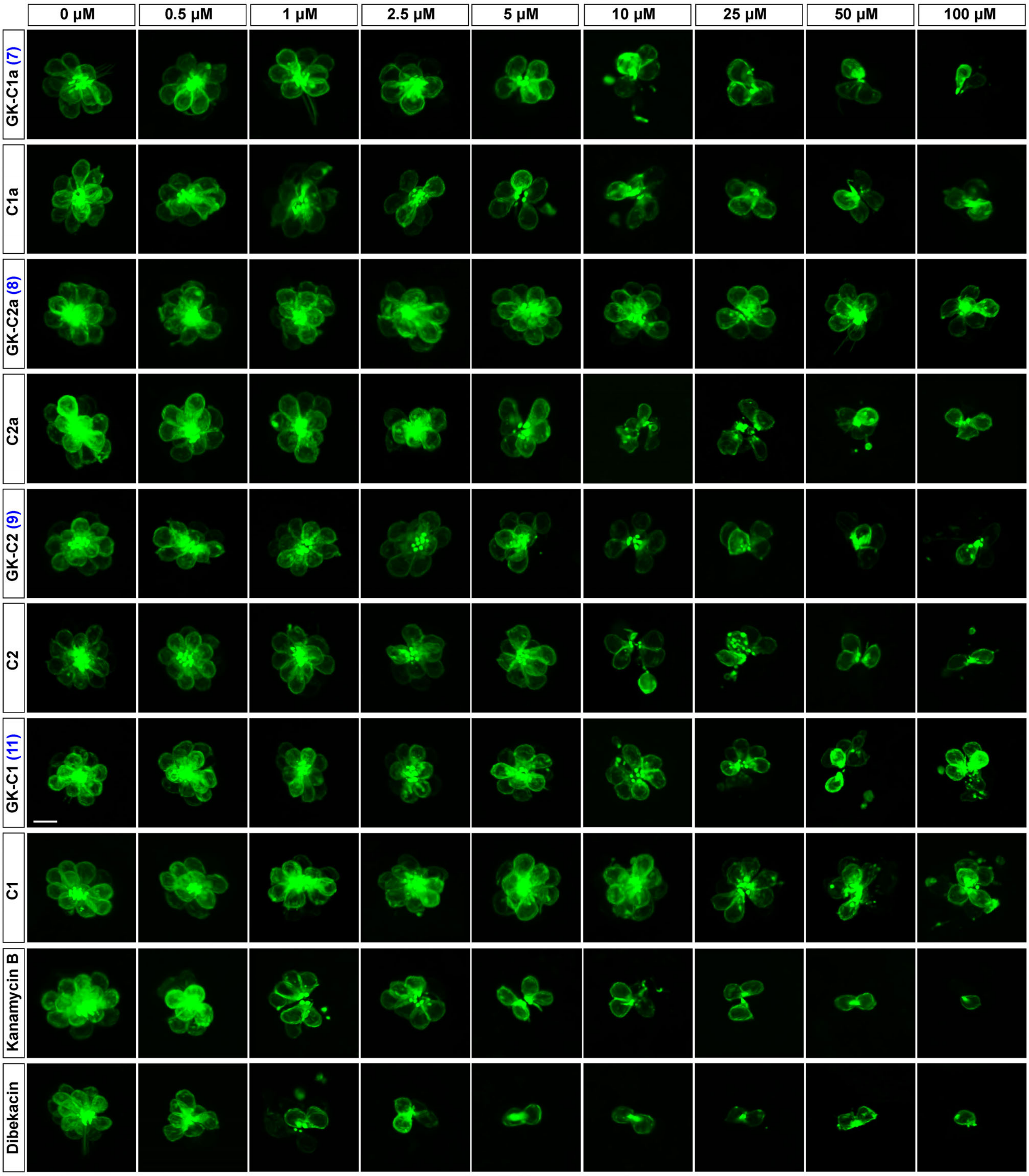

**Supplementary Fig. 55** Representative fluorescent images of HC cluster in L1 neuromast of pLL under different compounds treatment at concentrations of 0.5, 1, 2.5, 5, 10, 25, 50, 100 μM. Scale bar: 10 μm.

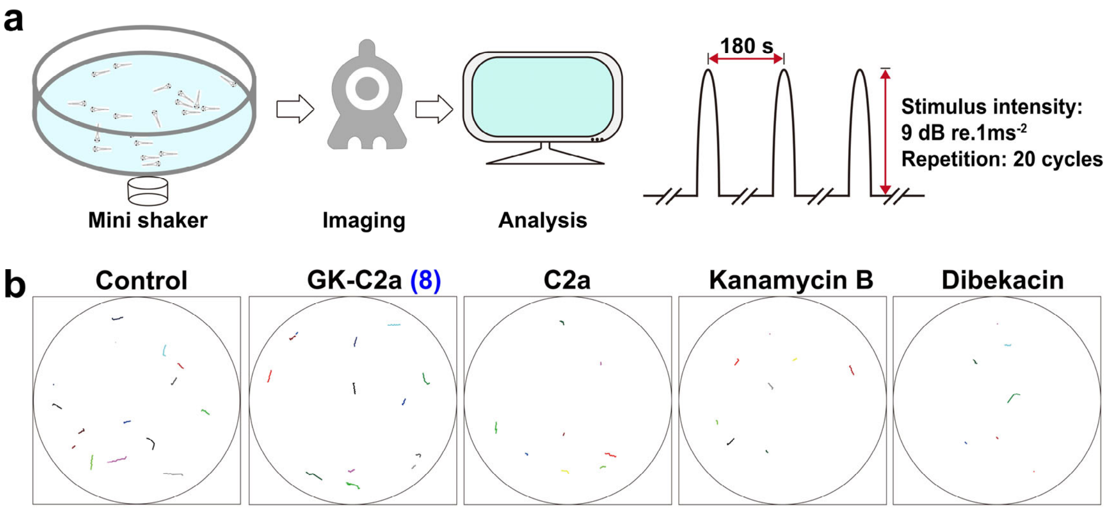

**Supplementary Fig. 56** The startle response assays of zebrafish larvae (5 dpf) under treatment with four compounds of GK-C2a (**8**), C2a, kanamycin B and dibekacin. (**a**) Schematic diagram of the startle response testing equipment and experimental parameters. (**b**) Extracted locomotion trajectories from the response behaviors under a one-time stimulus of 9 dB re.1 $ms^{-2}$ under 10 μM different compounds treatment.
